# Adjacent Rare-Earth Separation via a Protein Atomic Ruler

**DOI:** 10.1101/2025.10.21.683075

**Authors:** Xining Qian, Yunxiang Du, Chao Ma, Huijing Cui, Huaxia Zhu, Tongjia Shi, Yang Bai, Fangkun Wang, Zhengqing Li, Jing Yu, Huajian Gao, Hongjie Zhang, Lei Liu, Kai Liu

## Abstract

The separation of rare-earth elements (REEs) poses a formidable challenge due to their nearly identical ionic radii and chemical properties. Here, we present a transformative strategy for REE separation using a **m**ultimetal **i**on-stacking metalloprotein **f**ramework (MIF) that functions as an atomic ruler, effectively amplifying subtle differences in REE ionic radii. This approach achieves exceptional selectivity, enabling the separation of all 13 adjacent REE pairs with ultrahigh purity (up to 99.999%). Remarkably, the MIF retains its lanthanide-binding capacity over 410 consecutive absorption-desorption cycles, demonstrating outstanding stability. Structural insights reveal that the MIF’s densely packed REE coordination environment magnifies minute radius variations—a key mechanism underlying its unparalleled separation efficiency. Our MIF-based strategy offers a sustainable, scalable solution to rare-earth separation in industrial applications.

## Introduction

High-purity rare earth elements (REEs) are indispensable in modern technology, including electronics, photonics, and magnetic clean-energy devices (*1, 2*). A great challenge in the rare earth industry is the separation of chemically similar lanthanides (*3, 4*), whose nearly identical outer electron configurations, with a mere 0.2 Å difference in ionic radii from La^III^ to Lu^III^ (*5, 6*), render them notoriously difficult to isolate (*7*). Current hydrometallurgical separation processes rely heavily on liquid□liquid extraction technologies that employ organic solvents and corrosive phosphoric acid-based extractants (*8, 9*). These processes require hundreds to thousands of extraction stages to isolate adjacent REEs (*10, 11*), leading to substantial energy consumption and environmental pollution (*9, 12*). Aqueous strategies that use synthetic chelators (*13-20*) and proteinaceous ligands (*21-30*) have been explored as alternatives to solvent extraction for REE separation. However, most ligands have a metal-to-ligand ratio (R_m/l_) ≤ 1, indicating low binding capacity (*18, 27, 29, 31, 32*). Moreover, the coordination microenvironments of existing ligands fail to resolve the picometer-scale differences in ionic radii among REEs, making them ineffective for separating adjacent REEs (*24, 33-35*). These critical challenges underscore the urgent need for novel strategies that exploit picometer-scale disparities in ionic radii to achieve rare-earth isolation (*7*).

Herein, we report a groundbreaking strategy for REE separation using a unique, rigid multimetal ion-stacking metalloprotein framework (MIF, Fig. 1A). The MIF simultaneously coordinates 5-6 REE ions (R_m/l_ ≥ 5), enabling unprecedented all-aqueous, single-stage, full-spectrum separation of REEs. It outperforms other proteinaceous and traditional ligands (*24, 29, 34*), particularly in terms of separation factors for adjacent REEs (*35*). The MIF features a β-barrel structure with densely stacked coordination sites that accumulate disparities in ionic radii. Acting as an atomic ruler, it restricts residue displacement and amplifies picometer-scale differences into global structural deformation and binding energy gradients, thereby enabling adjacent separation of all REE pairs (excluding radioactive Pm).

**Fig. 1.**
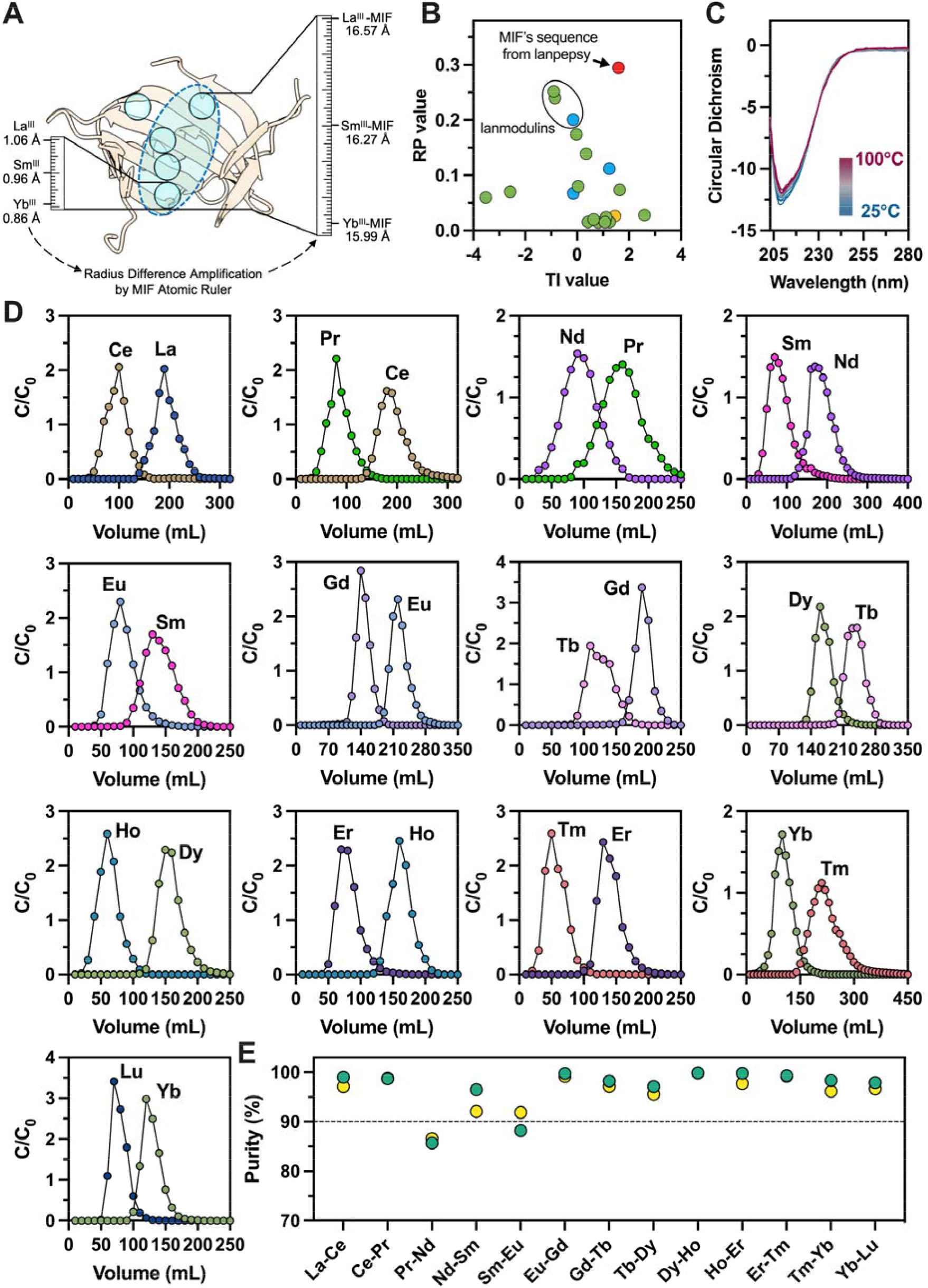
Identification of the MIF and its performance in separating adjacent lanthanides. **(A)** The MIF acts as an atomic ruler to discriminate REEs with very small ionic radius differences. **(B)** Identification of the MIF sequence using two metrics. Each circle represents a lanthanide-binding protein, plotted on the basis of its melting-temperature index (TI, a sequence-based predictor of thermostability) and REE payload (RP, lanthanide ions bound per□kilodalton). Red circle: the MIF’s sequence from lanpepsy. Green circles: natural lanthanide-binding sequences. Blue circles: engineered peptides. Yellow circles: *de*□*novo*-designed REE-binding proteins. The red circle combines the highest RP with a high TI to identify sequences suitable for MIF construction. **(C)** The REE-bound MIF maintains a regular structure at temperatures up to even 100□°C, as evidenced by CD spectra (with La^III^-MIF shown as an example). **(D)** Single-stage separation of 13 adjacent REE pairs by the MIF under aqueous conditions. **(E)** Purity of REEs achieved in the adjacent REE pairs after single-stage MIF separation. The yellow and green circles denote the first and second elements of adjacent REE pairs, respectively.

## Result and discussion

### Finding an atomic ruler protein for adjacent-REE separation

Our research commenced with the identification of a MIF with high stability and superior lanthanide-loading capacity. All 20 reported lanthanide-binding proteins with experimental data (see table S1) were evaluated using two metrics: (i) the melting-temperature index (TI) predicted from the protein sequence (*36*), which is positively correlated with the melting temperature (T_m_), and (ii) the REE payload (RP), defined as the number of lanthanide ions bound per kilodalton of protein. On the basis of this investigation, the sequence of lanpepsy from *Methylobacillus flagellates* (*26*) met the criteria for an MIF (Fig. 1B). It presented the highest REE payload in the cohort (RP = 0.295 Ln/kDa) and high stability, with a predicted T_m_ > 65°C (TI = 1.59). Our experiments revealed that the His_10_-tagged MIF could be expressed in *Escherichia coli*, reaching a yield of 1.16 g/L, suggesting its potential suitability for industrial-scale production (fig. S1). Variable-temperature circular dichroism (VT-CD) spectroscopy further confirmed the exceptional stability of the MIF: the protein retained more than 95% of its native secondary structure after incubation at 100°C for 30 min and underwent almost complete refolding upon cooling (Fig. 1C, fig. S2).

Next, we evaluated the performance of the MIF in separating adjacent lanthanide pairs. The protein was immobilized onto agarose beads, which were then packed into columns, and each pair of adjacent lanthanides (13 nonradioactive pairs in total, namely, La^III^-Ce^III^, Ce^III^-Pr^III^, Pr^III^-Nd^III^, Nd^III^-Sm^III^, Sm^III^-Eu^III^, Eu^III^-Gd^III^, Gd^III^-Tb^III^, Tb^III^-Dy^III^, Dy^III^-Ho^III^, Ho^III^-Er^III^, Er^III^-Tm^III^, Tm^III^-Yb^III^, and Yb^III^-Lu^III^) was tested sequentially at equimolar concentrations. Elution was performed using a citrate gradient (*22, 28*), and the effluent was collected in equal-volume fractions, with the REE content analyzed via inductively coupled plasma□optical emission spectrometry (ICP□OES). All 13 adjacent lanthanide pairs were found to elute as distinct peaks, with heavier REEs consistently eluting ahead of their lighter counterparts (Fig.□1D). For each pair, fractions containing separated REEs were collected, and the purity and yield of each REE were calculated **(*23*)**. Eleven of the 13 pairs exhibited excellent separation of individual REEs, with purities ranging from 92.1% to 99.8% and yields ranging from 90.6% to 99.6% (table S2). The Pr^III^-Nd^III^ and Sm^III^-Eu^III^ pairs exhibited purities of 86.6% (Pr^III^)/85.7% (Nd^III^) and 91.9% (Sm^III^)/88.2% (Eu^III^), respectively, which are sufficient for industrial applications (*37*) (*38, 39*). These observations demonstrate that our approach enables single-stage, full-spectrum REE separation. This performance has never been achieved by conventional chemical approaches using selective ligands such as alkyl-phosphorus extractants (*40, 41*) and organic multidentate ligands (*42-44*). Furthermore, we performed a control experiment using the recently reported lanmodulin ligands (*23-25*). Three adjacent pairs, La^III^-Ce^III^ (light REEs), Eu^III^-Gd^III^ (middle REEs), and Ho^III^-Er^III^ (heavy REEs), were evaluated. The results showed lanmodulin cannot discriminate adjacent lanthanide pairs, exhibiting only limited purities (59.0-68.7%) and yields (46.9-76.2%) (fig. S3). Overall, the MIF provides the first ligand capable of single-stage, aqueous-phase separation of all adjacent lanthanide pairs.

### Crystal structure analyses reveal a multi-metal ion-stacking metalloprotein framework

Next, we determined the X-ray crystal structures of the ion-free (apo) form of the MIF at a resolution of 2.08 Å (fig. S4) and the La^III^-bound form at 1.25 Å (Fig. 2A). Both structures exhibit a compact, channel-like architecture consisting of two PepSY domains (*45*) arranged into a pseudo-C_2_ symmetric central electronegative β-barrel (fig. S4). The eight antiparallel β-strands are sequentially designated β1 to β8 from the N-terminus to the C-terminus (**f**ig. S5). Alignment of the Cα backbones of the apo- and La^III^-bound structures using PyMOL (*46*) revealed highly similar backbone conformations (fig. S6), confirming the high structural rigidity of the protein framework. The La^III^-bound structure reveals five lanthanide-binding sites arranged in an atypical Y-shaped configuration along the β-barrel axis, forming a rigid multimetal-coordinated framework, an unprecedented MIF architecture among all metalloproteins (Fig. 2B, fig. S7). Sites 1-3 in this MIF are densely stacked within the acidic β-barrel channel and slightly tilted toward the dimerization interface of the PepSY β4-β5 strands. These three La^III^ ions exhibit dense stacking with an average interionic distance of 4.3 Å (Fig. 2C), even approaching the La^III^-La^III^ distance in the densely packed lattice of inorganic LaPO_4_ crystals (*47*). Both sites 4 and 5 are located at the exit of the acidic channel: site 4 is proximal to the PepSY β4-β5 dimerization interface, and site 5 is near the β1-β8 dimerization interface. With respect to the coordination microenvironment of the MIF, 19 residues (15 glutamate (E), 3 aspartate (D), and 1 glutamine (Q)) are utilized to coordinate La^III^ ions (fig. S5). Notably, residues D54, E68, E106, E108, E118, and E132 act as bridging ligands to coordinate neighboring La^III^ ions, stabilizing this unique, densely stacked MIF architecture (Fig. 2C, fig. S8). Additional stabilization is provided by ε-amino groups from K29, K33, and K130, as well as water molecules, which form hydrogen bonds with multiple acidic E and D residues (fig. S5). Thus, the distance between REE sites in the MIF is densely stacked at only 4.3 Å. In stark contrast, another lanthanide-binding protein, lanmodulin (PDB ID: 8DQ2), contains multiple REE-binding sites with distances ranging from 12 to 30 Å and forms dispersed coordination spheres without any bridging interactions (fig. S9). Collectively, the structures of both apo- and La^III^-bound MIF reveal a unique lanthanide coordination environment that differs from that of all known lanthanide-binding proteins reported to date (*21, 24, 25, 29, 48, 49*).

**Fig. 2.**
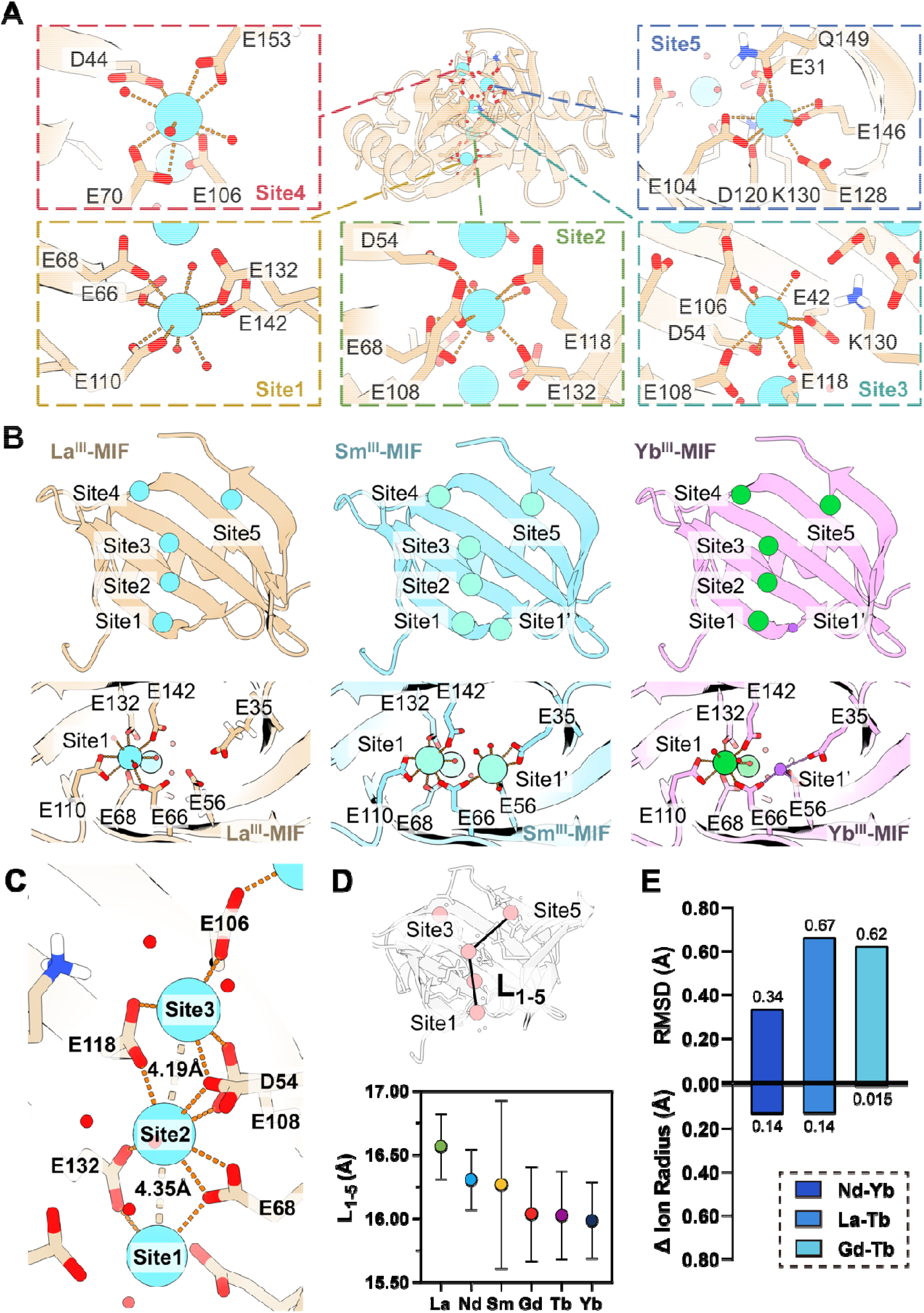
Structural basis for adjacent REE separation by the MIF. **(A)** Crystal structure of the La^III^-MIF complex, featuring five densely stacked REE-binding sites. Carboxylate and amide O atoms (red) from Glu, Asp, and Gln coordinate with La^III^ ions (cyan spheres). **(B)** Structural details of the La^III^-, Sm^III^-, and Yb^III^-MIF complexes. The top panel shows the side view of the MIF. Coordination stoichiometries vary with different REEs: light (La^III^) and heavy (Yb^III^) REEs bind five sites, whereas middle REEs (Sm^III^) occupy six sites. The bottom panels show the top view of the MIFs. Notably, the middle REE-MIF complex has an additional coordination site, Site 1’, which is absent in the light REE-MIF complex. In the heavy REE-MIF complex, Site 1’ is instead occupied by a Na^I^ ion (purple sphere). **(C)** La^III^ ions at Sites 1-3 (cyan spheres) are densely stacked within the coordination microenvironment of the MIF, with an interionic distance of approximately 4.3 Å. Orange dashed lines depict interactions mediated by residues D54, E68, E106, E108, E118, and E132, which act as bridging ligands to coordinate neighboring La^III^ ions and stabilize the densely stacked MIF architecture. **(D)** L_1-5_ is defined as the sum of the distance from Site1 to Site3, and the distance from Site3 to Site5 (top panel). For the six REE–MIF complexes (bottom panel), the L_1-5_ values range from 16.57 to 15.99 Å, in contrast to the variation in the corresponding REE ionic radii (ranging from 1.06 to 0.86 Å). This finding indicates that the densely stacked architecture of the MIF amplifies picometer-scale differences in REE ionic radii. The error bars represent the root-mean-square error of each distance, which was calculated from the B-factor-derived mean-square displacement of each ion of interest in the crystal structures. **(E)** Upper bars show pairwise Cα root-mean-square deviation (RMSD) values for the three REE-MIF pairs (Nd-Yb, La-Tb, and Gd-Tb), ranging from 0.34 Å to 0.67□Å. Lower bars show the corresponding differences in ionic radius (abbreviated as Δ Ion Radius), ranging from 0.015 Å to 0.14 Å. The RMSD values of the REE-MIF backbones are markedly larger than the differences in REE ion radii, indicating that the MIF amplifies the picometer-scale differences in REE ionic radii to the angstrom level.

We solved more REE-MIF structures, including those bound to Nd^III^ (1.41 Å), Sm^III^ (2.16 Å), Gd^III^ (1.41 Å), Tb^III^ (1.58 Å), Ho^III^ (3.05 Å), and Yb^III^ (1.21 Å) (figs. S10-15). Global superimposition of these structures against each other revealed high overall structural conservation (fig. S6), wherein the densely Y-shaped stacked binding sites 1 through 5, a key structural feature of MIF, were consistently preserved across all the REE complexes. Nevertheless, we observed differences in the metal binding stoichiometries among these complexes (Fig. 2B). REE-MIFs with medium ionic radii (i.e., Sm^III^, Gd^III^, Tb^III^, and Ho^III^) exhibited an additional metal ion binding site, designated Site 1’, which is located beside Site 1 (fig. S16). Site 1’ was also observed in the structure of the MIF complex with Yb^III^ (a heavier REE with a smaller ionic radius); however, the site does not accommodate an anomalous difference in density consistent with a Yb^III^ ion and was thus modeled with Na^I^ (fig. S15). Furthermore, we observed differences in the coordination patterns among the REE-MIFs. For example, residues E68 (bridging between Sites 1 and 2) and E104 (at Site 5), which exhibit bidentate coordination with La^III^, shift to monodentate coordination in the Yb^III^ complex (fig. S17). Thus, REE ions with different ionic radii are associated with distinct metal-binding stoichiometries and coordination patterns within REE-MIFs.

Next, we quantitatively analyzed the coordination microenvironment of the MIF by measuring the length of the MIF channel (L_1-5_) as the sum of the distances between Site 1 and Site 3 and between Site 3 and Site 5 (Fig. 2D). We observed that the L_1-5_ values differed among the REEs in the MIF, with the changes in the L_1-5_ values significantly exceeding the changes in the corresponding REE ionic radii. For example, from the La^III^ (radius = 1.06 Å) to Yb^III^ (radius = 0.86 Å), although the radius changed only by only 0.2 Å, the L_1-5_ value changed by 0.58 Å (from 16.57 Å to 15.99 Å), representing a 3-fold amplification of the radius change. Moreover, despite a mere 0.026 Å difference in radius between Sm^III^ (radius = 0.96 Å) and Gd^III^ (radius = 0.94 Å), the L_1-5_ values between these two REEs differed by 0.23 Å, corresponding to a 9-fold amplification of the picometer-scale radius change. We also aligned all the REE-MIF structures (fig. S6) and calculated the pairwise RMSD values for all the Cα positions of the superimposed structures (Fig. 2E). The smallest RMSD difference (0.34 Å) was observed between the Nd^III^-bound (radius = 1.00 Å) and Yb^III^-bound (radius = 0.86 Å) structures, corresponding to a 2.5-fold amplification of the ionic radius change (**f**ig. S6). The largest RMSD change (0.67 Å) occurred between the La^III^-bound (radius = 1.06 Å) and Tb^III^-bound (radius = 0.92 Å) structures, representing a 4.9-fold amplification (**f**ig. S6). More strikingly, despite a mere 0.015 Å difference in ionic radius between Gd^III^ (radius = 0.94 Å) and Tb^III^ (radius = 0.92 Å), their structural RMSD difference reached 0.62 Å, showing a remarkable 42-fold amplification (fig. S6). The above results demonstrate that binding REEs with subtle ionic radius differences induces amplified changes in the MIF channel length and backbone positions. This establishes the MIF as an atomic ruler capable of picometer-scale discrimination of REEs on the basis of ionic radius.

### Energetic features of MIF’s selective binding with different REEs

To assess differences in binding affinity among REEs, we first measured the separation factors of the MIF for 13 adjacent lanthanide pairs. Free and protein-bound REE concentrations were quantified via ultrafiltration and ICP□OES, enabling the calculation of distribution values (D) (fig. S18). The results show that Sm^III^ has the highest binding affinity among all lanthanides to the MIF, with the affinity decreasing monotonically toward both lighter and heavier lanthanides. To quantify this selectivity behavior, we performed isothermal titration calorimetry (ITC) and measured the binding-associated heat and dissociation constants (*K*_d_) between the MIF and 14 nonradioactive lanthanide ions (fig. S19). The apparent binding free energy (ΔG_app_) is consistent with the binding affinity trend, reaching a minimum for Sm^III^ (ΔG_app_ = -45.6 ± 0.4 kJ/mol), compared with La^III^ (-42.0 ± 0.1 kJ/mol) and Tm^III^ (-38.1 ± 0.2 kJ/mol), which are representative of lighter and heavier lanthanides, respectively (Fig. 3A). The ITC results showed that the binding of each REE to the MIF is endothermic, and the accompanying entropic gain outweighs the enthalpic penalty, resulting in an overall favorable negative Gibbs free energy. Sequential binding model analysis of the *K*_d_ values for the first five REE binding sites in the MIF revealed four high-affinity sites (*K*_d1-4_), with *K*_d_ values ranging from 23.9 pM to 3.69 µM, and one low-affinity site with a *K*_d_ > 10 µM (Fig. 3B). The ratio of maximum to minimum *K*_d_ values spans 5-fold (for *K*_d2_) to 15,000-fold (for *K*_d4_), leading to a 110-fold difference between the maximum global apparent *K*_d_ (Lu^III^-MIF) and the minimum (Sm^III^-MIF), with adjacent lanthanide pairs differing by an average of 1.7-fold (fig. S18). These results are consistent with the REE selectivity of the MIF observed in the distribution value measurements.

**Fig. 3.**
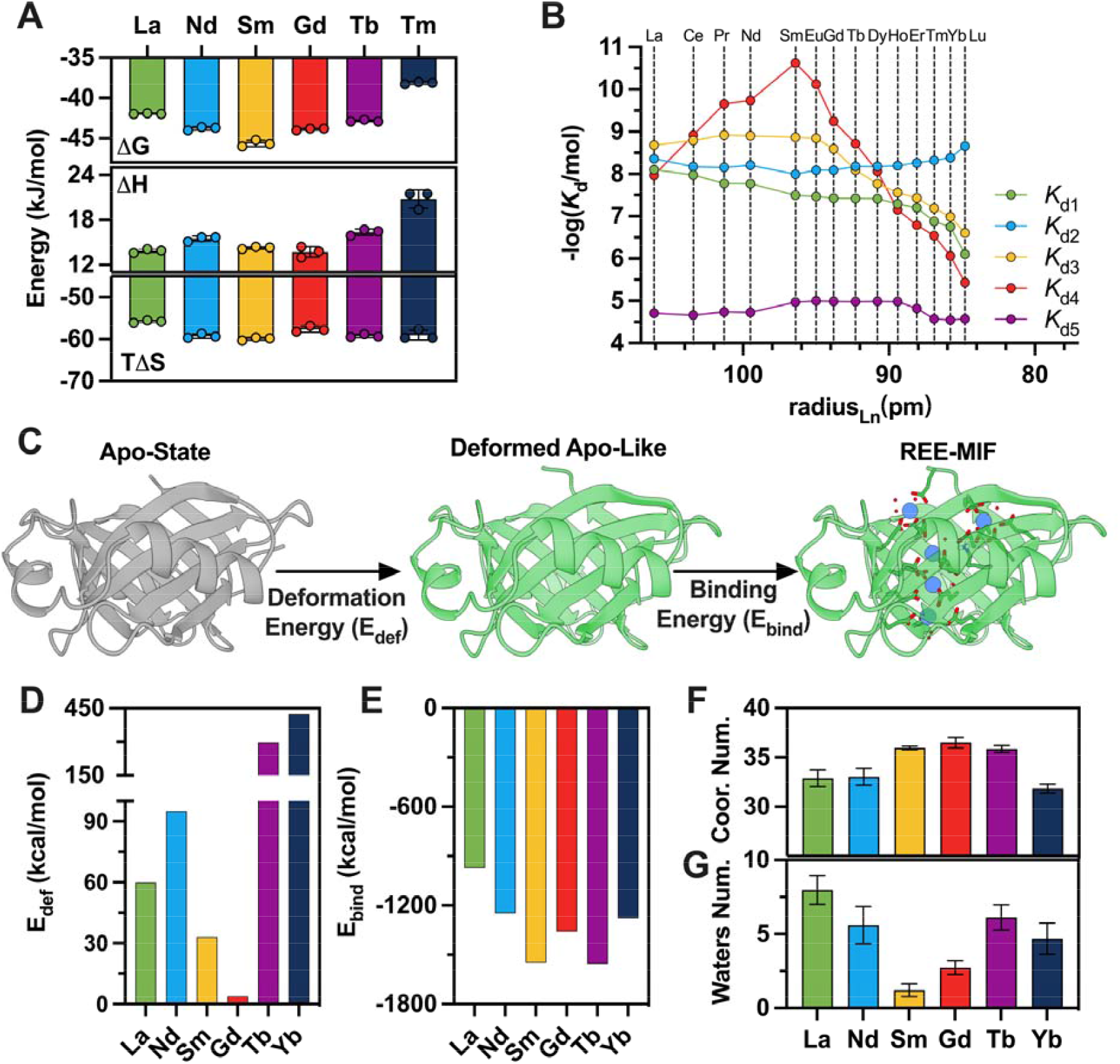
Energetic basis of the selectivity of MIF for different REEs. **(A)** Thermodynamic parameters (ΔG, ΔH, and -TΔS) for MIF binding to REEs (La^III^, Nd^III^, Sm^III^, Gd^III^, Tb^III^, and Tm^III^) as determined by ITC. Each bar represents the mean value from three independent replicates (n = 3). All the binding processes are endothermic (ΔH > 0), indicating that the REE□MIF interactions are predominantly entropy driven. **(B)** Dissociation constants (*K*_d_) for binding of the MIF with different REEs, as determined by ITC. The results revealed a selectivity trend toward middle REEs. **(C)** Two-step energetic calculations for REE binding to the MIF. **(D)** MIF deformation energy (E_def_) after binding to different REEs. **(E)** MIF’s binding energy (E_bind_) to different REEs. Both the E_def_ and E_bind_ reach their minima for middle REE-MIF complexes and increase progressively for light and heavy REE-MIF complexes. **(F)** Total coordination number (Coor. Num.) for the MIF with different REEs, as calculated from MD simulations, revealing a maximum Coor. Num. for middle REE-MIF complexes. The error bars indicate fluctuations in Coor. Num. observed throughout the MD trajectory over 2700-3000 ns. **(G)** Number of water molecules (Water Num.) within the 6-Å sphere centered at Site 2 of the MIF when bound to different REEs. Consistent with the MIF having the highest affinity for middle REEs, these complexes have the lowest number of water molecules, indicating a strong desolvation effect. The error bars indicate fluctuations in Water Num. observed throughout the MD trajectory over 2700-3000 ns.

The aforementioned ITC experiments demonstrate that the selectivity of the MIF originates from its stronger binding affinity for middle REEs than for light and heavy REEs. To gain deeper mechanistic insights into this selectivity, we calculated the binding energy of REE-MIF complexes using the structures of apo-MIF and REE-MIFs. Owing to the large size of the MIF protein (over 2000 atoms), which exceeds the computational limits of density functional theory, we employed a two-step energy calculation approach to determine the deformation energy (E_def_) and REE-MIF binding energy (E_bind_) (Figs. 3C-E, fig. S20, and tables S4, S5). The results reveal that the REE-MIF complexes of the middle REEs present the lowest E_def_ and E_bind_ values, with both energies increasing toward lighter and heavier lanthanides. This energetic trend aligns with the binding affinity profiles determined by ITC experiments. The large difference in E_bind_ between light and middle REEs indicates that binding energy is the primary driver of selectivity for these pairs. In contrast, for middle and heavy REEs, the difference in E_def_ is more pronounced than that between light and middle REEs, indicating that deformation energy is the main contributor to selectivity in these pairs.

To further characterize the dynamics of the coordination microenvironment, we conducted μs-scale all-atom MD simulations on each REE-MIF complex (fig. S21). The time-averaged coordination number statistics reveal the highest coordination (n = 35-37) for middle REEs, whereas light and heavy REEs have coordination numbers ≤ 33.4 (Fig. 3F and fig. S22). Desolvation was assessed by counting water molecules within a 6 Å sphere centered on Site 2—the site closest to the MIF’s mass center (fig. S23). Consistent with the MIF’s strong binding to middle REEs, these complexes contained few water molecules, indicating a strong desolvation effect (Fig. 3G and fig. S18). Collectively, the above results establish the structure□energy relationship of REE□MIF complexes.

### Single-stage high-purity REE separation with an MIF atomic ruler

To evaluate the industrial applicability of the MIF, we packed agarose beads with covalently immobilized MIF into a chromatography column integrated with an automated gradient elution system and performed breakthrough analysis (*23*). La^III^ was selected as a model ion, and a La^III^ solution was used for breakthrough analysis. The results showed that the functionalized beads captured 623 μg/mL REE ions, corresponding to 3.9 REE ions per MIF molecule (fig. S24), closely matching the four high-affinity binding sites identified by ITC measurements (Fig. 3B). These results suggest that covalent immobilization retained the binding stoichiometry between REEs and the MIF in aqueous solution. We next loaded equimolar amounts of all 16 nonradioactive REEs onto the MIF separation column. Using a fully aqueous citrate system (without strong acids or bases), the MIF achieved single-stage separation of 16 distinct elution peaks (Fig. 4A). Sc^III^ eluted first, followed by lanthanides in descending order of atomic number (fig. S24). The 14 resolved peaks (including both single and overlapping peaks) presented purities of 87.0–99.2% and yields of 87.4–99.6% (fig. S24). Notably, pairs with sub-picometer ionic radius differences, Ho^III^-Y^III^ (Δr = 0.1 pm) and Pr^III^-Nd^III^ (Δr = 0.7 pm), initially showed overlapping peaks but could be separated with >90% purity and yield using modified elution conditions (see in supplementary methods) with the same MIF column (Fig. 4B and Fig. 4C).

**Fig. 4.**
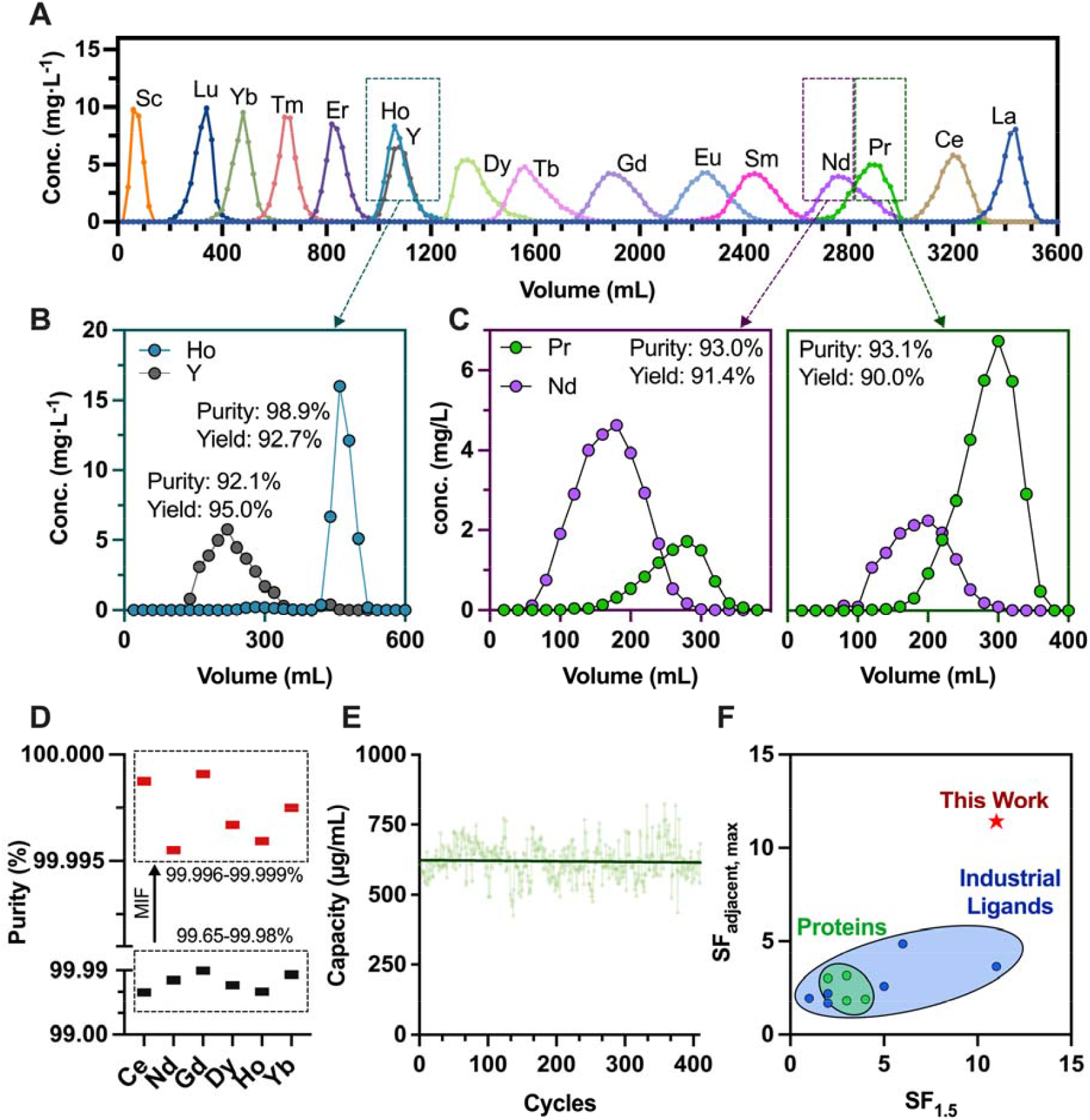
Full-spectrum REE separation and durability of the immobilized MIF. **(A)** Single-stage, all-aqueous separation of 16 REEs (Sc^III^, Y^III^ and lanthanides) by the MIF. **(B-C)** Second-stage separation of challenging REE pairs. **(B)** The Ho/Y pair was resolved via pH gradient elution. **(C)** Pr/Nd separations achieved > 90% purity and yield via rechromatography, respectively. **(D)** The MIF column was used for ultrahigh-purity REE preparation. Commercial light REEs (Ce, 99.65% and Nd, 99.84%), middle REEs (Gd, 99.98% and Dy, 99.76%), and heavy REEs (Ho, 99.66% and Yb, 99.93%) samples were investigated (in Black); the corresponding purities of Ce (99.997%), Nd (99.996%), Gd (99.999%), Dy (99.997%), Ho (99.996%), and Yb (99.999%) were determined by a single-stage MIF chromatographic process (in Red). **(E)** Durability assessment of the MIF-immobilized column after 410 adsorption□desorption cycles (La^III^ was used as a model ion). The binding capacity remained >95% of the initial value. **(F)** Ashby plot comparing the separation performance between the MIF and other ligands. SF_adjacent,max_ is defined as the largest separation factor among the 13 adjacent lanthanide pairs, whereas SF_1.5_ represents the number of the 13 pairs whose separation factor exceeded 1.5.

To validate the ability of the MIF to produce REEs with ultrahigh purity, we performed single-stage purification of representative light (Ce^III^, Nd^III^), middle (Gd^III^, Dy^III^), and heavy (Ho^III^, Yb^III^) REEs (Fig. 4D and fig. S25). The MIF enabled REE separation with purities up to 99.999%, representing the first protein-based platform to achieve ultrahigh-purity REE production. Long-term durability was also tested: the column retained >95% of its original capacity over 410 consecutive adsorption□desorption cycles (Fig. 4E), demonstrating the excellent stability of the MIF. Separation factors (SF_X-Y_, X, and Y represent paired REEs) were calculated for each paired REE (table S3). The average SF of the MIF between adjacent lanthanides reached 2.8, with a maximum value of 11.4 (SF_Sm/Nd_), and the SF for eleven adjacent lanthanide pairs exceeded 1.5 (Fig. 4F). Finally, we compared the REE separation efficacy of the MIF against both industrial ligands and previously reported lanthanide-binding proteins (*20, 24, 29, 33, 34*). These results highlight the superior adjacent REE separation performance of the MIF as an atomic ruler, featuring high selectivity for REEs with strong potential for industrial applications.

## Conclusion

We report a versatile, pollution-free, and structurally encodable approach for rare-earth separation enabled by an atomic ruler with a metalloprotein framework (MIF) structure. This atomic ruler allows single-stage, aqueous separation of 16 REEs with exceptional purity and yields, achieving ultrahigh purity levels (e.g., 99.999%). The MIF exhibits remarkable thermal stability, retaining its secondary structure even in boiling water (100°C) and remaining conformationally intact across heating□cooling cycles. When immobilized on a matrix, the rigid molecular framework maintains over 95% of its REE-selective binding capacity after 410 absorption□desorption cycles. These properties position the MIF as a rapidly deployable, protein-based platform for REE separation, greatly outperforming all other proteinaceous and traditional ligands in resolving adjacent lanthanides (*20, 23, 24, 35*).

The picometer-scale REE discrimination performance of the MIF can be attributed to its ultrarigid β-barrel backbone and densely packed multimetal ion stacking, which amplify sub-angstrom ionic radius differences into pronounced energy-driven discrimination. Tightly stacked REE ions generate cumulative steric mismatches within the channel-like microenvironment through bridging coordination, and these mismatches are effectively transduced by the rigid framework. This structural-deformation-energy-based discrimination achieves a resolution far exceeding that of conventional ligands. Our findings provide a pollution-free paradigm for REE separation using the MIF-based atomic ruler strategy, presenting a rigid protein framework for high-resolution, scalable REE isolation. More broadly, the metal-stacking amplification strategy demonstrated here exemplifies an encodable and structurally tunable platform for converting atomic-scale steric disparities into macroscale separation performance. This concept can be extended to the purification of other critical metals (e.g., actinides) and to applications such as metal-responsive biosensing for environmental detection of trace amounts.

## Supporting information

manucript-MIF

## Acknowledgments

**We thank Q. Yu, Y. Pang, and H. Su for their assistance with large-scale protein fermentation and purification, and Dr Q. Han at the Laboratory for Separation Science, Department of Chemistry, Tsinghua University, for performing elemental analyses**.

## Author contributions

Conceptualization: KL, LL, HJZ

Methodology: XNQ, YXD, CM, JY, HJC, HXZ

Investigation: XNQ, YXD, HXZ, TJS, YB, FKW, ZQL,

Visualization: XNQ, YXD, HXZ

Funding acquisition: KL, LL, HJZ

Project administration: KL, LL, HJZ, HJG

Supervision: KL, LL

Writing – original draft: XNQ, DYX, CM, HJC, HXZ, KL, LL

Writing – review & editing: KL, LL, JY, HJZ, HJG

## Data and materials availability

All data needed to evaluate the conclusions in the paper are present in the paper and/or the Supplementary Materials. Crystal structures of the apo MIF, MIF-La(III), MIF-Nd(III), MIF-Sm(III), MIF-Gd(III), MIF-Tb(III), MIF-Ho(III), and MIF-Yb(III) complexes have been published in the Protein Data Bank (PDB ID code 9VY3, 9VY7, 9VYA, 9VY8, 9VY9, 9VY5, 9VY6, and 9VY4, respectively). The apo WT LanM structure is available under PDB accession code 8DQ2.

## References and Notes

1. “Mineral commodity summaries 2025,” Mineral Commodity Summaries (Reston, VA, 2025).

2. Y. Bai et al., Harnessing Synthetic Biology for Sustainable Recovery of Critical Metal Materials from Electronic Waste. Advanced Functional Materials n/a, e09900.

3. D. S. Sholl, R. P. Lively, Seven chemical separations to change the world. Nature 532, 435–437 (2016).

4. S. Cotton, Lanthanide and actinide chemistry. (John Wiley & Sons, 2024).

5. F. H. Spedding, A. H. Daane, The Rare Earths. (Wiley, 1961).

6. R. Shannon, Revised effective ionic radii and systematic studies of interatomic distances in halides and chalcogenides. Acta Crystallographica Section A 32, 751–767 (1976).

7. T. Cheisson, E. J. Schelter, Rare earth elements: Mendeleev’s bane, modern marvels. Science 363, 489–493 (2019).

8. B. Weaver, F. A. Kappelmann, A. C. Topp, Quantity Separation of Rare Earths by Liquid—Liquid Extraction. I. The First Kilogram of Gadolinium Oxide1. Journal of the American Chemical Society 75, 3943–3945 (1953).

9. G. J. P. Deblonde, A. Chagnes, G. Cote, Recent Advances in the Chemistry of Hydrometallurgical Methods. Separation & Purification Reviews 52, 221–241 (2023).

10. D. Qi, in Hydrometallurgy of Rare Earths, D. Qi, Ed. (Elsevier, 2018), pp. 533–589.

11. F. Xie, T. A. Zhang, D. Dreisinger, F. Doyle, A critical review on solvent extraction of rare earths from aqueous solutions. Minerals Engineering 56, 10–28 (2014).

12. E. Vahidi, F. Zhao, Environmental life cycle assessment on the separation of rare earth oxides through solvent extraction. Journal of Environmental Management 203, 255–263 (2017).

13. X. Zheng, E. Liu, F. Zhang, Y. Yan, J. Pan, Efficient adsorption and separation of dysprosium from NdFeB magnets in an acidic system by ion imprinted mesoporous silica sealed in a dialysis bag. Green Chemistry 18, 5031–5040 (2016).

14. L. L. Perreault et al., Functionalization of Mesoporous Carbon Materials for Selective Separation of Lanthanides under Acidic Conditions. ACS Applied Materials & Interfaces 9, 12003–12012 (2017).

15. Y. Hu, E. Drouin, D. Larivière, F. Kleitz, F.-G. Fontaine, Highly Efficient and Selective Recovery of Rare Earth Elements Using Mesoporous Silica Functionalized by Preorganized Chelating Ligands. ACS Applied Materials & Interfaces 9, 38584–38593 (2017).

16. W. Zhang et al., Intralanthanide Separation on Layered Titanium(IV) Organophosphate Materials via a Selective Transmetalation Process. ACS Applied Materials & Interfaces 10, 22083–22093 (2018).

17. M. Mon et al., Lanthanide Discrimination with Hydroxyl-Decorated Flexible Metal–Organic Frameworks. Inorganic Chemistry 57, 13895–13900 (2018).

18. T. Cheisson, B. E. Cole, B. C. Manor, P. J. Carroll, E. J. Schelter, Phosphoryl-Ligand Adducts of Rare Earth-TriNOx Complexes: Systematic Studies and Implications for Separations Chemistry. ACS Sustainable Chemistry & Engineering 7, 4993–5001 (2019).

19. B. E. Cole et al., Understanding Molecular Factors That Determine Performance in the Rare Earth (TriNOx) Separations System. ACS Sustainable Chemistry & Engineering 8, 14786–14794 (2020).

20. D. Qi, in Hydrometallurgy of Rare Earths, D. Qi, Ed. (Elsevier, 2018), pp. 631–669.

21. X. Qian, C. Ma, H. Zhang, K. Liu, Bioseparation of rare earth elements and high value-added biomaterials applications. Bioorganic Chemistry 143, 107040 (2024).

22. Z. Dong et al., Bridging Hydrometallurgy and Biochemistry: A Protein-Based Process for Recovery and Separation of Rare Earth Elements. ACS Central Science 7, 1798–1808 (2021).

23. H. Cui et al., The Construction of a Microbial Synthesis System for Rare Earth Enrichment and Material Applications. Advanced Materials 35, 2303457 (2023).

24. J. A. Mattocks et al., Enhanced rare-earth separation with a metal-sensitive lanmodulin dimer. Nature 618, 87–93 (2023).

25. J. A. Cotruvo, Jr., E. R. Featherston, J. A. Mattocks, J. V. Ho, T. N. Laremore, Lanmodulin: A Highly Selective Lanthanide-Binding Protein from a Lanthanide-Utilizing Bacterium. Journal of the American Chemical Society 140, 15056–15061 (2018).

26. J. L. Hemmann et al., Lanpepsy is a novel lanthanide-binding protein involved in the lanthanide response of the obligate methylotroph Methylobacillus flagellatus. Journal of Biological Chemistry 299, (2023).

27. K. J. Franz, M. Nitz, B. Imperiali, Lanthanide-binding tags as versatile protein coexpression probes. ChemBioChem 4, 265–271 (2003).

28. H. Cui, F. Wang, C. Ma, H. Zhang, K. Liu, Microbial-driven fabrication of rare earth materials. Science China Materials 67, 2376–2392 (2024).

29. W. B. Larrinaga, J. J. Jung, C.-Y. Lin, A. K. Boal, J. A. Cotruvo, Modulating metal-centered dimerization of a lanthanide chaperone protein for separation of light lanthanides. Proceedings of the National Academy of Sciences 121, e2410926121 (2024).

30. Z. Yuewen, Q. Xining, M. Chao, L. Kai, Z. Hongjie, Rare Earth Biological Manufacturing and High Value-added Material Application★. Acta Chimica Sinica 81, 1624 (2023).

31. R. J. Ellis et al., In the Bottlebrush Garden: The Structural Aspects of Coordination Polymer Phases formed in Lanthanide Extraction with Alkyl Phosphoric Acids. The Journal of Physical Chemistry B 119, 11910–11927 (2015).

32. P. Weßling, U. Müllich, E. Guerinoni, A. Geist, P. J. Panak, Solvent extraction of An(III) and Ln(III) using TODGA in aromatic diluents to suppress third phase formation. Hydrometallurgy 192, 105248 (2020).

33. Z. Dong, J. A. Mattocks, J. A. Seidel, J. A. Cotruvo, D. M. Park, Protein-based approach for high-purity Sc, Y, and grouped lanthanide separation. Separation and Purification Technology 333, 125919 (2024).

34. D. Qi, in Hydrometallurgy of Rare Earths, D. Qi, Ed. (Elsevier, 2018), pp. 187–389.

35. J. A. Mattocks, J. A. Cotruvo, Biological, biomolecular, and bio-inspired strategies for detection, extraction, and separations of lanthanides and actinides. Chemical Society Reviews 49, 8315–8334 (2020).

36. T. Ku et al., Predicting melting temperature directly from protein sequences. Computational Biology and Chemistry 33, 445–450 (2009).

37. For exemple, Pr-Nd mixtures with 75% purity have already met the industrial requirements for permanent magnet.

38. K. Opelt et al., Upscaling the 2-Powder Method for the Manufacturing of Heavy Rare-Earth-Lean Sintered didymium-Based Magnets. Advanced Engineering Materials 23, 2100459 (2021).

39. B. J. Smith, M. E. Riddle, M. R. Earlam, C. Iloeje, D. Diamond, “Rare earth permanent magnets: supply chain deep dive assessment,” (USDOE Office of Policy (PO), Washington DC (United States), 2022).

40. M. Dembowski et al., Separation of rare earth element radioisotopes by reverse-phase high-speed counter-current chromatography. Journal of Chromatography A 1712, 464478 (2023).

41. J. Ward, B. Bucher, K. Carney, M. Snow, Exploring lanthanide separations using Eichrom’s Ln Resin and low-pressure liquid chromatography. Journal of Radioanalytical and Nuclear Chemistry 327, 307–316 (2021).

42. S. M. Qaim, H. Ollig, G. Blessing, Separation of Lanthanides by Preparative High Pressure Liquid Chromatography. Radiochimica Acta 26, 59–62 (1979).

43. M. Krättli, T. Müller-Späth, N. Ulmer, G. Ströhlein, M. Morbidelli, Separation of Lanthanides by Continuous Chromatography. Industrial & Engineering Chemistry Research 52, 8880–8886 (2013).

44. L. Ling, N.-H. L. Wang, Ligand-assisted elution chromatography for separation of lanthanides. Journal of Chromatography A 1389, 28–38 (2015).

45. C. Yeats, N. D. Rawlings, A. Bateman, The PepSY domain: a regulator of peptidase activity in the microbial environment? Trends in Biochemical Sciences 29, 169–172 (2004).

46. W. L. DeLano, S. Bromberg, PyMOL user’s guide. DeLano Scientific LLC 629, (2004).

47. W. Han, Y. Zhang, R. Liu, Y. Sun, M. Li, Solidification of glass-ceramics of simulated fission products RE3+, Sr2+, Ba2+ in chlorine-containing waste molten salt. Ceramics International 49, 15133–15144 (2023).

48. F. F. Khoury et al., Mining peptides for mining solutions: evaluation of calcium-binding peptides for rare earth element separations. Chemical Science, (2025).

49. Q. Ye, D. Wang, N. Wei, Engineering biomaterials for the recovery of rare earth elements. Trends in Biotechnology 42, 575–590 (2024).

