## Supplementary material for "Adjacent Rare-Earth Separation via a Protein Atomic Ruler": manucript-MIF

**The PDF file includes:**

Materials

MIF Synthesis and Metal-Binding Analysis

Crystallization and structure determination

Computational Methods

MIF Immobilization and REE separations

Figs. S1 to S25

Tables S1 to S10

References

Materials

All inorganic salts and chemicals were purchased from Shanghai Aladdin Bio-Chem Technology Co., Ltd. (Shanghai, China) at the highest available purity, unless otherwise specified. Ultrapure water (resistivity: 18.2 MΩ·cm at 25 °C) was obtained from a Create Fun ultrapure water system (Create Fun, Ltd., Chengdu, China). All chemicals used for MIF expression were used as received without further purification. DNA sequences for the MIF construct were synthesized by Genewiz (Shanghai, China). *Escherichia coli* BL21 (DE3) competent cells for MIF expression were obtained from Novagen (USA). The HisTrap FF column, Superdex 200 20/600 GL column, and Superdex 75 10/300 GL column used for MIF purification were purchased from Cytiva (USA). The Q Chromstar® HP anion exchange column was purchased from Bio-Link (China). Rare-earth elements (REEs) were quantified using inductively coupled plasma optical emission spectrometry (ICP-OES; Thermo Scientific iCAP 7400) and inductively coupled plasma mass spectrometry (ICP-MS; Thermo Scientific iCAP Q). A multi‑element ICP calibration solution (NCS 141749, 1000 µg/mL each element in 10 % HNO_3_, NCS Testing Technology Co., Ltd., Beijing, China) was used to prepare all external calibration standards for ICP‑OES and ICP‑MS measurements. Both ICP-OES and ICP-MS measurements were conducted in the Laboratory for Separation Science, Department of Chemistry, Tsinghua University. For protein immobilization, NHS-Activated Chromstar® 4FF agarose beads were purchased from Bio-Link and used without further purification.

MIF Synthesis and Metal-Binding Analysis

Expression in 5000 mL Shaker Flasks

The pET‑25b vectors carrying either the MIF sequence or its N‑terminal His10‑tagged variant (sequences in Supplementary Table S6) were synthesised by Genewiz (Shanghai, China). Each plasmid was separately transformed into *Escherichia coli* BL21 (DE3), and the resulting strains were expressed under identical conditions as described below. A seed culture was prepared by inoculating sterile 20 mL LB medium supplemented with 100 μg/mL ampicillin, followed by incubation at 37 °C and 220 rpm until the OD_600_ reached approximately 4. The 20 mL seed culture was then transferred into 1000 mL Terrific Broth (TB) medium in a 5 L shaker flask (2% v/v) and grown under identical conditions until the OD_600_ reached approximately 4 (about 4 hours). The culture, as well as the shaker, was then cooled to 16 °C for 30 minutes before induction with 1 mM isopropyl β-D-1-thiogalactopyranoside (IPTG). The induced culture was incubated for 16-18 hours at 16 °C and 220 rpm before harvesting the cells by centrifugation at 8000 × g and 4 °C for 20 minutes (Fiberlite F9-6 x 1000 LEX Fixed Angle Rotor for LYNX 6000 Superspeed Centrifuge). Cells were flash‑frozen in liquid N₂ and stored at -80 °C.

Expression in 50 L Bioreactor

For large-scale expression, 1000 mL of the seed culture grown in LB medium was inoculated into 30 L (~3% v/v) sterile bacterial culture medium containing 6.67 g/L KH_2_PO_4_, 13 g/L peptone, 3.25 g/L citric acid monohydrate, 0.05 g/L CaCl_2_·H_2_O, 2.5 g/L MgSO_4_·7H_2_O, 5 mg/L FeSO_4_·7H_2_O, 200 μg/mL ampicillin, and 5 g/L glucose. The pH of the medium was adjusted to 6.8–7.0 using 28% ammonia solution. The fermentation was carried out in a 50 L bioreactor with 30 L of working volume (60 % of total) at 37 °C, 500 rpm, with aeration at 1–3 vvm (volume of air per volume of culture per minute) and an internal pressure of 0.02 MPa. The pH was maintained between 6.8 and 7.0 by the addition of 28% ammonia solution. After approximately 3 hours of cultivation, when the dissolved oxygen (DO) level sharply increased (indicating the glucose was nearly depleted), sterile glucose solution was continuously fed to maintain the DO level around 30%. When the OD600 reached approximately 55, glucose feeding was stopped, and wait until the DO level increase to approximately 55%. The culture was then cooled to 16 °C in ~30 min by chilled water jacket, and IPTG was added to a final concentration of 0.5 mM to induce protein expression. The entire induction process lasted 12 hours. Glucose feeding was resumed to maintain the DO level at approximately 70% until 2 hours before the end of fermentation. Finally, the cells were harvested by centrifugation at 8000 × g and 4 °C for 20 minutes (Fiberlite F9-6 x 1000 LEX Fixed Angle Rotor for LYNX 6000 Superspeed Centrifuge). Cells were flash‑frozen in liquid N₂ and stored at -80 °C.

Purification of His_10_-tagged MIF

Cells were resuspended in Ni lysis buffer consisting of 50 mM sodium phosphate buffer (pH 7.4), 10mM imidazole, and 500 mM NaCl to a final concentration of 10–20% (w/v), and then lysed using a high-pressure cell disruptor (ATS Ltd., Jiangsu, China). The cell lysate was clarified by centrifugation at 16,000 × g for 30 minutes at 4 °C (Fiberlite F14-6x250y Fixed Angle Rotor for LYNX 6000 Superspeed Centrifuge), followed by filtration through a 0.45 μm membrane to remove residual debris. The resulting supernatant was loaded onto a pre-equilibrated HisTrap FF 70 mL column (Cytiva) for nickel-affinity purification. All buffers used in protein purification were 0.45 µm filtered and degassed. All chromatographic steps were performed at room temperature (25 °C), unless otherwise specified. The column was washed with 10 column volumes (CV) of buffer containing 50 mM sodium phosphate (pH 7.4), 500 mM NaCl, and 90 mM imidazole. A pre‑screened step‑gradient had shown that 90 mM imidazole was sufficient to remove non‑specific host proteins. His_10_-tagged MIF was eluted using the same buffer supplemented with 6 CV of 350 mM imidazole. The flow rate is 20 mL/min for loading, elution, and washing. The eluted protein was desalted and concentrated using a Sartorius ultrafiltration system with a 10-kDa cutoff membrane. Five cycles of 10 × volume of deionized water were used for buffer exchange to remove imidazole/salt. The purified His10-tagged MIF was either lyophilized for long-term storage or stored as an aqueous solution at 4 °C in either deionized water or MES buffer (25 mM MES, 150 mM NaCl, pH 6.0) for immediate use.

Purification of untagged MIF

Cells were resuspended in lysis buffer containing 20 mM Tris and 20 mM NaCl (pH 5.9) to a final concentration of 10–20% (w/v), and lysed using a high-pressure cell disruptor (ATS Ltd., Jiangsu, China). The cell lysate was clarified by centrifugation at 16,000 × g for 30 minutes at 4 °C, followed by filtration through a 0.45 μm membrane to remove residual debris. All buffers used in protein purification were 0.45 µm filtered and degassed. All chromatographic steps were performed at room temperature (25 °C), unless otherwise specified. The resulting supernatant was loaded onto a pre-equilibrated Q Chromstar® HP 70 mL column (Bio-Link) for anion exchange chromatography. The flow rate is 20 mL/min. The column was washed with 15 CV of buffer containing 20 mM Tris and 28.4 mM NaCl (pH 5.9) to remove contaminant proteins. MIF was eluted using a 6 CV of linear NaCl gradient ranging from 28.4 mM to 32.8 mM (pre‑screened step‑gradient indicated elution between 28-33 mM) while maintaining constant buffer composition and pH. Eluted fractions were analyzed by SDS-PAGE, and those containing MIF were pooled and concentrated. The pooled protein solution was desalted and concentrated using a Sartorius ultrafiltration system with a 10 kDa molecular weight cutoff membrane and buffer-exchanged into 20 mM Tris and 20 mM NaCl (pH 5.9) after five cycles of 10 × volume. The resulting concentrated sample (sample load < 4 mL) was further purified by size-exclusion chromatography (SEC) on a Superdex 200 20/600 GL column (Cytiva) equilibrated with the same buffer, at a flow rate of 1 mL/min. MIF-containing fractions were pooled, concentrated, and subjected to a second round of SEC using a Superdex 75 10/300 GL (Cytiva) column (sample load 500 µL, the chromatography was performed at 4 °C at a flow rate of 0.5 mL/min) to achieve higher purity. The final purified protein was concentrated using a 10 kDa centrifugal filter unit (Amicon Ultra, Merck).

The purified MIF was either supplemented with 20% (v/v) glycerol and stored at –20 °C for long-term preservation, or stored as an aqueous solution at 4 °C in either deionized water or MES buffer (25 mM MES, 150 mM NaCl, pH 6.0) for immediate use.

SDS-PAGE analysis of MIF

Cells (wet weight : lysis buffer = 1:9, m/v) were resuspended in NP‑40 Lysis Buffer (MCE, HY‑K1002) and incubated for 30 min at room temperature with gentle agitation. Lysates were clarified by centrifugation at 14 000 × g, 4 °C, 5 min, and 30 µL of the supernatant was mixed with 10 µL of 4 × SDS‑PAGE loading buffer (LabLead, G2526‑1) supplemented with dithiothreitol to give a final DTT concentration of 1 mM. A standard sample containing 30 µL of 1 mg/mL⁻ purified MIF in MES buffer was prepared in the same way. All samples were boiled for 10 min before loading. 10 µL of each reduced sample was applied per lane on LabPAGE 4-12％ 12 Wells (LabLead, P41212), together with a Prestained Protein Marker (10-180 kD, Servicebio, G2091-250UL, or BeyoColor 10-170 kD, Beyotime, P0077). Electrophoresis was carried out in Tris-MOPS-SDS running buffer (LabLead, T7205M/B) at 140 V constant voltage for ~50 min, until the tracking dye reached the bottom edge of the gel cassette. After electrophoresis, the gels were stained with eStain L1 Protein Staining System (GenScript, USA). Images of the stained gels were captured with a fully automated chemiluminescence imaging system (Tanon 5200, Tanon Science & Technology, China). The yield of MIF was estimated by comparing the band intensity of the cell lysis with that of a 1 mg/mL MIF standard solution on the same SDS-PAGE gel. Band intensities were quantified using ImageJ, and background was subtracted using the rolling‑ball algorithm (radius = 50 pixels).

Variable-temperature circular dichroism spectroscopy analysis of MIF

VT-CD data of MIF was collected by using a Chirascan Plus CD spectrometer (Applied Photophysics). For sample preparation, the apo-MIF was buffer-exchanged into CD buffer (25 mM MES, pH 6.2) using a Superdex 75 Increase 10/300 GL column (GE Healthcare). The proteins were then incubated with the corresponding REEs (La, Sm, and Yb) in a 1:10 stoichiometric ratio and diluted to a concentration of 0.1-0.2 mg/mL. The measurement parameters were set up as follows: wavelength, 200-280 nm; step resolution, 1 nm; bandwidth, 1nm; temperature, from 25 to 100℃ in stepped ramping mode. CD spectra of proteins were obtained 3 times from 200 to 280 nm in 1 mm cuvettes every 5℃ increase. CD spectra were collected again when the sample was cooled to 25℃. The curves were smoothed using standard parameters. The secondary structure content of MIF was analyzed by using software CDNN 2.1 with standard parameters (*1*).

Matrix-Assisted Laser Desorption/Ionization Time-of-Flight Mass spectrometry analysis of MIF

A polished ground steel MALDI target plate (Bruker Daltonik GmbH, Bremen, Germany) was used for MALDI‐TOF MS measurements, which were performed with UltrafleXtreme TOF mass spectrometer (Bruker Daltonik GmbH, Bremen, Germany).  The mixture of sample (0.5 μL) and SA matrix solution (0.5  μL, 10 mg/ml) was spotted onto the plate. The MS is equipped with a 2 kHz smartbeam‐II laser and controlled by the FlexControl software (version: 3.4, Bruker Daltonik GmbH, Bremen, Germany). Spectra were recorded in positive linear mode in the mass range between 10000 and 2100 00 Da. The parameter settings were optimized as follows: ion source 1:20.00 kV, ion source 2:18.50 kV, pulsed ion extraction time: 550 ns.

BSA was used for instrument calibration as an external standard. All spectra are based on the average of 1000 pulses, accumulated from different points of the target. Data processing was performed using the FlexAnalysis software (version: 3.4, Bruker Daltonik GmbH, Bremen, Germany).

Quadrupole Time-of-Flight Mass spectrometry analysis of MIF

For LC-MS analysis, the analytes were separated by a 60 min gradient elution at a flow rate of 0.5 µl/min with a nanoACQUITY UPLC system, which was directly interfaced with a SYNAPT-G2-Si mass spectrometer produced by the Waters company. The analytical column was a Protein BEH C4 silica capillary column (150 µm ID, 100 mm length; Made in Ireland) packed with C-4 resin (300 Å, 1.7 µm) purchased from the Waters company. Mobile phase A consisted of 0.1% formic acid aqueous solution, and mobile phase B consisted of 100% acetonitrile and 0.1% formic acid. Aliquots of 3 µl of analytes were loaded into an autosampler for nanoelectrospray ionization. Samples were analyzed on a Q-TOF mass spectrometer (SYNAPT G2-Si, Waters company) instrument optimized for high-mass protein analysis. The measurements were performed with capillary 2000–2500 V, and data were collected over the m/z range of 500–3000. Once having acquired raw electrospray mass spectra, the raw spectrum can be deconvoluted by MaxEnt 1 (Waters) to generate a spectrum (relative intensity versus mass) where all the charge-state peaks of a single species have been collapsed into a single (zero-charge) peak.

Proteomic analysis of MIF

In-solution Digestion

The sample was reduced disulfide bonds and carbamidomethylated cysteine residues by adding a 1:10 volume of reduction/alkylation buffer （Reduction/alkylation buffer contains 100 mM TCEP and 400 mM 2-chloroacetamide (CAM), brought to pH 7–8 with KOH）to the samples and incubating for 1h at 25 °C in the dark. Digestion was performed by adding trypsin (Promega) at a 1:50 enzyme-to-substrate ratio and incubating overnight at 37 °C. Digests were acidified by the addition of neat trifluoroacetic acid (TFA) to 1%, centrifuged to pellet insoluble matter, and the supernatant was dried by vacuum centrifugation.

LC-MS/MS Analysis

The digestion products were separated by a 60 min gradient elution at a flow rate of 0.300 µL/min with a Thermo-Dionex Ultimate 3000 HPLC system, which was directly interfaced with the Thermo Scientific™ Orbitrap FusionTM mass spectrometer. The analytical column was a home-made fused silica capillary column (100 µm ID, 150 mm length; Upchurch, Oak Harbor, WA) packed with C-18 resin (120 Å, 1.9 µm, Dr.Maisch, Ammerbuch, Germany). Mobile phase A consisted of 0.1% formic acid, and mobile phase B consisted of 80% acetonitrile and 0.1% formic acid. The mass spectrometer was operated in the data-dependent acquisition mode using Xcalibur 4.1 software, and there is a single full-scan mass spectrum in the Orbitrap (300-1500 m/z, 120,000) resolution, followed by top-speed MS/MS scans in the Orbitrap.

The MS/MS spectra from each LC-MS/MS run were searched against the target sequence using Proteome Discoverer (Version PD 2.5, Thermo-Fisher Scientific, USA). The search criteria were set as follows: full tryptic specificity was required, and two missed cleavage sites were allowed. Oxidation (M) was set as variable modification, and Carbamidomethyl (C) was set as fixed modification; precursor ion mass tolerances were set at 20 ppm for all MS acquired in an Orbitrap mass analyzer, and the fragment ion mass tolerance was set at 0.02 Da for all MS2 spectra acquired.

Isothermal titration calorimetry

Isothermal titration calorimetry (ITC) measurements were performed using a MicroCal PEAQ-ITC instrument (Malvern Panalytical). The sample cell was filled with 280 µL of purified, untagged MIF dissolved in a buffer containing 25 mM MES and 150 mM NaCl at pH 6.0. The syringe (38.9 µL) was loaded with REE ions dissolved in the same buffer to avoid buffer mismatch. Both solutions were adjusted to pH 6 with 1 M HCl or NaOH, passed through a 0.22 µm PVDF syringe filter before loading, and no bubbles were detected in the cell or syringe. Preliminary trials showed dilution heats of metal‑into‑buffer to be within the baseline noise. For each REE-MIF titration, the concentration of MIF (cell) and REE (syringe), the number of total injections, the volume of titrant injected per shot, and the intervals were shown in Supplementary Table S7. The first injection volume was reduced to 0.2 μL. The experiments were carried out at 25 °C, with a stirring speed of 750 rpm and a reference power of 20.9 μW. No visible turbidity or precipitation was observed before or after the titration. All ITC data were baseline-corrected, fitted, and plotted using the manufacturer’s PEAQ-ITC Analysis Software (Version: 1.41). For La^III^, Nd^III^, Sm^III^, Gd^III^, Tb^III^, and Tm^III^, the enthalpy data (ΔH) were fitted with a one‑set‑of‑sites model and five binding sites, and each titration was performed in triplicate (n = 3). Dissociation constants for all lanthanides were obtained by fitting the integrated heats to a sequential binding model (*2*).

Crystallization and structure determination

To obtain the crystal structure of apo MIF, the protein was concentrated to 15 mg/mL in buffer consisting of 20 mM HEPES (pH 7.0), 120 mM NaCl, 1 mM DL-dithiothreitol, and then crystallized in the reservoir buffer of 0.2 M Ammonium fluoride, 20% w/v Polyethylene glycol 3,350 with sitting-drop vapor diffusion. Crystallization was under 291K, and each drop contained 0.5 μL protein solution and 0.5 μL crystallization reservoir buffer. For the crystal structure of MIF-REE complexes, 1.0 mg/mL MIF was incubated with 8.0 mM lanthanide chloride for 30-60 minutes at 4 °C, followed by concentrated MIF to 13.5 mg/mL, which yielded a final stoichiometric ratio of 1:10 between the protein and REE ions. The crystallization procedure was consistent with that of apo MIF, except that the composition of the reservoir solutions varied among different MIF-REE complexes: MIF-La^III^, 0.2 M ammonium acetate, 0.1 M BIS-TRIS pH 5.5, 25% w/v Polyethylene glycol 3,350; MIF-Nd^III^, 0.1 M BIS-TRIS pH 5.5, 25% w/v polyethylene glycol 3,350; MIF-Sm^III^, 0.1 M BIS-TRIS pH 6.1, 29.4% w/v polyethylene glycol 3,350, 0.5% w/v n-octyl-β-D-glucoside; MIF-Gd^III^, 0.15 M KBr, 30% w/v polyethylene glycol monomethyl ether 2,000, 10 mM sarcosine; MIF-Tb^III^, 10% v/v 2-propanol, 0.1 M BICINE pH 8.5, 30% polyethylene glycol 3,350; MIF-Ho^III^, 0.2 M ammonium sulfate, 0.1 M MES pH 6.5, 20% w/v Polyethylene glycol 8,000; MIF-Yb^III^, 0.1 M MES pH 6.5, 25% w/v Polyethylene glycol 4,000, 0.7% v/v 1-butanol. All harvested crystals were directly flash frozen and preserved in liquid nitrogen.

Diffraction data of all the crystals, including the apo MIF and MIF-La^III^, MIF-Nd^III^, MIF-Sm^III^, MIF-Gd^III^, MIF-Tb^III^, MIF-Ho^III^, MIF-Yb^III^ complexes, were collected under the wavelength of 0.979176 Å and temperature of 100 K at the SSRF BEAMLINE BL02U1 of Shanghai Synchrotron Radiation Facility. Raw diffraction data were processed and scaled by Aimless 0.7.4 (*3*) and HKL2000 (*4*). Molecular replacement was performed for phasing of data in PHENIX 1.19.2_4158 (*5*). We used the in-silico model of MIF predicted by AlphaFold3 (*6*) (AF-Q1H2G7-F1) for molecular replacement phasing of apo MIF data, while utilizing the chain A of the refined apo MIF’s structure for phasing of all MIF-REE complexes data. The model building was visualized and operated by COOT (*7*), and REE ions are initially placed at the peak of the difference map (F_o_-F_c_). Finally, all crystal structures were refined in PHENIX 1.19.2_4158 (*5*). Collection and refinement statistics of our structures are presented in Supplementary Table S8.

Computational Methods

Bioinformatics methods

All reported lanthanide-binding proteins were collected through a comprehensive literature and database search (data retrieved by Jan. 2025). Specifically, the Web of Science database was searched using the keywords “lanthanide binding protein” and “rare earth binding protein.” In parallel, the Protein Data Bank (PDB) was queried for structures containing lanthanide ions in which the metal was coordinated by at least three protein residues within a monomer or oligomeric assembly. The amino acid sequences of all identified proteins were retrieved and curated. The theoretical molecular weights of the proteins were calculated using the ProtParam tool (ExPASy) (*8*), and based on these values, the REE payload (defined as the number of lanthanide ions bound per kilodalton of protein) was determined. The thermostability of each protein was predicted using the Tm Predictor to obtain the melting temperature index (Tm index) (*9*). For proteins containing N-terminal signal peptides, predictions were performed using SignalP 6.0 (*10*), and the signal peptide regions were removed before further analysis.

All-atom molecular dynamics simulations

All-atom molecular dynamics (MD) simulations were performed using the GROMACS software package (*11, 12*). Each Ln³⁺-MIF system was modeled with the CHARMM36 force field (*13*), while the solvent environment was represented using the modified TIP3P water model under neutral pH conditions. Lanthanide ions (La^III^, Nd^III^, Sm^III^, Gd^III^, Tb^III^, Yb^III^) were modeled using the improved CHARMM-compatible parameters developed by Migliorati et al. (*14*), which enable efficient and accurate molecular dynamics simulations of lanthanide speciation in aqueous environments. These additive force field parameters reproduce key experimental observables, including hydration free energies and first-shell coordination features. Each REE-MIF complex was solvated in a cubic box of explicit water molecules, with periodic boundary conditions applied in in the x-, y-, and z-three directions. All glutamate (Glu) and aspartate (Asp) residues were modeled in their deprotonated (charged) forms, while other titratable residues were maintained in their neutral states. Sodium and chloride ions were added to neutralize the system and to achieve a physiological ionic strength of 150 mM NaCl. Long-range electrostatic interactions were calculated using the Particle Mesh Ewald (PME) algorithm (*15*), with a real-space cutoff of 1.2 nm applied to both electrostatic and Lennard-Jones interactions. All covalent bonds involving hydrogen atoms were constrained using the LINCS algorithm (*16*), allowing the use of an integration time step of 2 fs. Initial configurations for each Ln³⁺-MIF complex were obtained from their respective X-ray measurements. Prior to production simulations, each solvated system underwent energy minimization using the steepest descent (SD) algorithm to remove steric clashes and optimize geometry. This was followed by two equilibration phases: a 100 ps isochoric isothermal (NVT) simulation at room temperature of 298.15 K using a stochastic velocity rescaling algorithm (*17*), followed by a 100 ps isobaric isothermal (NPT) simulation under the room temperature and a pressure of 1 atm. Production MD simulations were then performed for 3μs in the NPT ensemble, maintaining constant temperature and pressure using the same thermostat and Parrinello-Rahman barostat schemes (*18*). A time step of 2 fs was employed throughout all simulations.

The two-step energy calculation approach

We constructed the hypothetical Deformed Apo-Like Conformation (DALC) for each lanthanide by individually removing the metal ions from the corresponding REE-MIF crystal structures. The electron energies of each DALC and the apo-MIF were computed, and the energy difference of DALC to apo-MIF was defined as the deformation energy (E_def_). The binding energy (E_bind_) between each DALC and its corresponding REE ions was then calculated. Since the protein framework in the DALC had already undergone conformational deformation, only the REE ions with coordination associated molecules (residues and water) were included in the quantum mechanical model, making DFT-based calculations feasible.

Calculation of deformation energy

The deformation energy (E_def_) was calculated as the MIF’s formation heat difference between the DALC (E_DALC_) and its ion-free state (E_Apo_):

E_def_ = E_DALC_ - E_Apo_

The structure optimization was performed exclusively on H atoms of the DALC using the CHARMM36 force field (*13*). The deformation energy calculations were computed using the PM6-D3H4X method, which incorporated corrections for London dispersion forces as well as hydrogen and halogen bonding interactions (*19, 20*). All semi-empirical calculations were performed using MOPAC, for which the MOZYME linear scaling algorithm was employed during computation (*21*).

Calculation of binding energy

The binding energy (E_bind_) between the DALC and rare earth ions was computed as the difference between the energy of the REE-MIF complex (E_AB_) and the sum of individual energies of the isolated DALC and ions (E_A_ + E_B_) (*22*):

ΔE_bind_ = E_AB_ - (E_A_ + E_B_)

Since the protein framework in the DALC had already undergone conformational deformation, we employed the quantum chemical cluster approach to compute binding energies by selecting a relatively small part of the REE-MIF complex as the model (*23*). The models were constructed based on the structural snapshot taken at 3 μs during the MD simulation of REE-MIF complexes. Each cluster model comprised the rare-earth ions along with coordinating residues and water molecules within a 4 Å coordination sphere. To simplify the system while preserving key structural features, all backbone atoms, except C-α at the edge of the model, were removed and subsequently saturated with H atoms. During geometry optimization, we allowed full relaxation of the rare-earth ions, their coordinating ligands, adjacent methylene groups, and all H atoms, while keeping the remaining carbon atoms in order to mimic the coordination microenvironment around the binding sites and to prevent excessive structural distortion.

All DFT calculations were performed with Gaussian 09. The models were optimized at the PBE0-D3(BJ) level within the IEFPCM model (water) (*24-27*). The def2-SVP basis set was applied to all groups directly coordinating the rare-earth ions, while the 3-21G basis set was utilized for the remaining atoms in the model (*28, 29*). Pseudopotential basis sets were adopted for La (MWB46), Gd (MWB53), Sm (MWB51), Nd (MWB49), Tb (MWB54), and Yb (MWB59) elements. The def2-TZVP basis set and same pseudopotential basis sets were employed for single-point energy calculations of the REE-MIF complex, using the SMD model (water) (*30*).

MIF Immobilization and REE separations

Immobilization of MIF

130 mg of purified His_10_-tagged MIF was dissolved in 6 mL of coupling buffer consisting of 0.1 M sodium bicarbonate and 0.5 M NaCl at pH 8.3. Simultaneously, 6 mL of NHS-activated agarose beads (Chromstar® 4FF, Bio-Link) were transferred into a 50 mL three-necked flask and washed 10 times with 18 mL of 1 mM HCl on ice to remove residual contaminants and prevent premature NHS hydrolysis. After washing, the beads were suspended in 9 mL of coupling buffer, and the pH was adjusted to 8.5. The dissolved MIF solution was then added, and the pH was carefully readjusted to 8.3. The coupling reaction was performed at 24 °C with gentle stirring. The absorbance at 280 nm (A280) of the supernatant was monitored periodically. The reaction was considered complete when the A280 value stabilized, indicating saturation of the coupling sites. Following protein coupling, the reaction mixture was removed, and the beads were incubated with 18 mL of blocking buffer (0.1 M Tris-HCl, pH 8.5) to quench any remaining active NHS groups. The A280 of the supernatant was again monitored until stabilization. The beads were then washed 10 times with 18 mL of 0.1 M acetic acid on ice, followed by 10 washes with 18 mL of ultrapure water. The resulting MIF-immobilized agarose beads were stored in ultrapure water at 4 °C until further use. The immobilized His_10_-tagged MIF (molecular weight 18.63 kDa) loading was approximately 21 mg/mL. For larger-scale preparations, all reagent volumes and protein quantities can be increased proportionally to obtain the desired amount of immobilized MIF under the same pH, temperature, and stirring conditions.

Batch experiment to determine separation factors

MIF-immobilized agarose microbeads were washed thoroughly with deionized water. For each batch adsorption experiment, 900 μL of feed solution containing an equimolar mixture of adjacent lanthanides was added to 100 μL of MIF-immobilized microbeads and incubated for 2 hours at 25 °C with gentle shaking (60 rpm). After incubation, the supernatant was collected, and the concentrations of REEs were quantified by inductively coupled plasma optical emission spectrometry (ICP-OES) after 10-fold dilution. The measured values were recorded as [Free]. Subsequently, 900 μL of 10 g/L sodium citrate solution in MES buffer (25 mM MES, pH 6.0) was used to desorb the bound REEs from the microbeads. The desorbed REE concentrations were measured by ICP-OES and recorded as [Desorb].

The distribution value (D) of each REE between the MIF phase and the solution phase was calculated as:

D = [Bound]/(10×[Free]).

where [Bound] and [Free] represent the molar concentrations of the metal ion in the microbead phase and the aqueous phase at equilibrium, respectively. To account for the free solution volume retained within the agarose beads, the following correction was applied:

[Bound]=[Desorb]−[Free]

Ref. (*31*) shows that distribution values (D) from separate experiments can be normalized without affecting the calculated separation factors (SF). This allows the calculation of binding affinity from several sets of experiments that share the same elements. We therefore divided the lanthanide series into 13 adjacent pairs. For each pair, we scaled the D value so that the first element kept the same D it had in the previous pair. D value of La^III^ was fixed to 1 as the reference (D_La_ = 1).

The SF between two REEs (X and Y) was defined as:

SF_X−Y_ = D_X_/D_Y_

where D_X_ and D_Y_ are the respective distribution values of REE_X_ and REE_Y_.

Breakthrough column experiments

Columns were filled as described in our previous work (*32*), at a column volume of 1.2 mL. Then, the column was washed with ultrapure water, 10 g/L sodium citrate solution in MES buffer (25 mM MES, pH 6.0), and conditioned with MES buffer as equilibrium solution before conducting breakthrough experiments.

La^III^ was chosen as a model ion, and the loading solution was prepared by dissolving lanthanum chloride in MES buffer. The concentration of La^III^ in the solution is 96.35 mg/L, determined by ICP-OES. The REE solutions were added at 1 mL/min to the column, and the column effluent was collected in 1.0 mL. ICP-OES was used to determine the concentrations of each REE.

Separation of Adjacent Rare-Earth Element Pairs

For the separation of adjacent REE pairs, each REE mixture solution was prepared by dissolving the chloride of two adjacent REEs in MES buffer; the concentrations are shown in Supplementary Table S2. 100 mL REE mixture solution was loaded onto a 75 mL MIF-immobilized column (95.5 cm, r = 0.5 cm) pre-equilibrated with the same MES buffer at a flow rate of 2 mL/min. After sample loading, the column was re-equilibrated with 150 mL MES buffer, followed by elution using a citrate gradient (shown in Supplementary Table S2) in MES buffer. Fractions were collected at 10 mL intervals throughout the elution process. The concentration of each REE in the collected fractions was quantified by ICP-OES and ICP-MS. The purities were calculated as purity [REE1] = [REE1] / [total REE]. REEs with concentrations below the detection limit (50ppb for ICP-OES, 5ppt for ICP-MS) were considered as zero in the purity calculations.

Full-spectrum REE separation experiments

For full-spectrum REE separation, the loading solution was prepared by dissolving the chloride of the 16 non-radioactive REEs (La, Ce, Pr, Nd, Sm, Eu, Gd, Tb, Dy, Ho, Er, Tm, Yb, Lu, Sc, Y) in MES buffer; the concentrations are shown in Supplementary Table S9. 80 mL loading solution was loaded onto a 150 mL MIF-immobilized column (191 cm, r = 0.5 cm) pre-equilibrated with the same MES buffer at a flow rate of 1 mL/min. After loading and re-equilibration with MES buffer, elution was performed using a citrate gradient ranging from 60 to 360 mg/L in MES buffer (pH 6.0). The entire elution process spanned 3600 mL. Fractions were collected at 20 mL intervals and analyzed for REE concentrations using ICP-OES. The purity of each element was calculated based on the relative concentrations of the target REE and total REE according to the method described earlier.

Modified elution for Pr^III^-Nd^III^ and Ho^III^-Y^III^ separation experiments

Pretreatment of fractions

REE-rich fractions collected from the full-spectrum run were transferred to PTFE beakers, concentrated to near dryness on a 60-80 °C hotplate, and digested with concentrated HNO_3_ (semiconductor grade). After cooling, the digest was diluted with ultrapure water and the pH was carefully adjusted to 4.0 with 1 M NaOH.

Pr^III^–Nd^III^ separation by re-chromatography

The loading solution was diluted with MES buffer (25 mM MES and 150 mM NaCl, pH 6.0) to 50 mL total volume and readjusted to pH 6.0. The sample was loaded at 1.0 mL/min onto the same 150 mL MIF-immobilized column (191 cm, r = 0.5 cm), pre-equilibrated in MES buffer. After loading and re-equilibration with MES buffer, elution was performed with a linear citrate gradient from 240 to 300 mg/L (in MES, pH 6.0) over 400 mL. Fractions were collected every 20 mL and analyzed by ICP-OES. The purity of each element was calculated according to the method described earlier.

Ho^III^–Y^III^ separation via pH gradient elution

After the same digestion, the pH 4.0 residue was diluted to 50 mL with 100 mg/L NaCl at pH 4.0 (adjusted with HCl). The column was pre-equilibrated with 100 mg/L NaCl at pH 4.0 (adjusted with HCl). The sample was loaded at 1.0 mL min⁻¹, followed by re-equilibration. Elution used a linear pH gradient from 4.0 to 1.0 in 100 mg/L NaCl, decreasing by 0.1 pH unit per 20 mL (total 600 mL). Fractions (20 mL) were analyzed by ICP-OES; purity was computed as above.

Ultra-high purity REE preparation

Ultra-high-purity REEs were prepared using commercial REE chlorides purchased from Energy Chemical (Anachem, China) as starting materials. The purity and loading amount of each starting material are shown in Supplementary Table S10, determined by ICP-MS. The REE chlorides were dissolved in MES buffer (25 mM, pH 6.0) and loaded onto a 150 mL MIF-immobilized column (191 cm, r = 0.5 cm) pre-equilibrated with the same MES buffer at a flow rate of 1 mL/min. After sample loading, the column was re‑equilibrated with the same MES buffer (25 mM, pH 6.0) and then eluted with a three-step gradient elution with citrate in MES buffer. Specifically, the first step is impurity wash. A 140 mL linear gradient raised the citrate concentration from 0 to every REE's corresponding "impurity wash" concentration, which are listed in Supplementary Table S10, followed by an isocratic wash at that concentration for a further 800 mL. The second step is a linear elution. The citrate concentration was then increased linearly over 2000 mL to the endpoint shown in Supplementary Table S10, releasing the target element as a single peak. The third step is a strip to all REEs and regeneration to the column. The citrate concentration step increased to 2 g L⁻¹ over 140 mL to wash any remaining REEs, after which the column was re‑equilibrated with fresh MES buffer for the next run. Fractions were collected at 20 mL intervals throughout the elution process. The concentration of the target REE in each fraction was determined by inductively coupled plasma optical emission spectrometry (ICP-OES). The concentrations of residual impurity REEs were quantified by inductively coupled plasma mass spectrometry (ICP-MS). The purity of each collected fraction was calculated based on the relative concentrations of the target REE and impurity REEs according to the method described earlier.

Durability of MIF-immobilized agarose beads

Columns were filled as described in our previous work (*32*), using a volume of 0.4 mL per column. A lanthanum chloride solution in MES buffer (25 mM, pH 6.0) with a La^III^ concentration of approximately 1 g/L served as the feed. After each loading step (2 mL), the column was washed with 10 mL of deionized water and then eluted with 2 mL of MES buffer containing 10 g/L sodium citrate (pH 6.0). An additional 10 mL of MES buffer wash was performed before the next cycle. The adsorption-desorption cycle was repeated 410 times within the same column. The total La^III^ amount in the effluent from the 2 mL citrate elution and the 10 mL MES buffer wash was determined by ICP‑OES and defined as the capacity.

Supplementary Figures

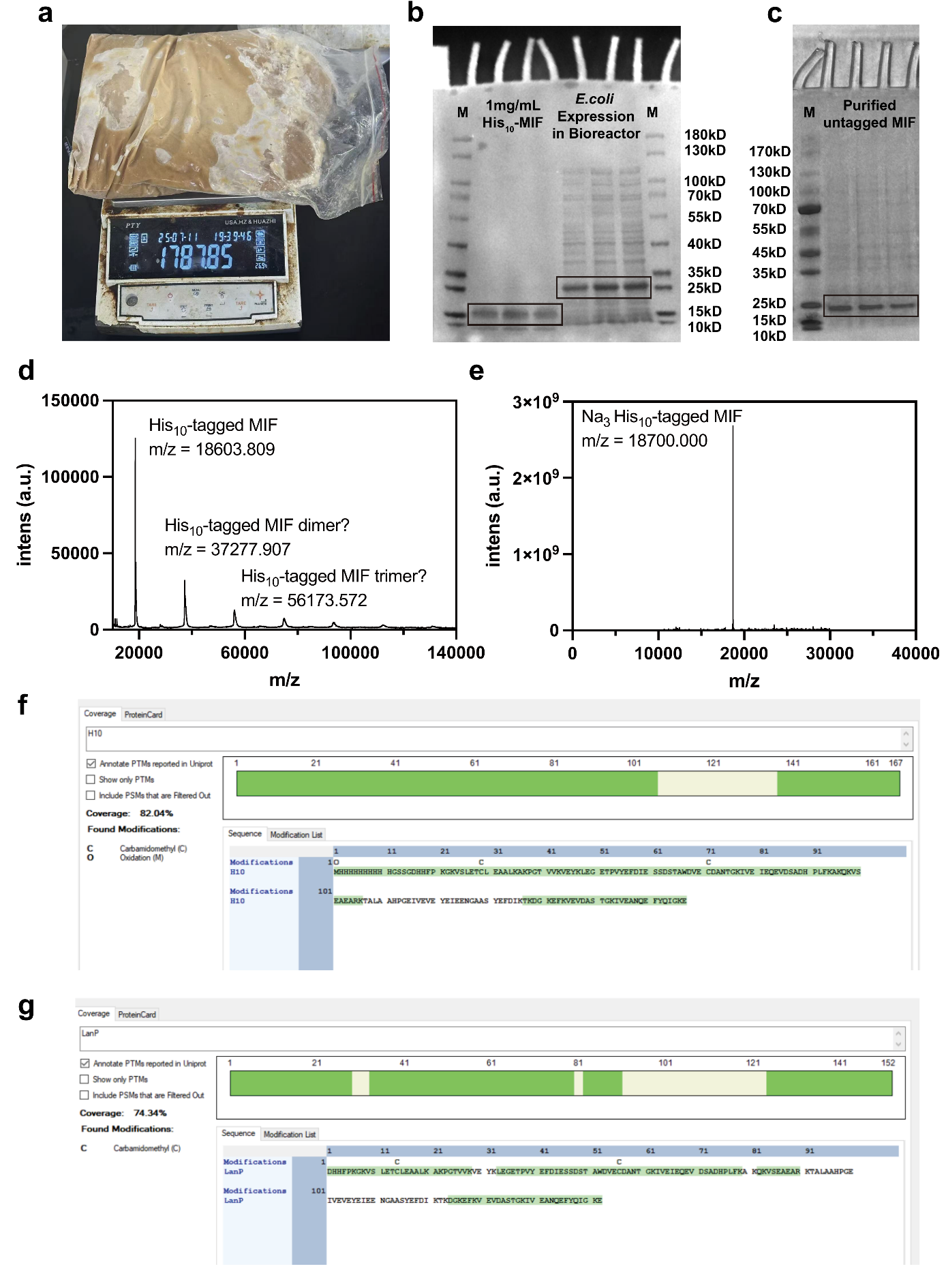

Fig. S1. Expression, purification, and characterization of His_10_-tagged lanpepsy (MIF, gene ID Mfla_0908) and untagged MIF
a, Photograph of the wet cell obtained after fermenting 25 L of *E. coli* BL21(DE3) expressing His_10_‑tagged MIF. The harvest yielded 1787.85 g of biomass. b, SDS-PAGE analysis of His_10_-tagged MIF expressed in a bioreactor. Lane M, molecular-weight marker. Grayscale intensity was used to estimate protein yield by ImageJ. c, SDS-PAGE analysis of purified untagged MIF, where a single sharp band with no detectable contaminants indicates high product purity. d, Matrix-Assisted Laser Desorption/Ionization Time-of-Flight Mass (MALDI-TOF) spectrometry analysis of purified His_10_-tagged MIF shows a major peak at m/z = 18,603.809 corresponding to the monomeric form, with additional peaks at m/z = 37,277.907 and 56,173.572 consistent with potential dimeric and trimeric species, respectively. e, Quadrupole Time-of-Flight (Q-TOF) mass spectrometry of His_10_-tagged MIF shows a peak at m/z = 18,700.000, corresponding to the trisodium-adducted form of MIF in solution. f-g, Proteomic validation of the purified His_10_-tagged MIF (f) and untagged MIF (g) showing high sequence coverage, confirming the expected identity of the protein.

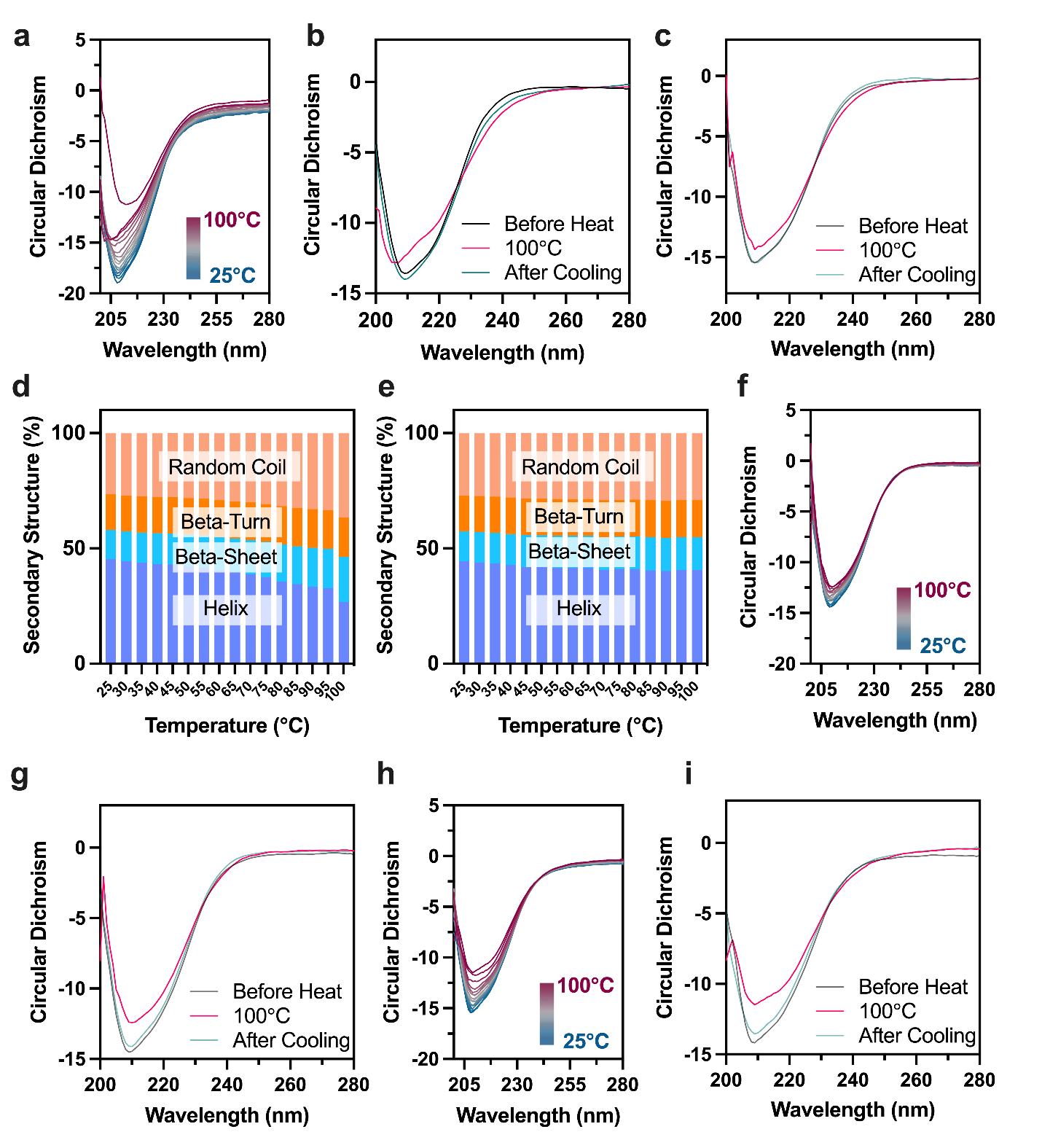

Supplementary Fig. S2. Variable-temperature circular dichroism (VT-CD) spectroscopy of Apo- and REE-bound MIF
a, VT-CD spectra of apo-MIF recorded from 25°C to 100°C; b, CD spectra of apo-MIF at 25°C (Before Heat), 100°C, and after cooling back to 25°C; c, CD spectra of La^III^-MIF at 25°C (Before Heat), 100°C, and after cooling; d, Secondary structure content of apo-MIF as a function of temperature, derived from CD spectral deconvolution; e, Secondary structure content of La^III^-MIF at varying temperatures; f, VT-CD spectra of Sm^III^-MIF; g, CD spectra of Sm^III^-MIF before heating (Before Heat), at 100°C, and after cooling; h, VT-CD spectra of Yb^III^-MIF; i, CD spectra of Yb^III^-MIF before heating, at 100°C, and after cooling. These VT‑CD data demonstrate that apo‑MIF is intrinsically heat‑stable and becomes even more thermoresistant upon binding REEs. Throughout a heating-cooling cycle from 25 °C to 100 °C and back to 25°C, the CD curves of MIF before heat and after cooling are essentially identical, underscoring the protein’s exceptional structural stability.

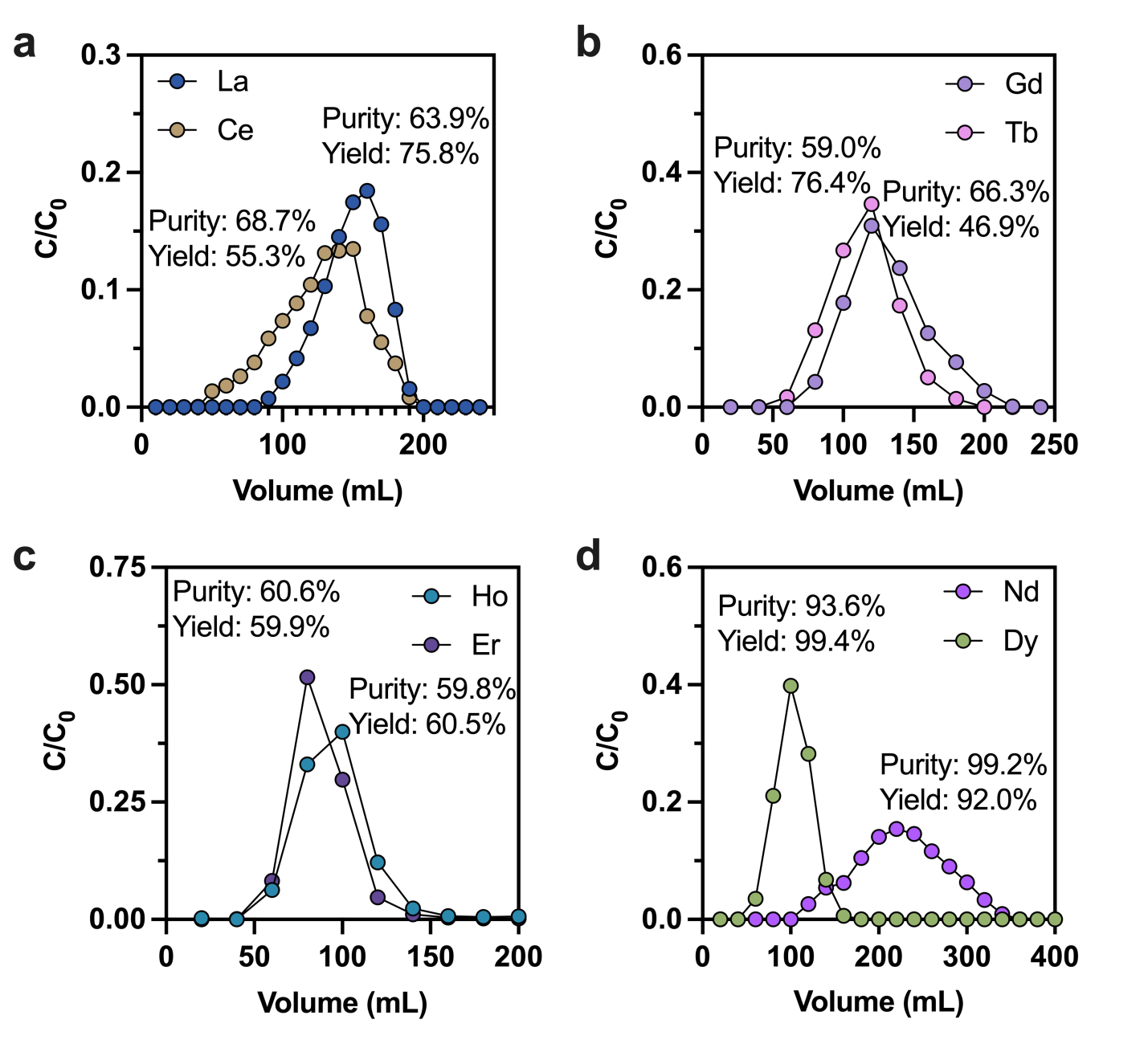

Supplementary Fig. S3. Separation performance of dimer lanmodulin (DLanM) as control experiments
a, Separation of La^III^-Ce^III^ (light REEs); b, Separation of Gd^III^-Tb^III^ (middle REEs); c, Separation of Ho^III^-Er^III^ (heavy REEs); d, Separation of a light-middle REE pair, Nd^III^-Dy^III^.These control experiments demonstrate that lanmodulin ligands cannot discriminate adjacent lanthanide pairs. Thus, MIF provides the first ligand capable of single-stage, aqueous-phase separation of all adjacent lanthanide pairs.

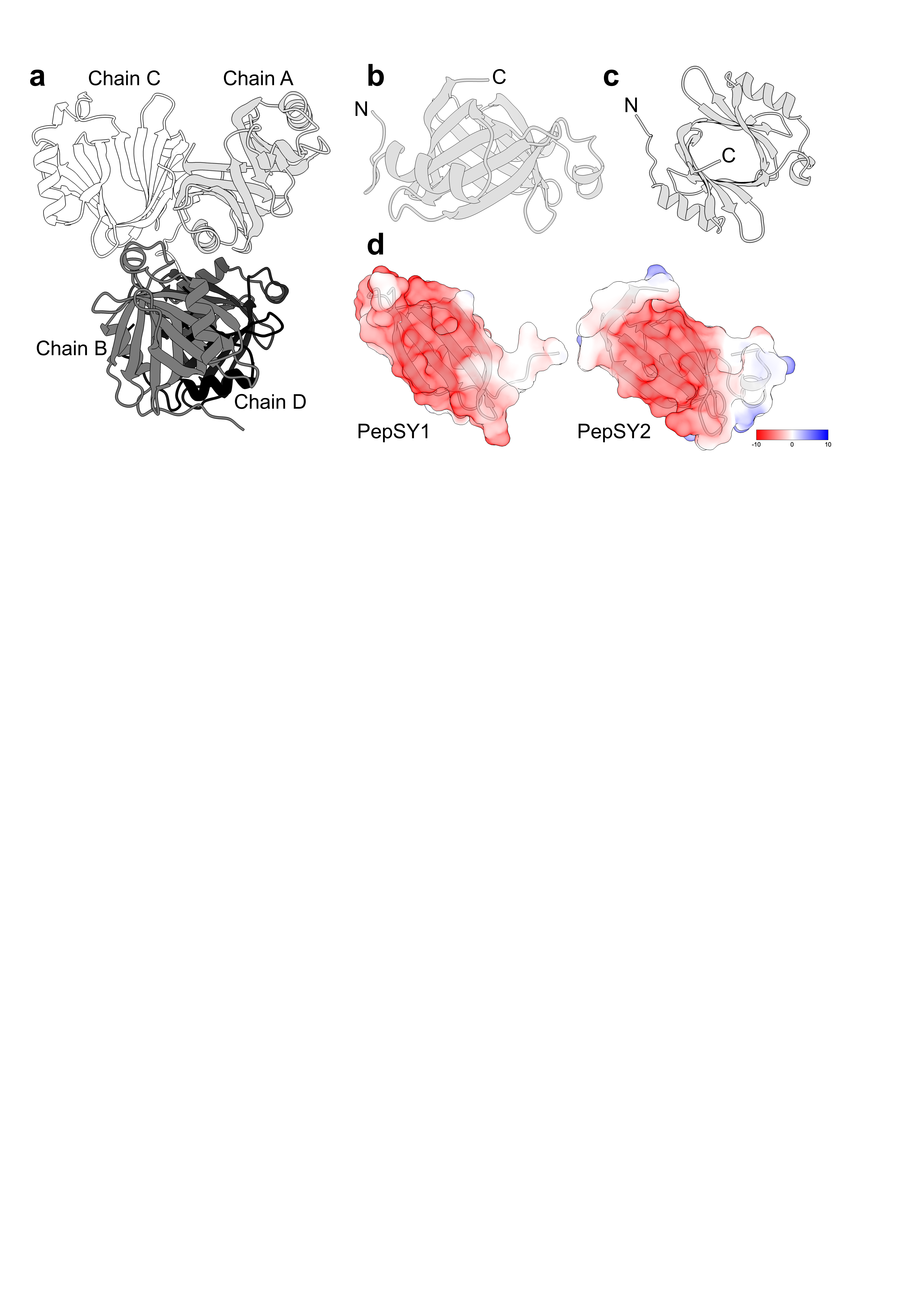

Supplementary Fig. S4. X-ray crystal structure of Apo-MIF, solved at 2.06 Å resolution
a, The asymmetric unit comprises four apo-MIF chains (Chains A–D); b,c, The structure of apo-MIF Chain A viewed from the side (b) and the top (c), revealing a pseudo-C2 symmetric architecture consisting of two PepSY domains.; d, Electrostatic surface potential maps of the PepSY1 and PepSY2 domains, highlighting the central electronegative β-barrel region. Electrostatic potentials were calculated using ChimeraX (*33*) and are shown in units of kcal·mol^-1^·e^-1^.

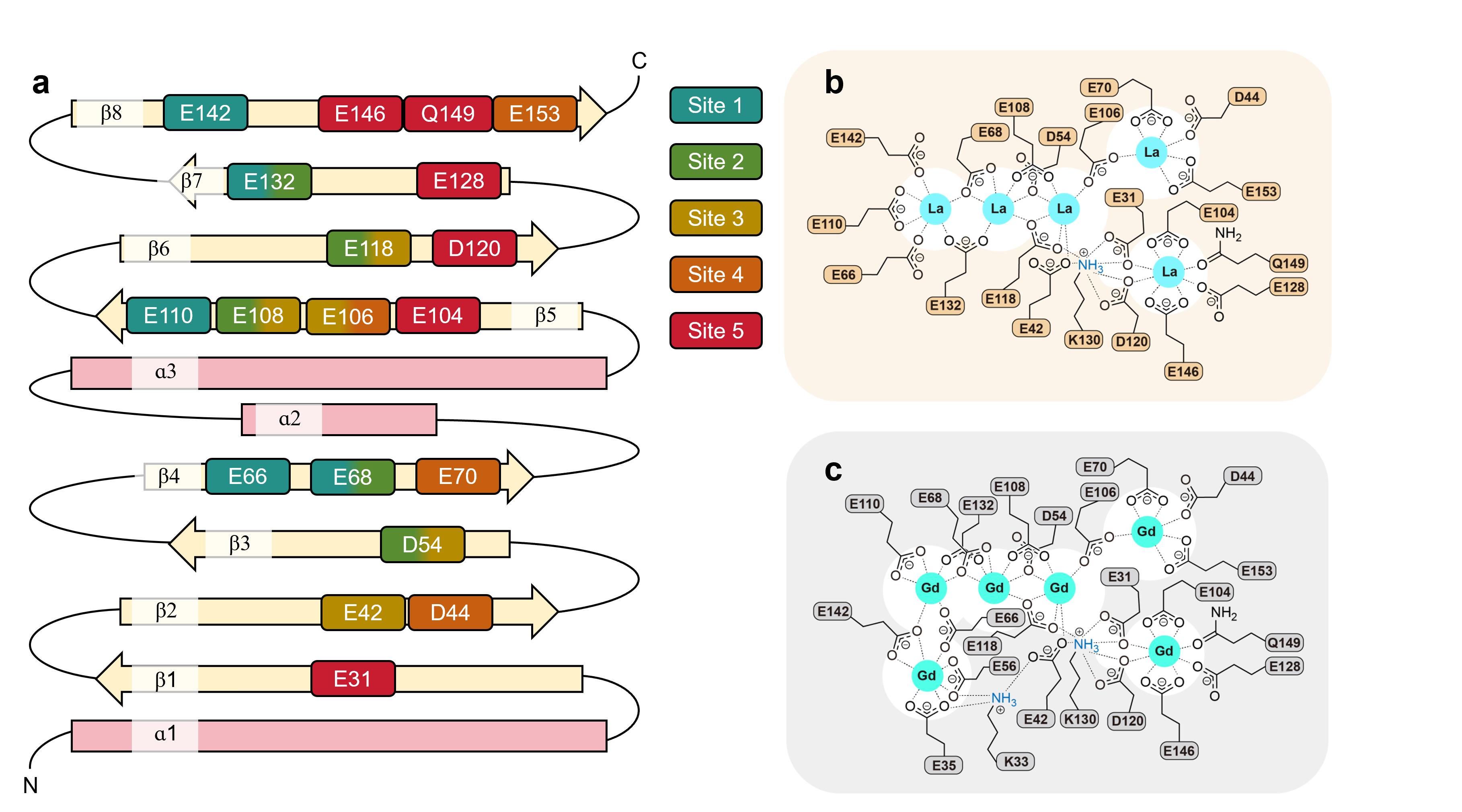

Supplementary Fig. S5. Topological diagram and coordination microenvironment of metal-binding residues in MIF
a, Topological diagram of the MIF, highlighting the distribution of key metal-coordinating residues across the secondary structure. The protein contains eight antiparallel β-strands, designated as β1-β8 from the N-terminus to the C-terminus, respectively. b-c, Coordination microenvironments of La^III^-MIF (b) and Gd^III^-MIF (c). In La^III^-MIF, 19 residues, including 15 glutamate (E), 3 aspartate (D), and 1 glutamine (Q), coordinate five La^III^ ions. In contrast, the Gd^III^-MIF employed two additional glutamate residues, E35 and E56, to coordinate the additional Gd^III^ ion at Site 1’, resulting in a total of 21 residues coordinating six Gd^III^ ions. Despite the difference in the number of binding sites, both La^III^- and Gd^III^-MIF complexes share a highly conserved set of bridging coordination residues, including D54, E68, E106, E108, E118, and E132, which support the densely metal stacking structure. In both structures of La^III^- and Gd^III^-MIF, K130 form hydrogen bonds with multiple acidic E and D residues by ε-amino groups and provide additional stabilization. The key difference between light and middle REE binding modes in Gd^III^-MIF lies in the presence of extra bridging residues—E66 and E142, bridging Site 1 and Site 1’ via bidentate coordination. Yet in La^III^-MIF, they act as monodentate ligands in Site 1.

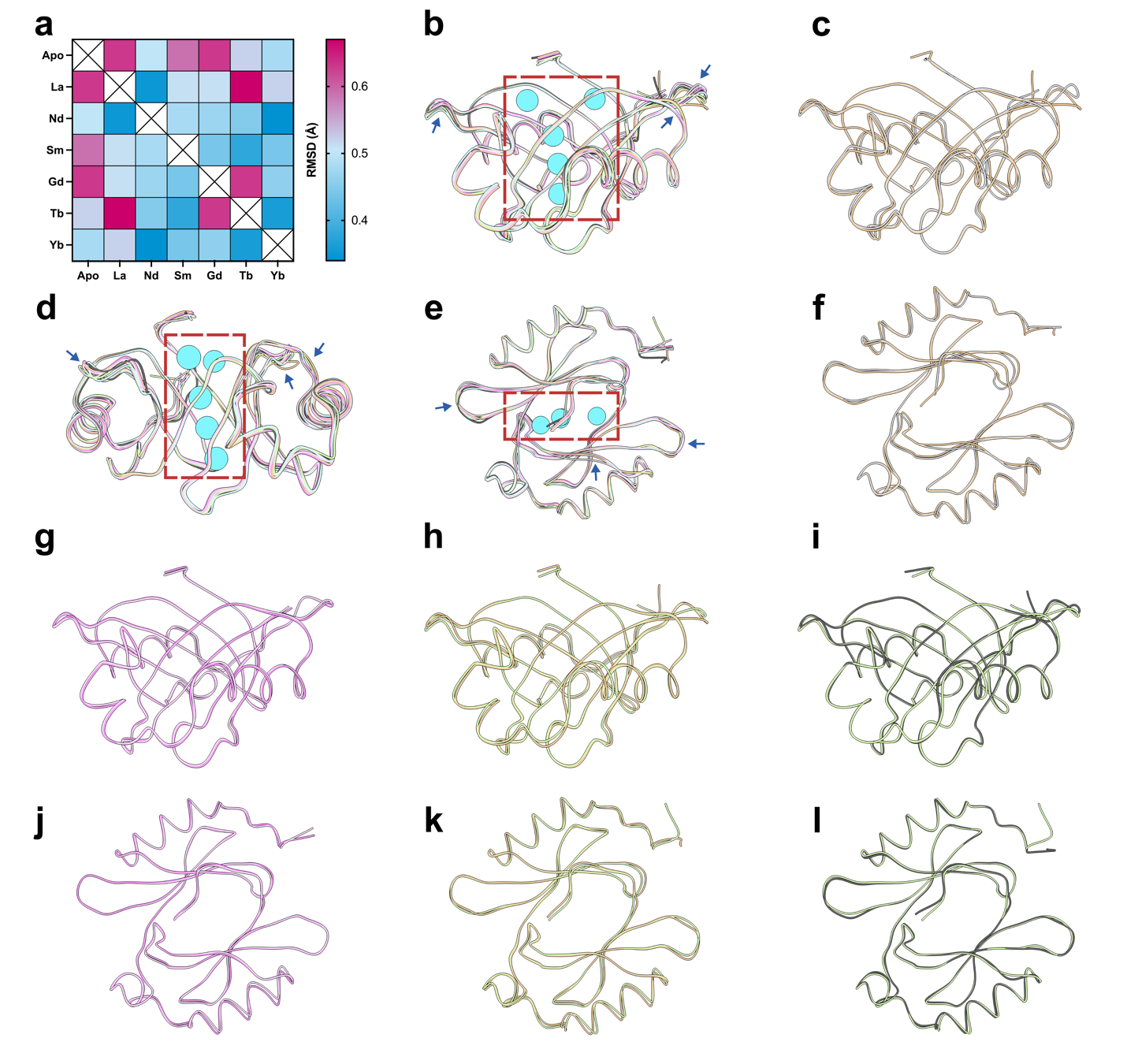

Supplementary Fig. S6. Structural comparison of apo- and REE-bound MIF complexes
a, Pairwise global Cα root mean square deviation (RMSD) heatmap among the six REE-MIF (La^III^, Nd^III^, Sm^III^, Gd^III^, Tb^III^, and Yb^III^) and apo‑MIF structure, the scale spans from 0.34 Å in cyan to 0.67 Å in magenta. Light (La^III^) and middle REE (Sm^III^) complexes deviate heavily from the apo structure, with an RMSD of approximately 0.6 Å. In contrast, the heavy REE complex (Yb^III^-MIF) is similar to apo-MIF, with an RMSD of 0.48 Å. White squares on the diagonal are the result of self‑comparisons. b, Superimposition of all MIF backbone structures in front view, with the five La^III^-binding sites indicated by cyan spheres (based on La^III^-MIF) for the conserved REE coordination microenvironment. The backbones are colored as follows: La^III^, coral yellow; Nd^III^, pink violet; Sm^III^, pale sky blue; Gd^III^, charcoal grey; Tb^III^, mint green; Yb^III^, light lavender; apo, silver grey. c, Comparison of La^III^-MIF (coral yellow) and apo-MIF (silver grey) in front view. d-e, Superimposition of all MIF backbone structures in side view (d) and top view (e). f, Top view of apo- and La^III^-MIF comparison (panel c); g, Nd^III^-MIF (pink violet) vs. Yb^III^–MIF (light lavender) in front view; h, La^III^-MIF (coral yellow) vs. Tb^III^-MIF (mint green) in front view; i, Gd^III^-MIF (charcoal grey) vs. Tb^III^-MIF (mint green) in front view; j–l, Corresponding top views for panels g–i. Superimposition of all MIF backbones shows a conserved coordination microenvironment (red dashed box) accompanied by backbone deformations (blue arrows) that accommodate different REE ions. These comparisons show that MIF amplifies picometre‑scale differences in REE ionic radii into angstrom-level RMSD differences between REE-MIF complexes.

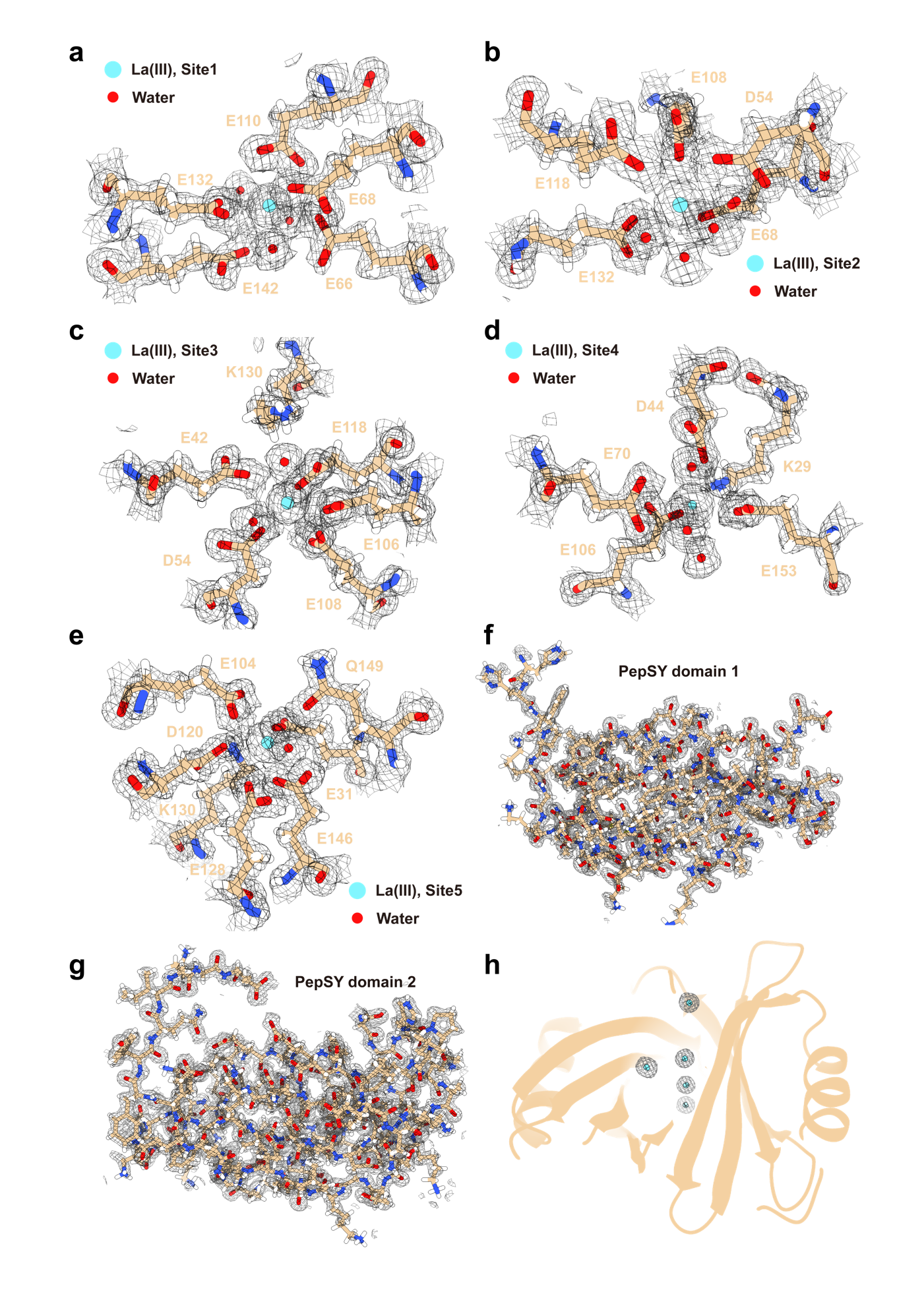

Supplementary Fig. S7. X-ray crystal structure of La^III^-MIF, solved at 1.25 Å resolution
a-e, Metal coordination microenvironments in the five binding sites (Sites 1-5) of the La^III^-MIF complex are shown with 2F_o_–F_c_ electron density maps contoured at 1.0 σ. La^III^ ions (blue spheres) and coordinating residues, including side chains and water molecules (red spheres), are indicated. The specific residues involved in metal coordination at each site are labeled. f-g, The overall electron density maps and structural models of PepSY1 (f) and PepSY2 (g) within the La^III^-MIF complex. h, The cartoon representation of the MIF scaffold with La^III^ ion coordination.

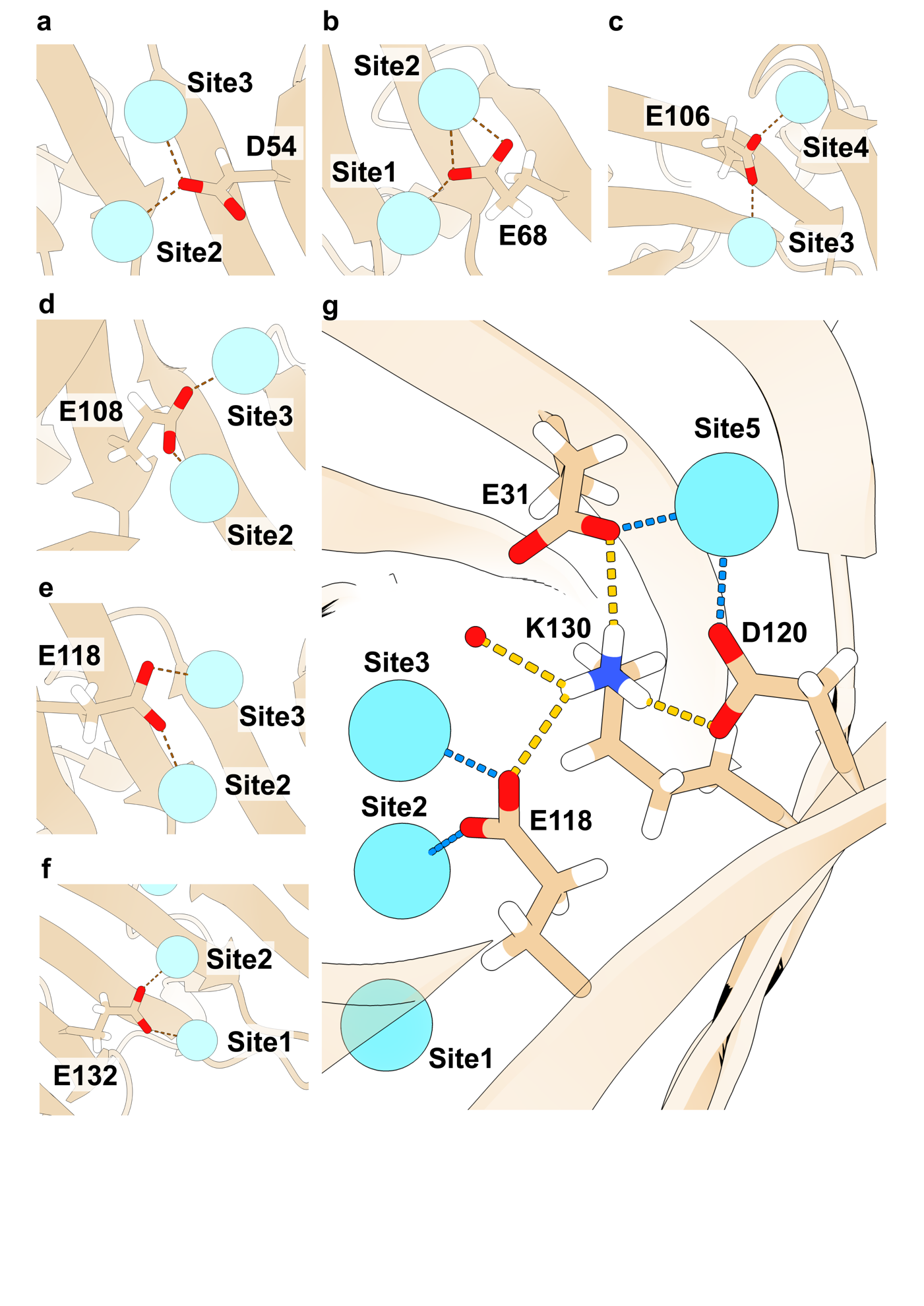

Supplementary Fig. S8. Detail of glutamate (E), aspartate (D), and lysine (K) that stabilize the densely stacked La^III^ ions in MIF
a-f. a. D54, b. E68, c. E106, d, E108, e, E118, f. E132. Each residue bridges two neighboring La^III^ ions (cyan spheres) via the oxygen atoms (red) of its carboxylate group, as indicated by dashed lines. These bridging residues stabilize MIF’s densely stacked architecture, allowing La^III^ ions to stack at a close distance of 4.3 Å between La^III^ ions. g. K130. The ε-amino group (blue, representing the nitrogen atom) of K130 forms hydrogen bonds (yellow dashed lines) with several carboxylate ligands that interact with La^III^, thereby further stabilising the densely stacked La^III^ ion in MIF.

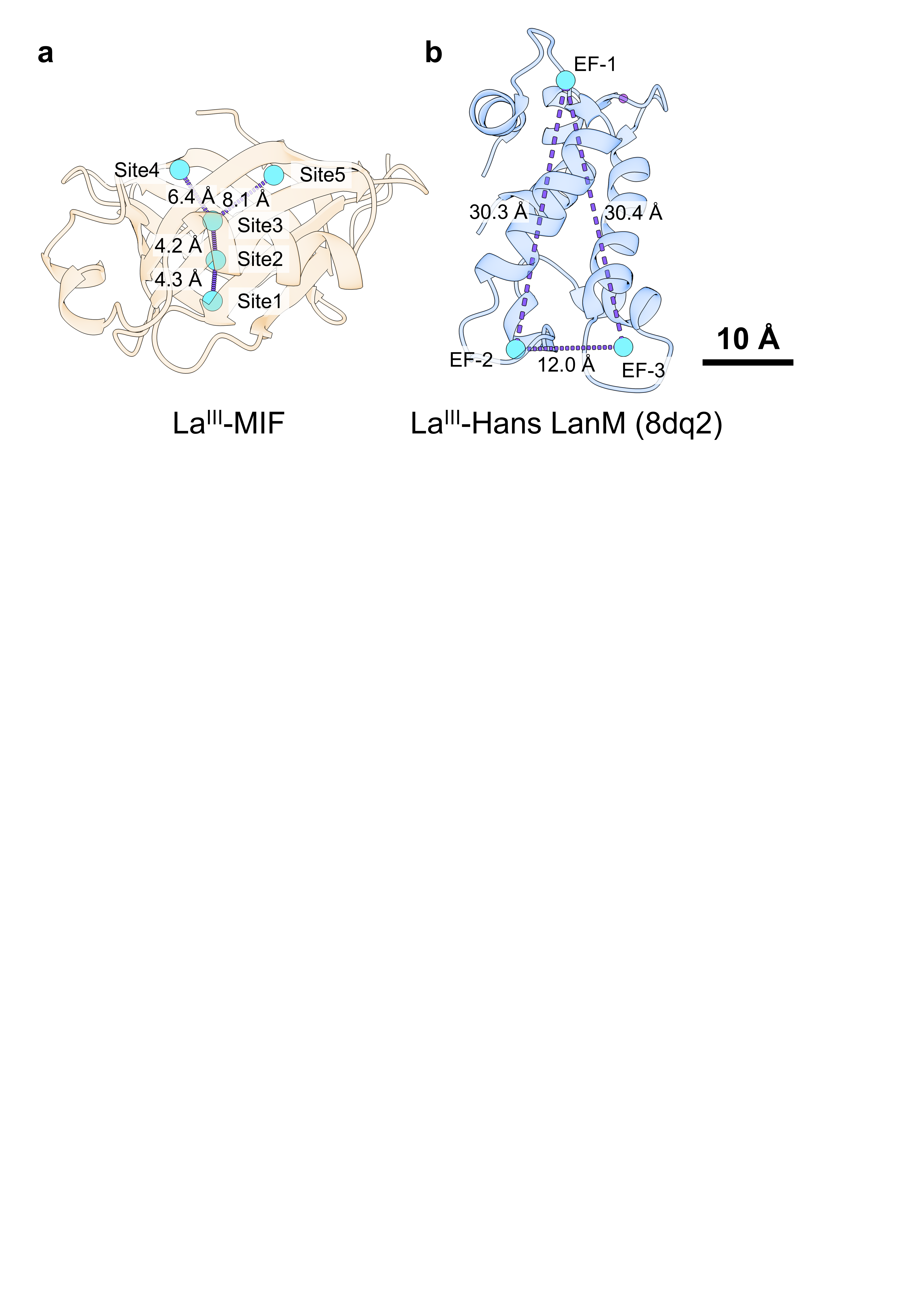

Supplementary Fig. S9. Structural characteristic of La^III^-MIF in comparison with La^III^-Lanmodulin (Hans LanM, PDB ID: 8DQ2)
Structure difference is observed in La^III^-MIF and La^III^-lanmodulin in terms of metal-binding architectures. MIF displays five densely stacked La^III^-binding sites within its coordination microenvironment, with metal-metal distances of 4.2-8.1 Å and bridging coordination mediated by acidic residues. In contrast, Lanmodulin contains four dispersed EF-hands, three of which are La^III^-binding sites, with ion-ion distances ranging from 12 to 30 Å and without inter-site bridging interaction. This structural difference underpins the unique stacking-based amplification mechanism of MIF, which differs from all known lanthanide-binding proteins reported to date.

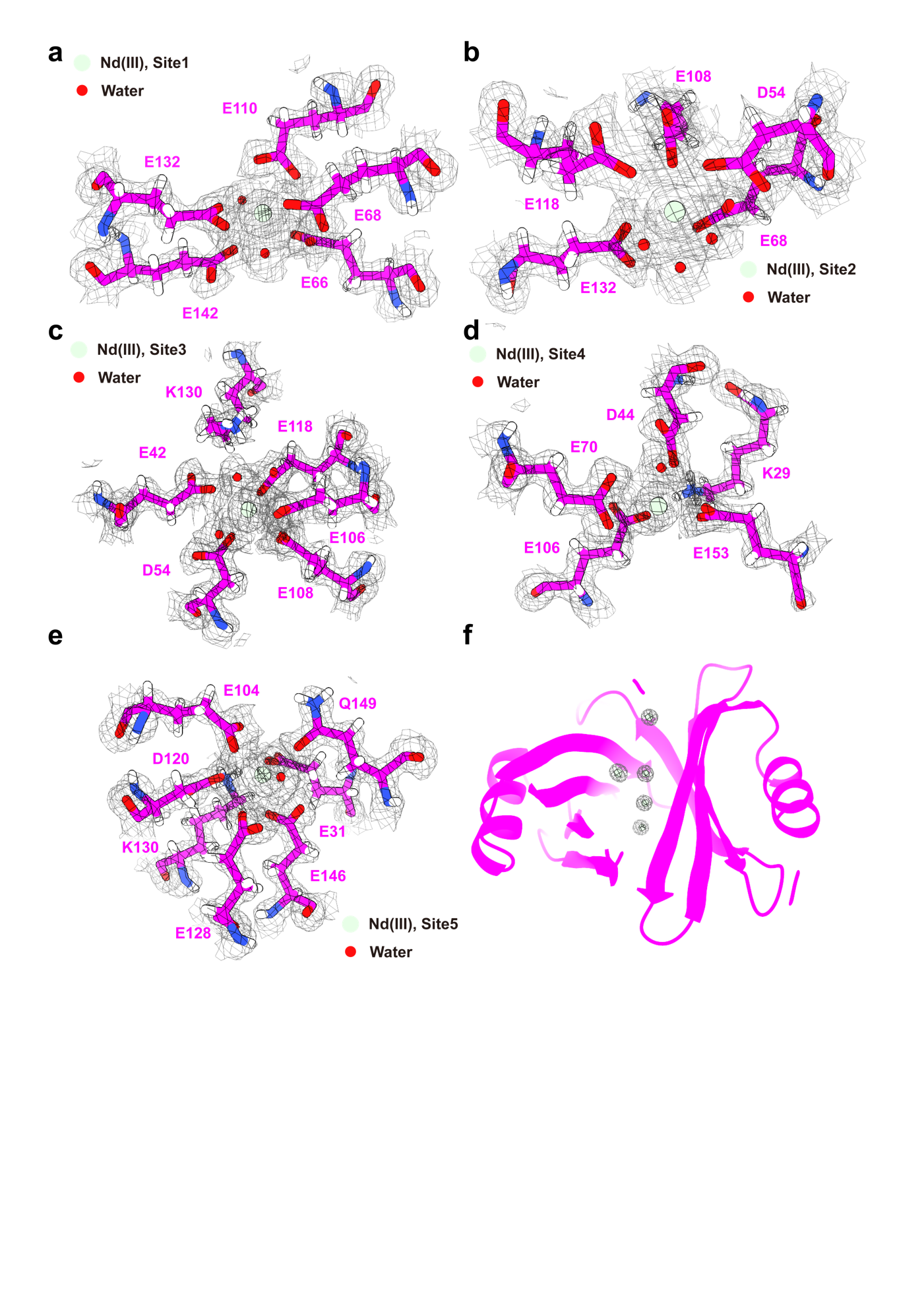

Supplementary Fig. S10. X-ray crystal structure of Nd^III^-MIF, solved at 1.41 Å resolution
a-e, Metal coordination microenvironments in the five binding sites (Sites 1-5) of the Nd^III^-MIF complex are shown with 2F_o_–F_c_ electron density maps contoured at 1.0 σ. Nd^III^ ions (pale green spheres) and coordinating residues, including side chains and water molecules (red spheres), are indicated. The specific residues involved in metal coordination at each site are labeled. f, The cartoon representation of the MIF scaffold with Nd^III^ ion coordination.

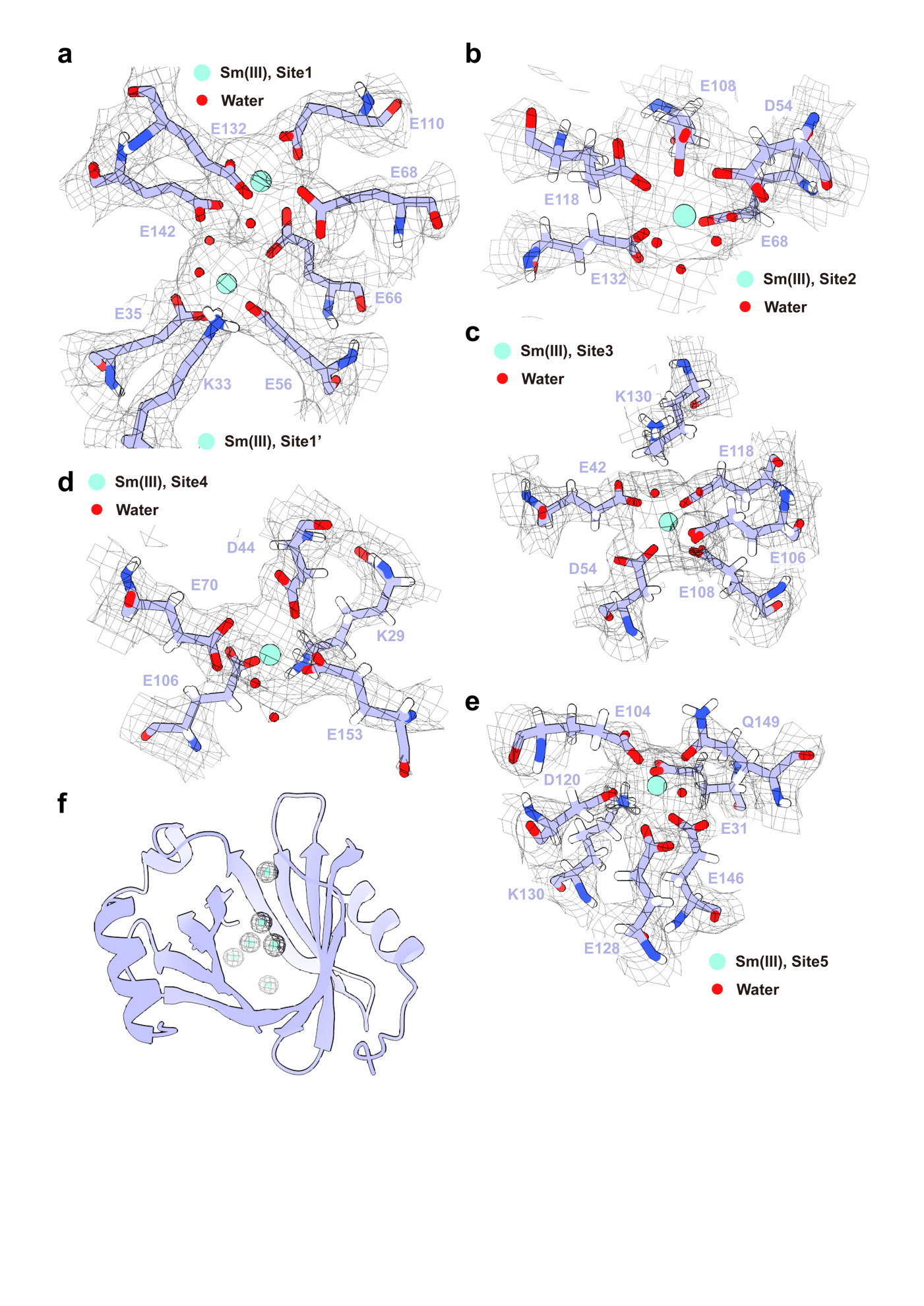

Supplementary Fig. S11. X-ray crystal structure of Sm^III^-MIF, solved at 2.04 Å resolution
a-e, Metal coordination microenvironments in the six binding sites (Sites 1-5, and Site 1’) of the Sm^III^-MIF complex are shown with 2F_o_–F_c_ electron density maps contoured at 1.0 σ. Sm^III^ ions (light aqua spheres) and coordinating residues, including side chains and water molecules (red spheres), are indicated. The specific residues involved in metal coordination at each site are labeled. f, the cartoon representation of the MIF scaffold with Sm^III^ ions coordination.

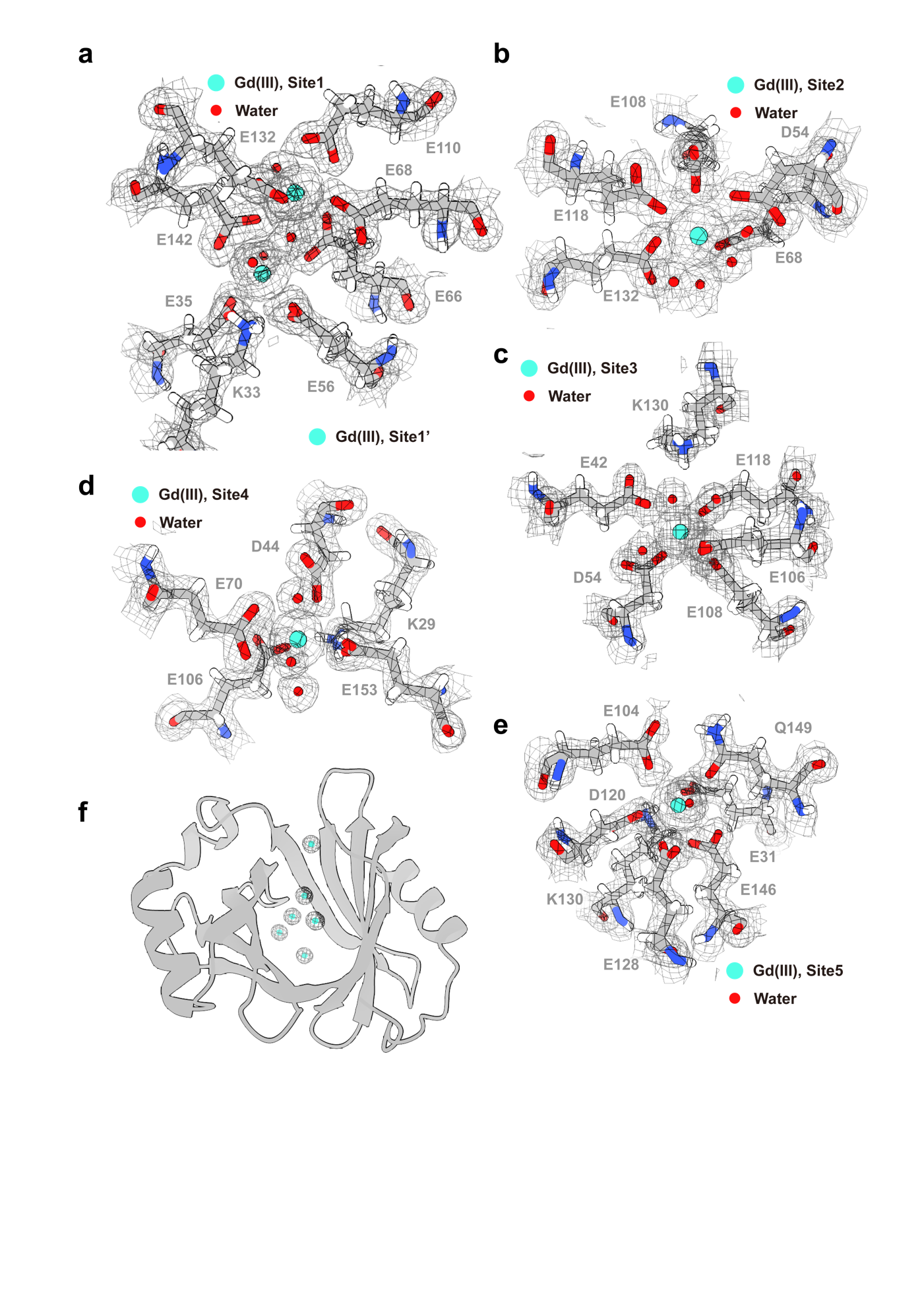

Supplementary Fig. S12. X-ray crystal structure of Gd^III^- MIF, solved at 1.41 Å resolution
a-e, Metal coordination microenvironments in the six binding sites (Sites 1-5, and Site1’) of the Gd^III^-MIF complex are shown with 2F_o_–F_c_ electron density maps contoured at 1.0 σ. Gd^III^ ions (cyan spheres) and coordinating residues, including side chains and water molecules (red spheres), are indicated. The specific residues involved in metal coordination at each site are labeled. f, The cartoon representation of the MIF scaffold with Gd^III^ ions coordination.

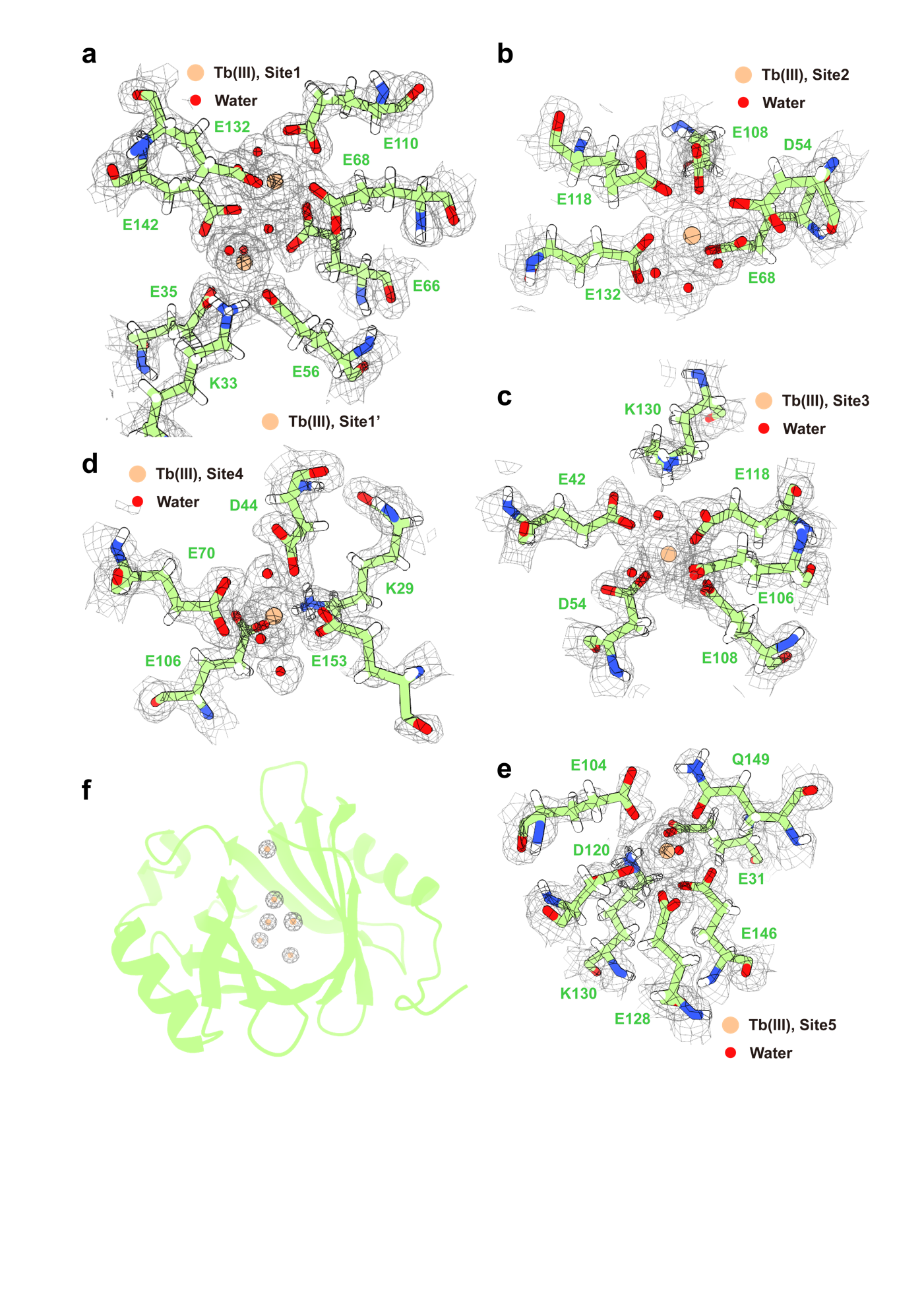

Supplementary Fig. S13. X-ray crystal structure of Tb^III^- MIF, solved at 1.55 Å resolution
a-e, Metal coordination microenvironments in the six binding sites (Sites1-5, and Site1’) of the Tb^III^-MIF complex are shown with 2F_o_–F_c_ electron density maps contoured at 1.0 σ. Tb^III^ ions (coral peach spheres) and coordinating residues, including side chains and water molecules (red spheres), are indicated. The specific residues involved in metal coordination at each site are labeled. f, The cartoon representation of the MIF scaffold with Tb^III^ ions coordination.

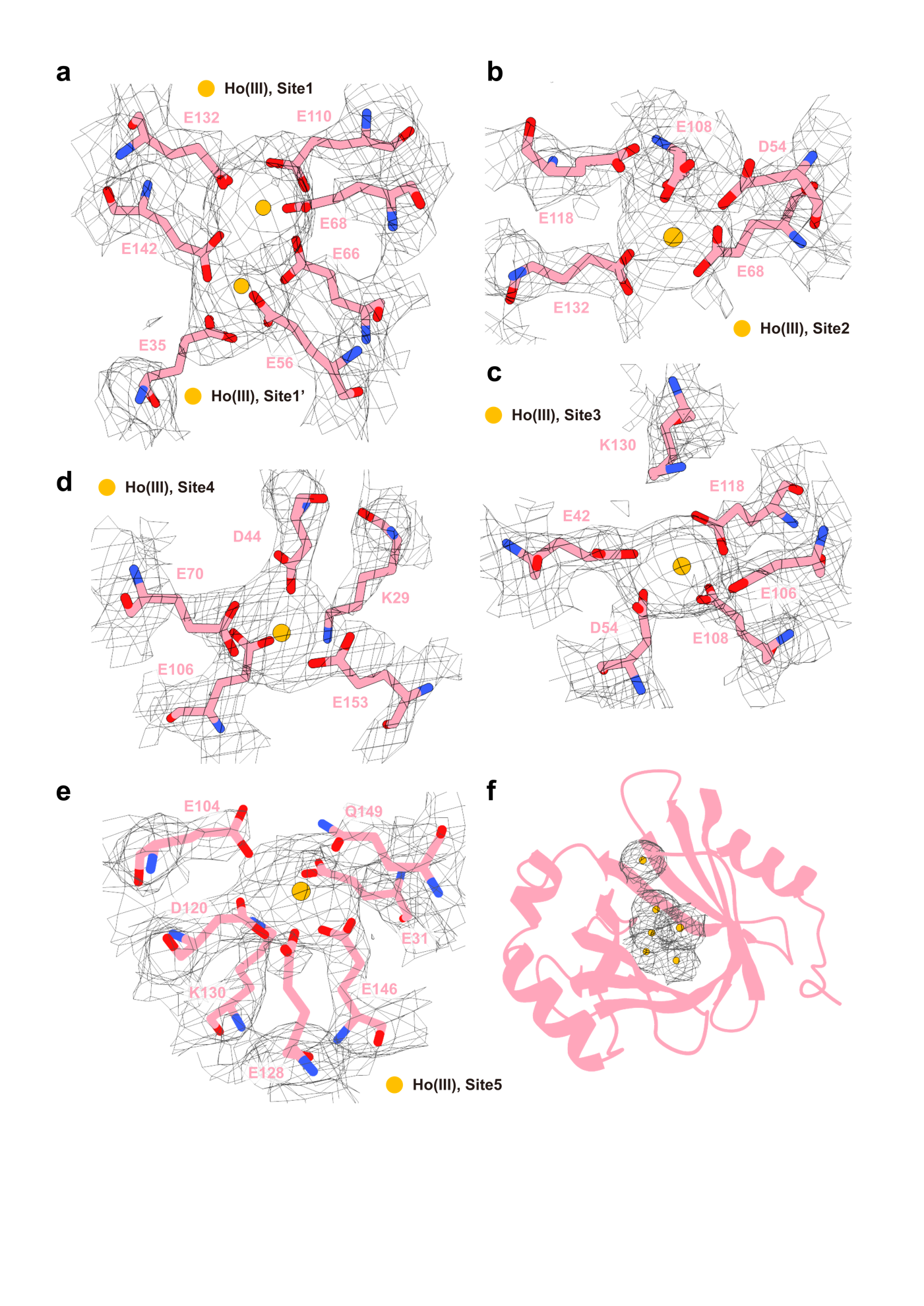

Supplementary Fig. S14. X-ray crystal structure of Ho^III^-MIF, solved at 3.05 Å resolution
a-e, Metal coordination microenvironments in the six binding sites (Sites 1-5, and Site 1’) of the Ho^III^-MIF complex are shown with 2F_o_–F_c_ electron density maps contoured at 1.0 σ. Ho^III^ ions (amber yellow spheres) and coordinating residues are indicated. The specific residues involved in metal coordination at each site are labeled. f, The cartoon representation of the MIF scaffold with Ho^III^ ions coordination.

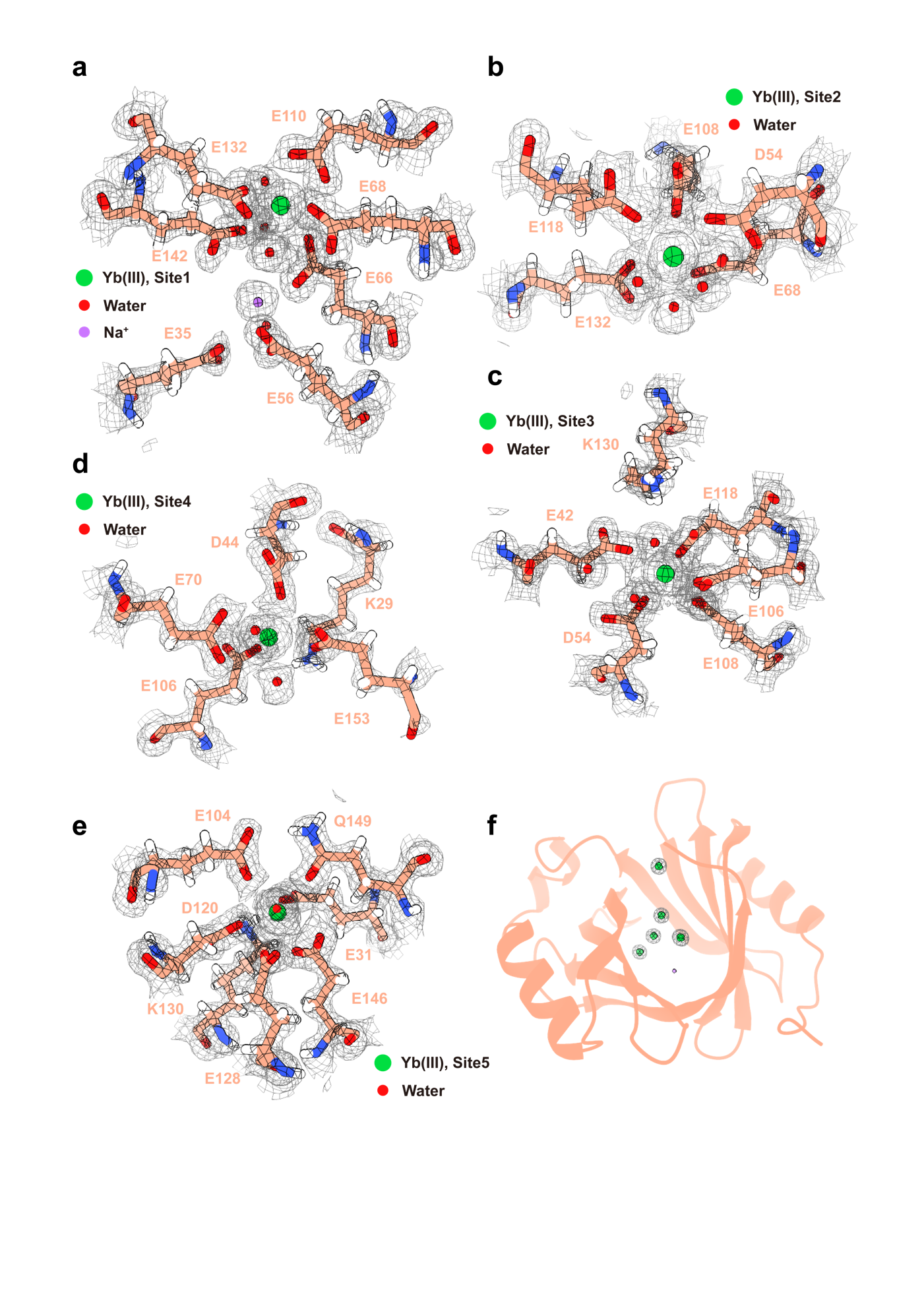

Supplementary Fig. S15. X-ray crystal structure of Yb^III^-MIF, solved at 1.21 Å resolution
a-e, Metal coordination microenvironments in the five REE binding sites (Sites 1-5) and a Na^I^ binding site (Site1’) of the Yb^III^-MIF complex are shown with 2F_o_–F_c_ electron density maps contoured at 1.0 σ. Yb^III^ ions (green spheres), Na^I^ ion (violet spheres) and coordinating residues, including side chains and water molecules (red spheres), are indicated. The specific residues involved in metal coordination at each site are labeled. f, The cartoon representation of the MIF scaffold with Yb^III^ ions coordination.

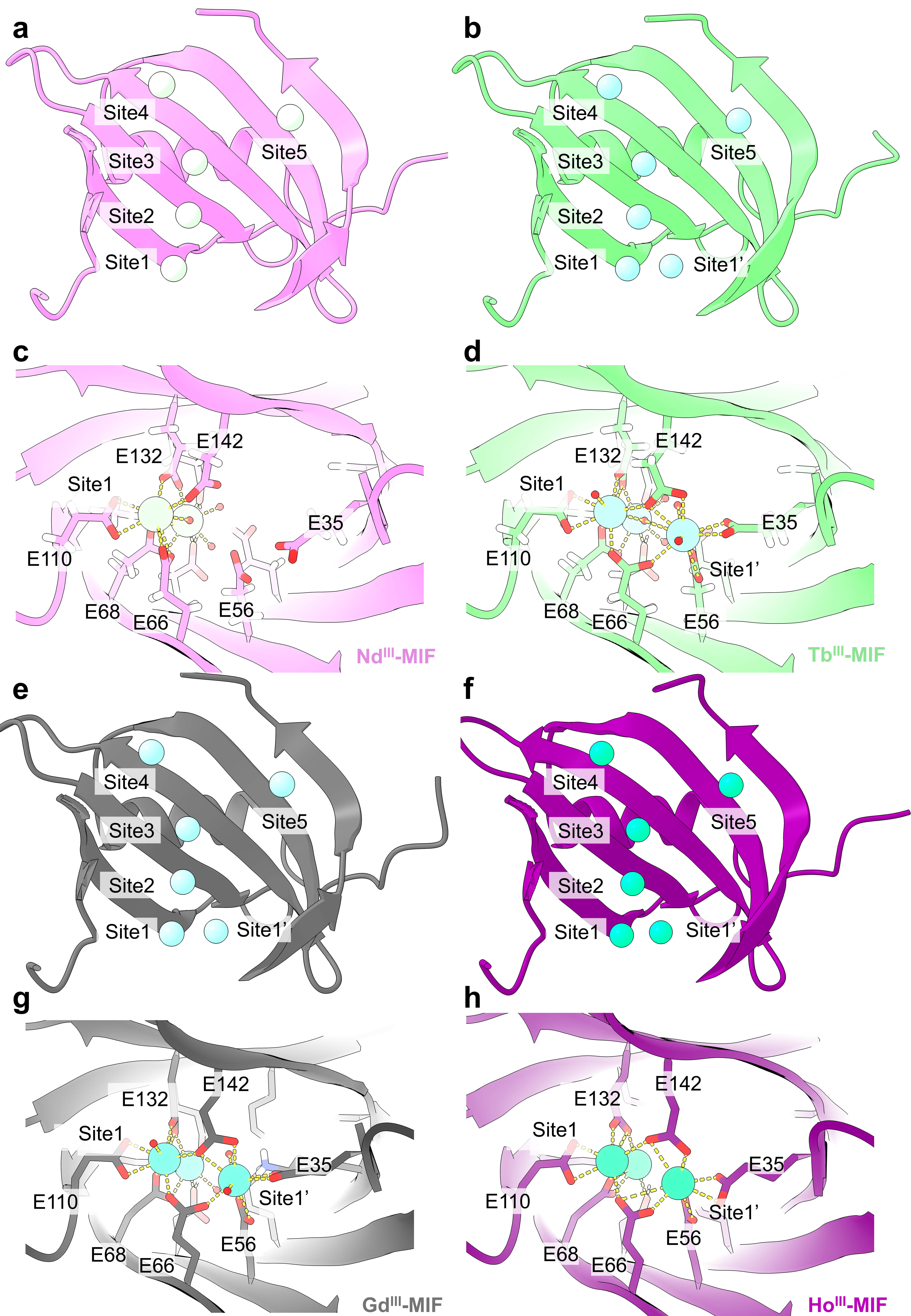

Supplementary Fig. S16. Structural details of Nd^III^-, Tb^III^-, Gd^III^-, and Ho^III^-MIF complexes
a-b, Nd^III^-MIF complex (pale green spheres for Nd^III^); c-d, Tb^III^-MIF complex (sky cyan spheres for Tb^III^); e-f, Gd^III^-MIF complex (bright cyan spheres for Gd^III^); g-h, Ho^III^-MIF complex (turquoise cyan spheres for Ho^III^). Panels a, c, e, and g show the side view of MIF. Panels b, d, f, and h present close-up, top views of MIF at Site 1 and Site 1’.

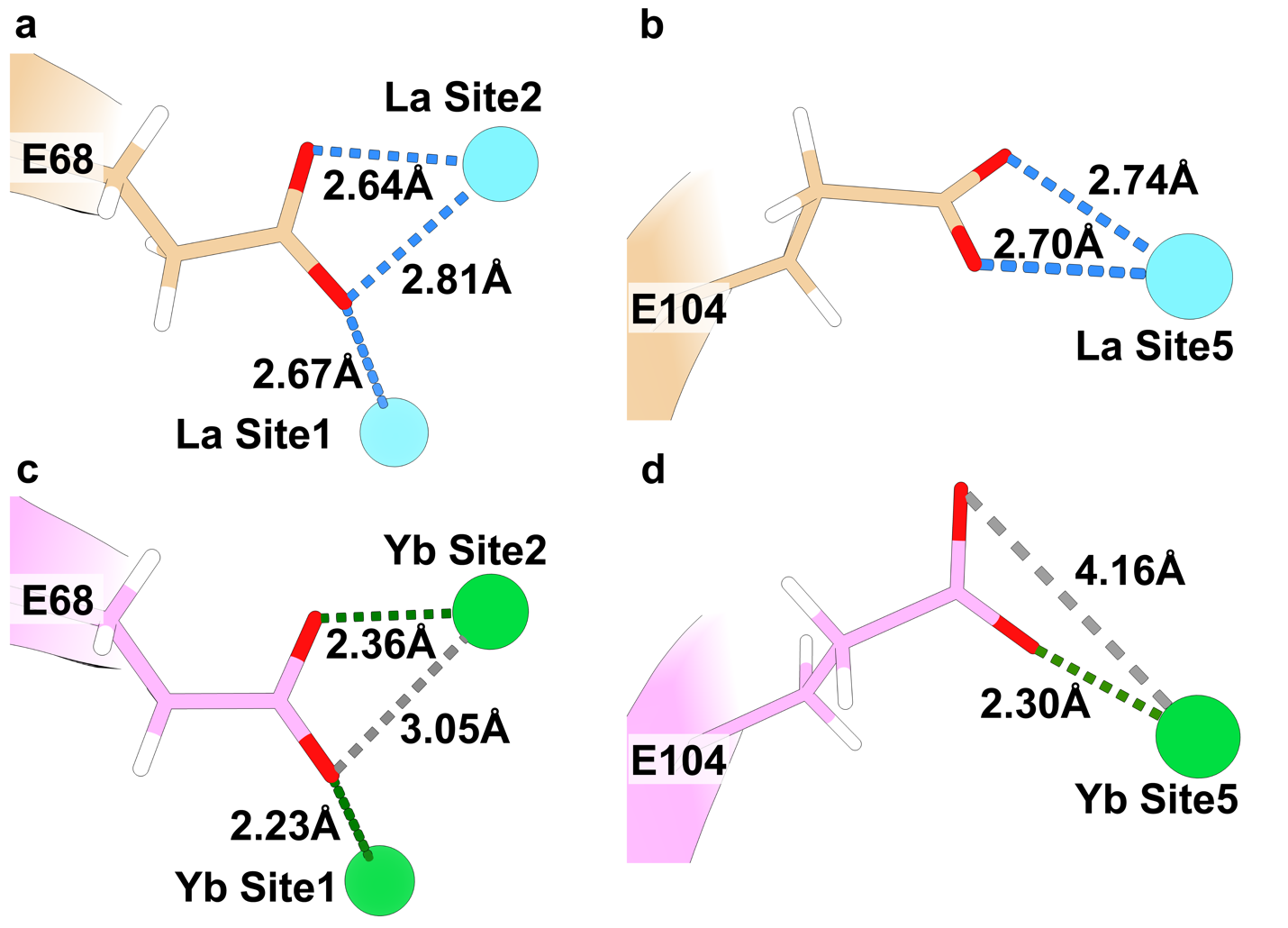

Supplementary Fig. S17. Coordination modes of E68 and E104 in La^III^‑MIF and Yb^III^‑MIF.
a, In the La^III^-MIF, E68 bridges Sites 1 and 2: its two carboxylate O atoms (red) interact with the La^III^ at Site 1 (2.64 and 2.81 Å, blue dashed lines) and one O atom coordinates the La^III^ at Site 2 (2.67 Å). b, In the same La^III^-MIF, E104 binds the La^III^ at Site 5 in a bidentate fashion (2.70 and 2.74 Å, blue dashed lines). c, In the Yb^III^-MIF, E68 still bridges Sites 1 and 2, but now with two shorter Yb-O contacts (2.23 and 2.36 Å, dark‑green dashed lines); the third Yb-O distance is 3.05 Å (grey dashed line) and is not considered a bond. d, E104 shifts to monodentate coordination with the Yb^III^ at Site 5 (2.30 Å, dark-green dashed line); the second O atom lies 4.16 Å away (grey dashed line) and is non‑bonding. All measured metal‑ligand or metal‑oxygen distances are labelled in Å.

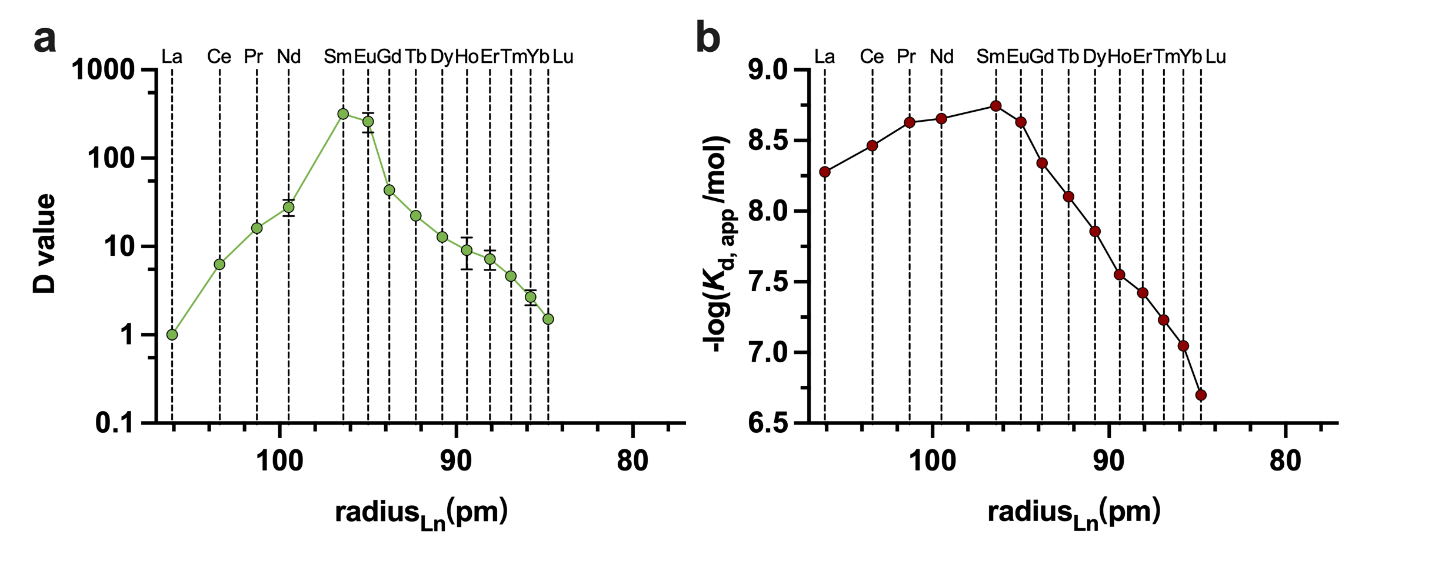

Supplementary Fig. S18. Lanthanide binding affinity trends to MIF
a, Experimental distribution values (D values) for MIF with each of the 14 lanthanide ions, determined by ICP-OES and plotted against their respective ionic radii. Distribution values collected in experiments of every adjacent pair were normalized as described in the method. The result shows a preference for the middle REE of MIF. Error bars represent the mean ± SD of three independent experiments (n = 3). Some error bars can't be seen because they're too small and are hidden by the circles that are plotted at the mean for the corresponding data. b, Apparent binding affinities between different REE and MIF determined by ITC. The results are expressed as the decadic negative logarithm of the apparent dissociation constant, here reported as the dimensionless quantity,  -log(*K*_d,app_/mol). Both datasets reveal a selectivity trend to middle lanthanides (Sm-Gd). The x-axis shows the ionic radii of trivalent lanthanides in picometers.

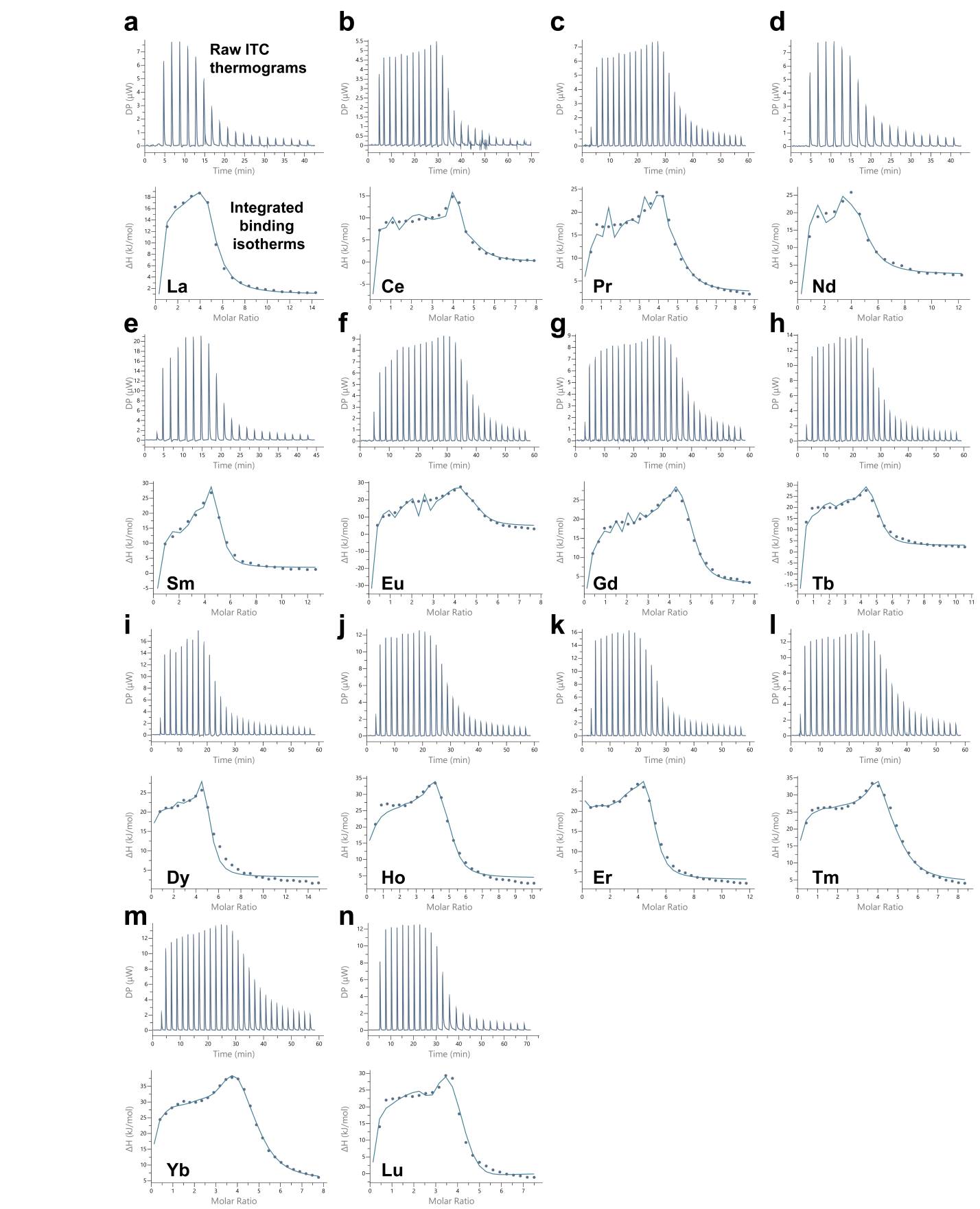

Supplementary Fig. S19. Isothermal titration calorimetry (ITC) analysis of MIF binding to 14 lanthanide ions
Raw ITC thermograms and integrated binding isotherms of MIF titrated with La^III^ (a), Ce^III^ (b), Pr^III^ (c), Nd^III^ (d), Sm^III^ (e), Eu^III^ (f), Gd^III^ (g), Tb^III^ (h), Dy^III^ (i), Ho^III^ (j), Er^III^ (k), Tm^III^ (l), Yb^III^ (m), and Lu^III^ (n), respectively. The binding data were fitted using a sequential binding model to derive thermodynamic parameters, including *K*_d_ of each site, and are shown in Fig. 3b. The distinct *K*_d_ values observed for each site of REE-MIFs highlight MIF’s selectivity toward different lanthanide ions.

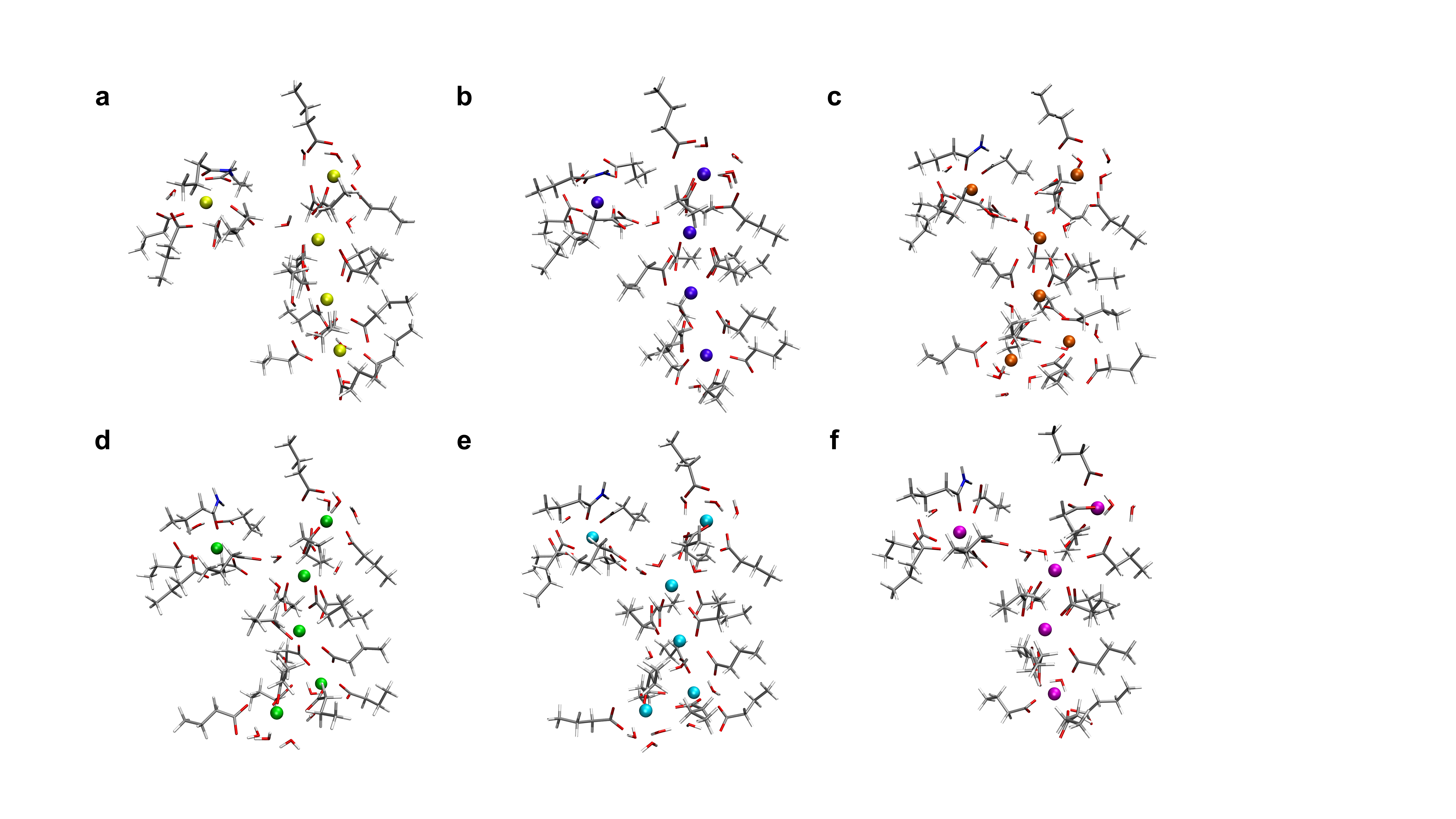

Supplementary Fig. S20. Cluster models extracted from the molecular-dynamics trajectories of the REE–MIF complexes at the 3-μs. a, La^III^, b, Nd^III^, c, Sm^III^, d, Gd^III^, e, Tb^III^, f, Yb^III^. The models were employed to investigate the binding energy between MIF and REE ions.

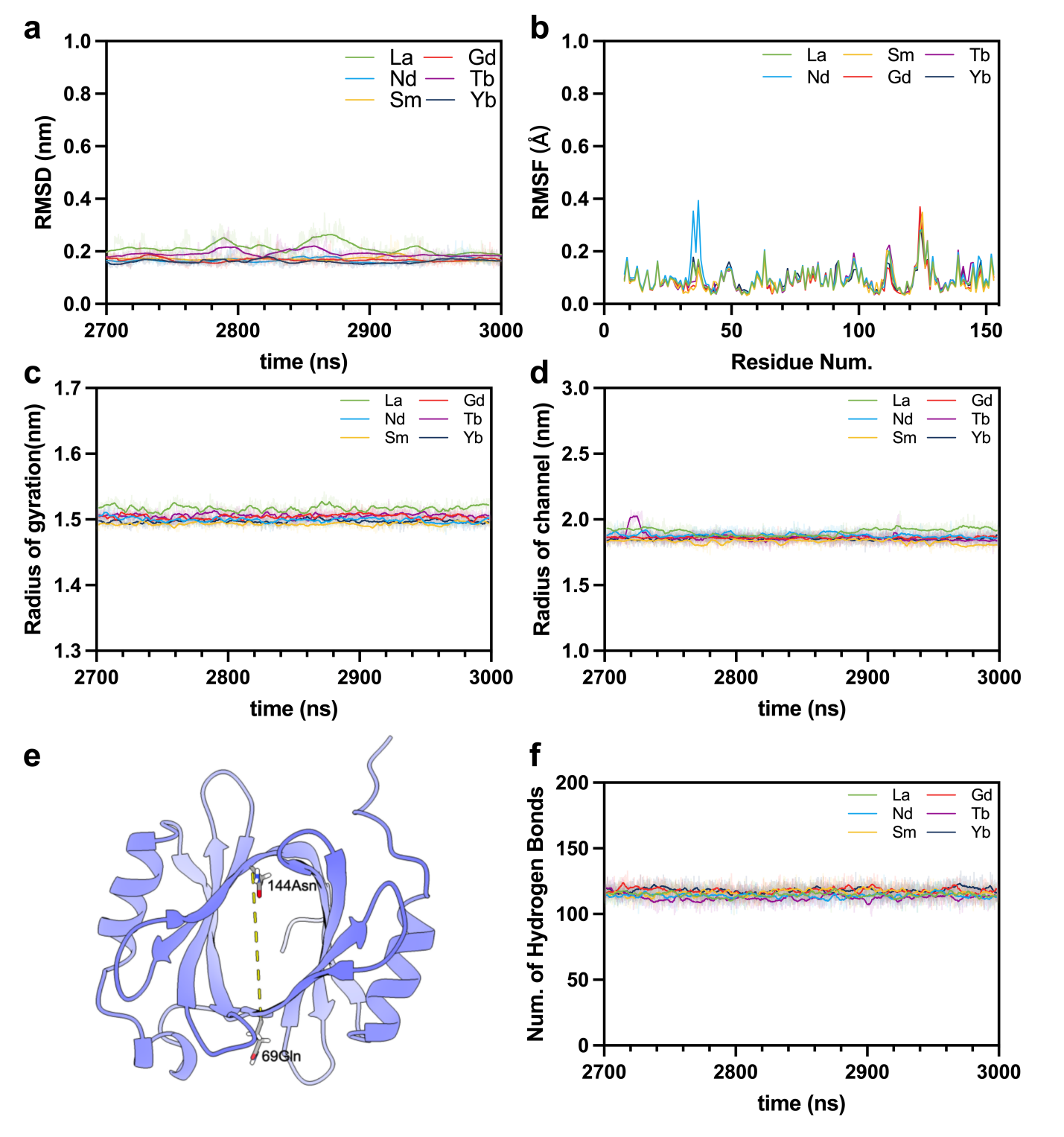

Supplementary Fig. S21. Molecular dynamics (MD) simulations reveal the structural stability of REE–MIF complexes at equilibrium.
a, RMSD of all MIF backbone Cα atoms over 2700–3000 ns for six REE–MIF complexes, relative to their initial conformations. The results exhibit small fluctuations and flat trends, indicating that all systems have reached dynamic equilibrium.b, Root mean square fluctuation (RMSF) of Cα atoms during the 2700–3000 ns simulation window. The results indicate minimal backbone flexibility across all REE–MIF complexes, confirming the overall rigidity of the β-barrel scaffold. c, The radius of gyration values remain stable throughout the 2700–3000 ns simulation window for each REE-MIF complex, further supporting that the systems have reached dynamic equilibrium. d, The radius of the central metal coordination channel, defined by the Cα–Cα distance between residues Q69 and N144 (illustrated in e), exhibits minimal drift, indicating the preservation of the channel-like geometry during the simulation. f, The total number of hydrogen bonds, serving as an indirect indicator of intramolecular stability, shows no significant changes over the 2700–3000 ns simulation trajectories, which might indicate that the secondary and tertiary structures remain stable throughout the simulations. Collectively, these results demonstrate that all REE–MIF complexes converge to stable equilibrated states during the 3 µs MD simulations.

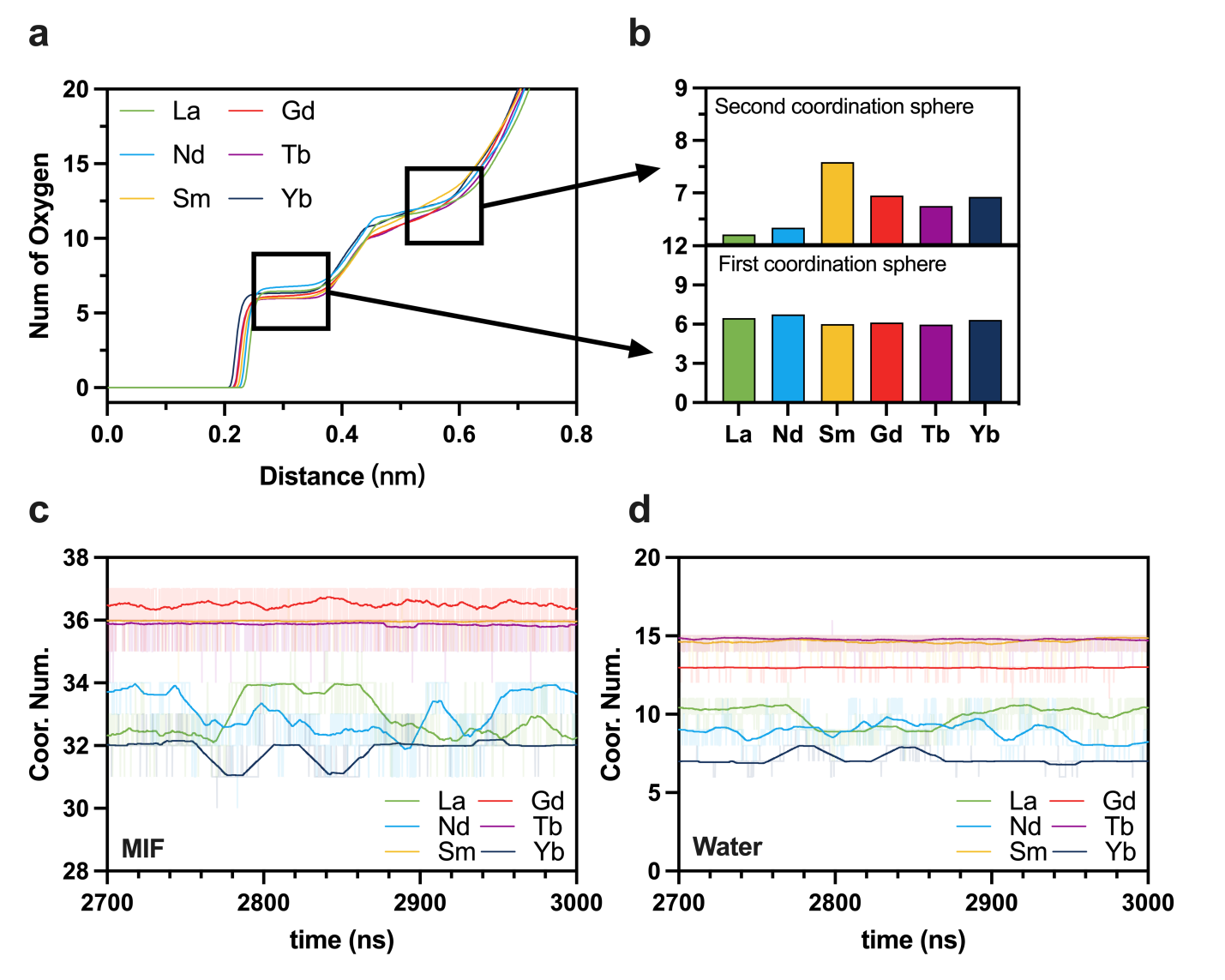

Supplementary Fig. S22. Coordination analysis of REE-MIF complexes from molecular dynamics trajectories
a, Radial distribution functions (RDFs) of MIF oxygen atoms around each REE ion were computed over the equilibrated MD trajectories (2700–3000 ns), averaged across all coordination sites. b, Average numbers of oxygen atoms within the first (≤0.3 nm) and second (0.3–0.6 nm) coordination spheres for each REE. Middle REEs exhibit higher second-sphere coordination numbers, indicating tighter packing of surrounding residues. c, Time evolution of the total coordination number derived from MIF residues over 2700–3000 ns, showing stable coordination microenvironments. d, Time evolution of the total coordination number derived from water molecules over 2700–3000 ns, also displaying stable behavior across trajectories. Together, the results suggest that both MIF- and water-mediated REE coordination microenvironments remain equilibrated and structurally stable throughout the simulations.

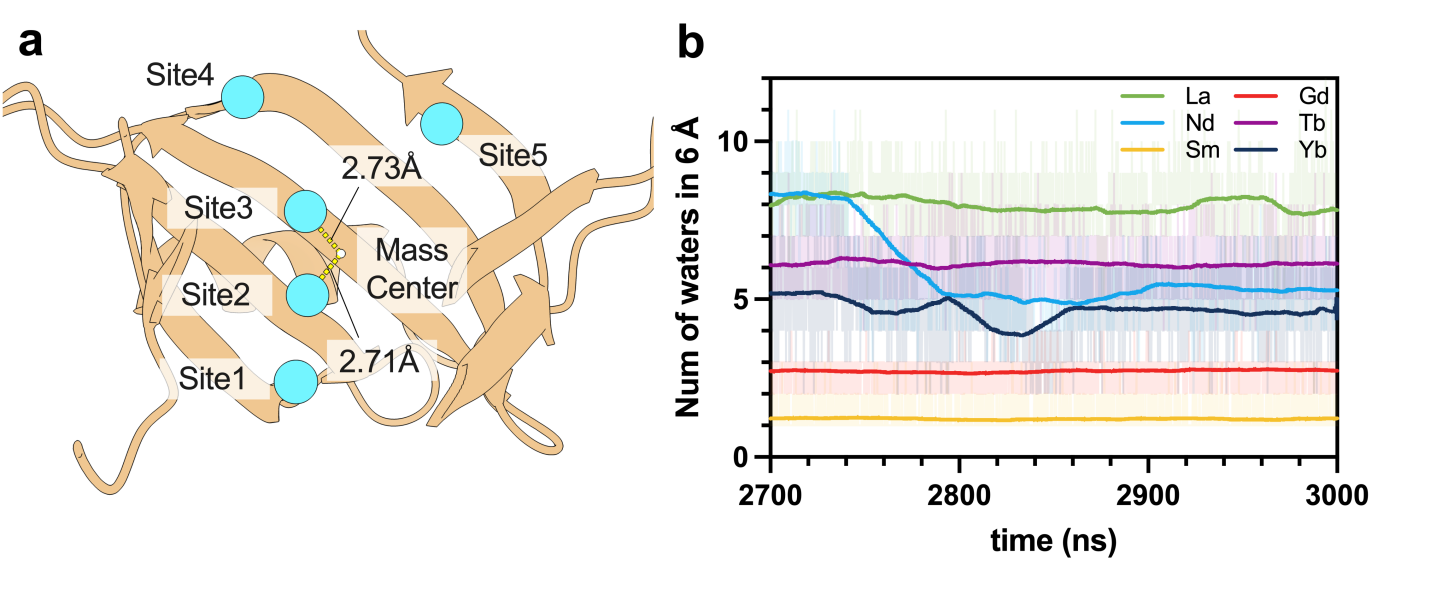

Supplementary Fig. S23. Water distribution near the central binding site of REE–MIF complexes
a, Structural illustration showing that Site 2 is the metal-binding site closest to the mass center of the MIF protein. Thus, a 6 Å sphere centered at Site 2 was selected for water molecule counting. b, Time-dependent number of water molecules within the 6 Å sphere over the last 300 ns (2700–3000 ns) for six REE–MIF complexes. All trajectories display stable water counts over time, indicating that water distributions around the coordination microenvironment is equilibrated.

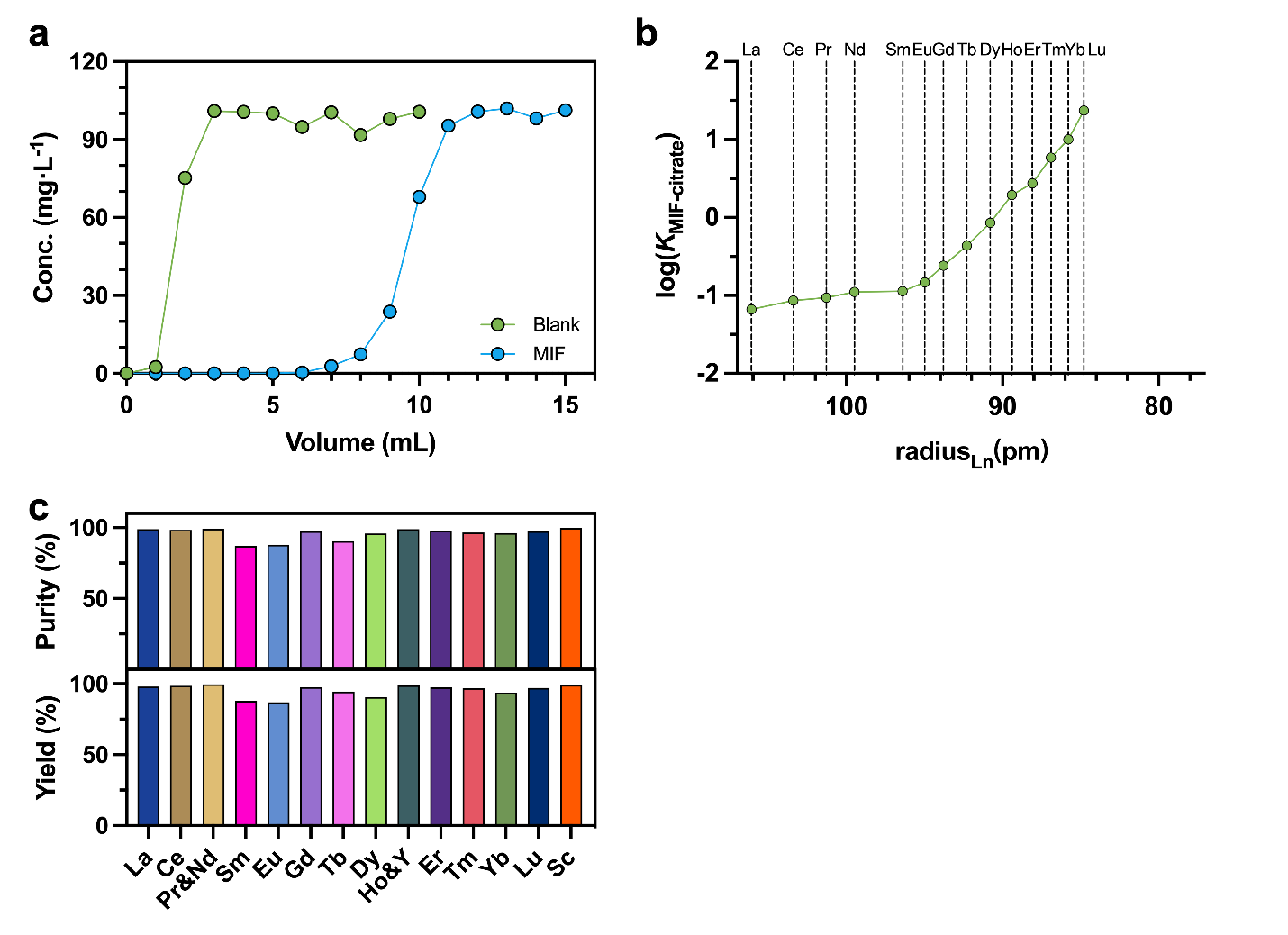

Supplementary Fig. S24. Properties of immobilized MIF and application in full-spectrum REE separation.
a, Immobilized MIF-agarose (MIF) and blank agarose (Blank) resins in a 1.2 mL column were used to evaluate column loading capacity. The immobilized His_10_-tagged MIF (molecular weight 18.63 kDa) loading was approximately 21 mg/mL. The adsorption of La^III^ on blank agarose was measured to be 1.75 µg/mL, while the La^III^ adsorption on agarose beads immobilized with MIF was 624.30 µg/mL. Thus, the net La^III^ binding capacity attributable to immobilized MIF was calculated to be 622.55 µg/mL. This corresponds to an average binding stoichiometry of approximately 3.98 La^III^ ions per MIF molecule. b, Estimated equilibrium constants for the conversion of REE-MIF to REE-citrate complexes. The result exhibited an increase of the equilibrium constant from La to Lu. The stability constant for citrate was taken from tabulated literature values (*34*). c, Purity (top panel, 87.0–99.2%) and yield (bottom panel, 87.1–99.6%) of each resolved peak from single-stage full-spectrum REE separation.

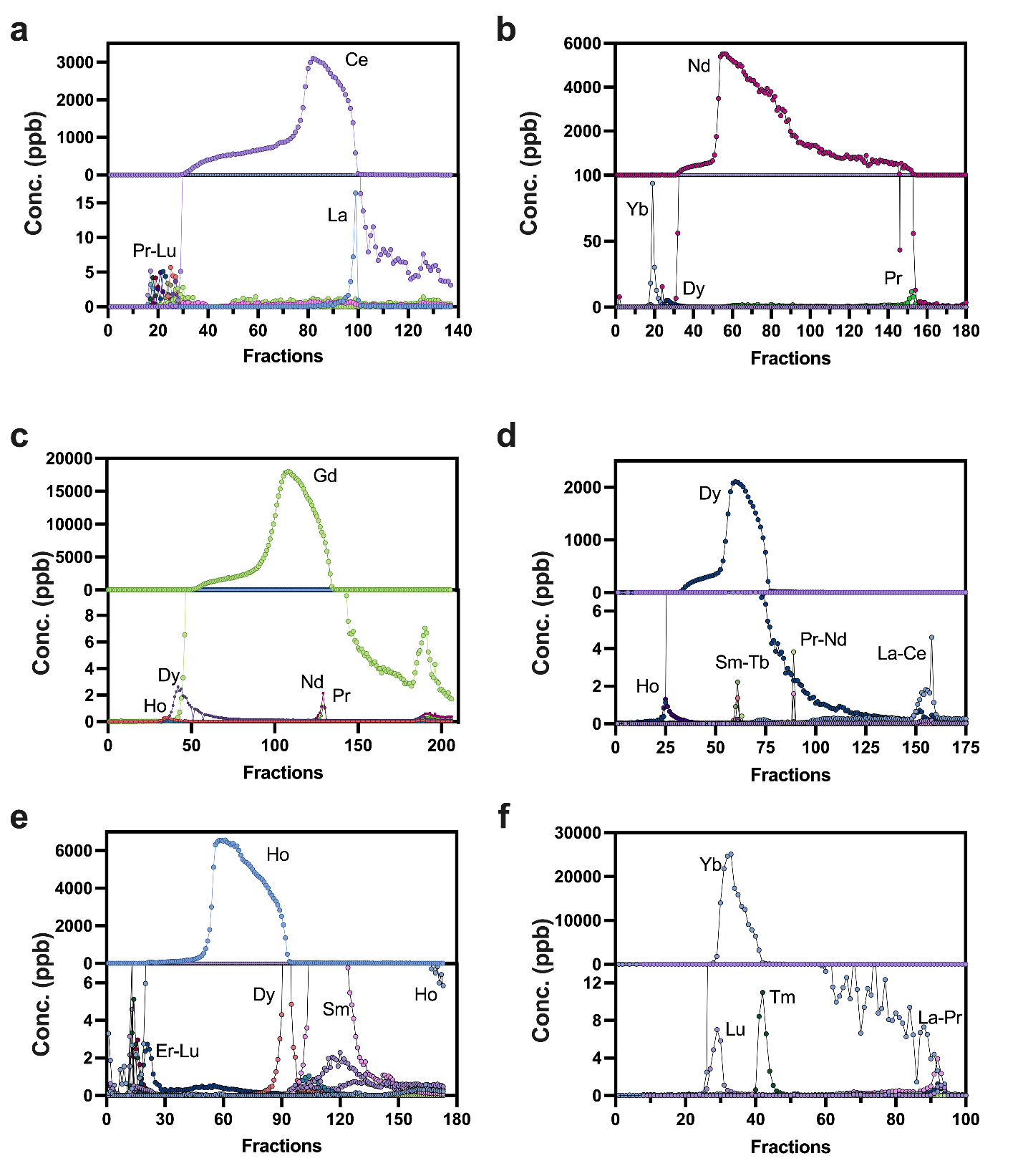

Supplementary Fig. S25. Ultra-high purity REE preparation from Commercial REE samples.
a, Preparation of Ce^III^ with purity of 99.997% from Commercial REE samples with purity of 99.65%. b, Purification of Nd^III^ from 99.84% to 99.996%. c, Purification of Gd^III^ from 99.98% to 99.999%. d, Purification of Dy^III^ from 99.76% to 99.997%. e, Purification of Ho^III^ from 99.66% to 99.996%. f, Purification of Yb^III^ from 99.93% to 99.999%. Using a single‑stage MIF chromatographic run, commercial REE samples were purified, and the preparation of REE with ultra-high purity was achieved.

Supplementary Table S1-S10

Supplementary Table S1. REE payload (RP) and Tm index (TI) of reported lanthanide binding proteins.

| Sequence Name | RP | TI | PDB ID | Ref. |
| --- | --- | --- | --- | --- |
| Mex-LanM | 0.239808153 | -0.851249507 | 6MI5 | (*35*) |
| Hans-LanM | 0.251889169 | -0.901977281 | 8DQ2 | (*36*) |
| **MIF sequence from lanpepsy** | **0.294637596** | **1.593952472** | - | (*37*) |
| XoxF Methylobacterium extorquens | 0.015403574 | 1.251596033 | 6OC5 | (*38*) |
| cod allergen Gad m 1.0201 | 0.174216028 | -0.026562531 | 9B26 | (*39*) |
| LanD-La | 0.073529412 | 1.635408837 | 9C8X | (*40*) |
| bovine alpha-lactalbumin | 0.070472163 | -2.579820673 | 6IP9 | (*41*) |
| XoxF from Methylomicrobium buryatense 5G | 0.014874312 | 0.812314308 | 6DAM | (*42*) |
| CaM | 0.059844405 | -3.518833406 | 2K0J | (*43*) |
| response regulator RegX3 from Mycobacterium tuberculosis | 0.079840319 | 0.038374802 | 2OQR | (*44*) |
| EF-hand Calcium binding Protein from Entamoeba Histolytica | 0.139082058 | 0.350057532 | 2I18 | (*45*) |
| CA LN CALBINDIN D9K | 0.111731844 | 1.236324334 | 1KSM | (*46*) |
| GSK3b in complex with CX-4945 | 0.024119633 | 1.118388663 | 7Z1F | (*47*) |
| Methanol dehydrogenase from Methylacidiphilum fumariolicum SolV | 0.01572327 | 0.417188699 | 4MAE | (*48*) |
| PedH from Pseudomonas putida KT2440 | 0.01525553 | 1.07112709 | 6ZCV | (*49*) |
| TIM barrel-ferredoxin fold fusion dimer | 0.026469031 | 1.455540224 | 6ZV9 | (*50*) |
| Nbe LBM | 0.067340067 | -0.144449074 | 6Y0E | (*51*) |
| cell-death related nuclease 4 (CRN-4) | 0.028121485 | 2.595741692 | 3CM6 | (*52*) |
| SPL-2 | 0.020169708 | 0.633669084 | - | (*53*) |
| DLanM | 0.200765586 | -0.150725583 | - | (*32*) |

Supplementary Table S2. Purities and yield of adjacent REE pairs separation.

| REE1-REE2 pairs | Purity | | Yield | | Concentration  (mg/L) | | Citrate Gradient  (mg/L) | |
| --- | --- | --- | --- | --- | --- | --- | --- | --- |
|  | REE1 | REE2 | REE1 | REE2 | REE1 | REE2 | Initial | Final |
| La-Ce | 97.1% | 99.0% | 98.7% | 96.9% | 16.28 | 18.17 | 300 | 420 |
| Ce-Pr | 98.8% | 98.6% | 98.3% | 98.6% | 19.00 | 17.94 | 270 | 360 |
| Pr-Nd | 86.6% | 85.7% | 86.2% | 85.4% | 10.90 | 10.43 | 240 | 300 |
| Nd-Sm | 92.1% | 96.5% | 93.0% | 92.9% | 19.15 | 28.91 | 228 | 282 |
| Sm-Eu | 91.9% | 88.2% | 87.4% | 92.0% | 18.84 | 19.04 | 210 | 264 |
| Eu-Gd | 99.1% | 99.7% | 99.5% | 99.0% | 18.21 | 19.31 | 180 | 240 |
| Gd-Tb | 97.2% | 98.2% | 97.3% | 94.6% | 19.98 | 19.76 | 168 | 216 |
| Tb-Dy | 95.6% | 97.1% | 90.6% | 90.7% | 18.77 | 19.80 | 144 | 204 |
| Dy-Ho | 99.8% | 99.8% | 99.6% | 99.1% | 19.94 | 20.06 | 138 | 186 |
| Ho-Er | 97.7% | 99.8% | 95.0% | 91.6% | 19.45 | 19.22 | 96 | 180 |
| Er-Tm | 99.2% | 99.3% | 97.7% | 98.2% | 20.19 | 20.17 | 90 | 138 |
| Tm-Yb | 96.2% | 98.4% | 98.1% | 95.9% | 17.99 | 18.36 | 84 | 108 |
| Yb-Lu | 96.7% | 97.9% | 97.6% | 96.7% | 11.09 | 12.17 | 75 | 96 |

Supplementary Table S3. Separation factors (SFs) of MIF for different adjacent lanthanide pairs.

| SF(La-Ce) | SF(Ce-Pr) | SF(Pr-Nd) | SF(Nd-Sm) | SF(Sm-Eu) | SF(Eu-Gd) | SF(Gd-Tb) |
| --- | --- | --- | --- | --- | --- | --- |
| 0.16 | 0.39 | 0.58 | 0.09 | 1.26 | 6.02 | 1.95 |
| SF(Tb-Dy) | SF(Dy-Ho) | SF(Ho-Er) | SF(Er-Tm) | SF(Tm-Yb) | SF(Yb-Lu) |  |
| 1.74 | 1.58 | 1.31 | 1.56 | 1.75 | 1.80 |  |

Supplementary Table S4. Calculated deformation energy (E_def_, kcal/mol).

|  | E (kcal/mol) | E_def_ (kcal/mol) |
| --- | --- | --- |
| La | -2041.6 | 59.8 |
| Nd | -2006.4 | 94.9 |
| Sm | -2068.2 | 33.1 |
| Gd | -2097.7 | 3.7 |
| Tb | -1804.4 | 297.0 |
| Yb | -1677.7 | 423.6 |
| Apo | -2101.3 | - |

Supplementary Table S5. Calculated binding energies (E_bind_, kcal/mol).

|  | E_AB_ (a.u.) | E_A_ (a.u.) | E_B_ (a.u.) | ΔE_bind_ (kcal/mol) |
| --- | --- | --- | --- | --- |
| La | -6924.92638 | -6765.88197 | -31.49866 | -973.3 |
| Nd | -6858.00740 | -6689.31938 | -33.34006 | -1247.3 |
| Sm | -7664.11330 | -7454.30571 | -34.55665 | -1548.5 |
| Gd | -7518.48789 | -7301.62604 | -35.78297 | -1358.0 |
| Tb | -7674.81328 | -7454.37342 | -36.32691 | -1555.2 |
| Yb | -6734.78646 | -6536.58038 | -39.23393 | -1277.9 |

Supplementary Table S6. Amino acid and DNA sequences of constructs used in this study.

| Construct | Protein or DNA sequence |
| --- | --- |
| Lanpepsy (full sequence with signal peptide underlined) | MVGKTLVVLSTFALAISSSLAIADHHFPKGKVSLETCLEAALKAKPGTVVKVEYKLEGETPVYEFDIESSDSTAWDVECDANTGKIVEIEQEVDSADHPLFKAKQKVSEAEARKTALAAHPGEIVEVEYEIEENGAASYEFDIKTKDGKEFKVEVDASTGKIVEANQEFYQIGKE |
| MIF sequence from lanpepsy | DHHFPKGKVSLETCLEAALKAKPGTVVKVEYKLEGETPVYEFDIESSDSTAWDVECDANTGKIVEIEQEVDSADHPLFKAKQKVSEAEARKTALAAHPGEIVEVEYEIEENGAASYEFDIKTKDGKEFKVEVDASTGKIVEANQEFYQIGKE |
| Untagged MIF sequence for this study | MDHHFPKGKVSLETCLEAALKAKPGTVVKVEYKLEGETPVYEFDIESSDSTAWDVECDANTGKIVEIEQEVDSADHPLFKAKQKVSEAEARKTALAAHPGEIVEVEYEIEENGAASYEFDIKTKDGKEFKVEVDASTGKIVEANQEFYQIGKE |
| His_10_-tagged MIF sequence for this study (His_10_-tag and SGGS linker underlined) | MHHHHHHHHHHSGGSDHHFPKGKVSLETCLEAALKAKPGTVVKVEYKLEGETPVYEFDIESSDSTAWDVECDANTGKIVEIEQEVDSADHPLFKAKQKVSEAEARKTALAAHPGEIVEVEYEIEENGAASYEFDIKTKDGKEFKVEVDASTGKIVEANQEFYQIGKE |
| Untagged MIF codon-optimized DNA sequence for this study (5'NdeI and 3'BlpI bolder, stop codon underlined) | **CATATG**GATCACCATTTTCCGAAAGGTAAAGTGAGCCTGGAAACCTGCCTGGAAGCGGCCCTGAAAGCCAAACCGGGCACCGTGGTGAAGGTGGAATATAAACTGGAAGGCGAAACCCCGGTGTACGAATTTGATATTGAAAGCAGCGATAGCACCGCGTGGGATGTGGAATGCGATGCGAACACCGGCAAAATTGTGGAAATTGAACAGGAAGTGGATAGCGCGGATCACCCGCTGTTTAAAGCGAAACAGAAAGTGAGCGAAGCCGAAGCGCGCAAAACCGCCCTGGCGGCGCACCCGGGCGAAATTGTGGAAGTGGAATATGAAATTGAGGAAAATGGCGCAGCCAGCTACGAATTTGACATCAAAACCAAAGACGGCAAAGAATTTAAAGTGGAAGTGGATGCCAGCACCGGTAAAATTGTGGAAGCGAACCAGGAATTTTATCAGATTGGCAAAGAATAACTCGAGATCAAACGGGCTAGCTGAGATCCGGCTGCTAACAAAGCCCGAAAGGAAGCTGAGTTGGCTGCTGCCACC**GCTGAGC** |
| His_10_-tagged MIF codon-optimized DNA sequence for this study (5'NdeI and 3'XhoI bolder, His_10_-tag and stop codon underlined) | **CATATG**CATCATCATCATCACCACCATCATCATCATTCGGGCGGCAGCGATCACCATTTTCCGAAAGGTAAAGTGAGCCTGGAAACCTGCCTGGAAGCGGCCCTGAAAGCCAAACCGGGCACCGTGGTGAAGGTGGAATATAAACTGGAAGGCGAAACCCCGGTGTACGAATTTGATATTGAAAGCAGCGATAGCACCGCGTGGGATGTGGAATGCGATGCGAACACCGGCAAAATTGTGGAAATTGAACAGGAAGTGGATAGCGCGGATCACCCGCTGTTTAAAGCGAAACAGAAAGTGAGCGAAGCCGAAGCGCGCAAAACCGCCCTGGCGGCGCACCCGGGCGAAATTGTGGAAGTGGAATATGAAATTGAGGAAAATGGCGCAGCCAGCTACGAATTTGACATCAAAACCAAAGACGGCAAAGAATTTAAAGTGGAAGTGGATGCCAGCACCGGTAAAATTGTGGAAGCGAACCAGGAATTTTATCAGATTGGCAAAGAATAA**CTCGAG** |

Supplementary Table S7. Test parameters for Isothermal Titration Calorimetry

| REEs | MIF concentration (µM) | REE concentration (mM) | Number of total injections | Volume per injection (µL) | Interval  (s) |
| --- | --- | --- | --- | --- | --- |
| La | 31.2 | 9.25 | 20 | 0.5 | 120 |
| Ce | 78.9 | 10.20 | 25 | 0.5 | 150 |
| Pr | 78.9 | 8.35 | 28 | 0.5 | 120 |
| Nd | 31.2 | 7.90 | 20 | 0.5 | 120 |
| Sm | 78.9 | 9.51 | 21 | 1.0 | 120 |
| Eu | 78.9 | 8.80 | 28 | 0.5 | 120 |
| Gd | 78.9 | 8.90 | 28 | 0.5 | 120 |
| Tb | 78.9 | 12.1 | 28 | 0.5 | 120 |
| Dy | 54.9 | 11.9 | 28 | 0.5 | 120 |
| Ho | 65.8 | 9.67 | 28 | 0.5 | 120 |
| Er | 78.9 | 13.60 | 28 | 0.5 | 120 |
| Tm | 78.9 | 9.55 | 28 | 0.5 | 120 |
| Yb | 78.9 | 8.90 | 28 | 0.5 | 120 |
| Lu | 78.9 | 9.67 | 25 | 0.5 | 150 |

Supplementary Table S8. Crystallographic data collection and refinement statistics.

|  | **Apo MIF^†^**  **(PDB 9VY3)** | **MIF-La(III) ^†^ (PDB 9VY7)** | **MIF-Nd(III) ^†^ (PDB 9VYA)** | **MIF-Sm(III) ^†^ (PDB 9VY8)** | **MIF-Gd(III) ^†^ (PDB 9VY9)** | **MIF-Tb(III) ^†^ (PDB 9VY5)** | **MIF-Ho(III) ^†^ (PDB 9VY6)** | **MIF-Yb(III) ^†^ (PDB 9VY4)** |
| --- | --- | --- | --- | --- | --- | --- | --- | --- |
| **Data collection** |  |  |  |  |  |  |  |  |
| Space group | P 2_1_ 2_1_ 2_1_ | C 1 2 1 | C 1 2 1 | I 4_1_ 2 2 | P 2_1_ 2_1_ 2_1_ | P 2_1_ 2_1_ 2 | C 1 2 1 | P 2_1_ 2_1_ 2 |
| Cell dimensions |  |  |  |  |  |  |  |  |
| *a*, *b*, *c* (Å) | 70.748, 70.817, 130.534 | 70.896, 40.741, 56.416 | 71.540, 40.560, 56.020 | 119.052, 119.052,79.209 | 57.804,101.146, 115.403 | 71.734,119.988, 36.28 | 200.033,64.491, 92.945 | 180.205,67.762, 81.967 |
| *α, β, γ* (°) | 90, 90, 90 | 90, 112.519, 90 | 90, 112.530, 90 | 90, 90, 90 | 90, 90, 90 | 90, 90, 90 | 90, 114.644, 90 | 90, 112.519, 90 |
| Resolution range (Å) | 39.72 - 2.08 (2.15 - 2.08)^a^ | 52.11 - 1.19 (1.26 - 1.19)^a^ | 51.74 - 1.41 (1.45 - 1.41)^a^ | 84.18 - 2.16 (2.24 - 2.16)^a^ | 76.07 - 1.41 (1.49 - 1.41)^a^ | 36.28 - 1.55 (1.59 - 1.55)^a^ | 90.91 - 2.97 (3.22 - 2.97)^a^ | 61.92 - 1.21 (1.28 - 1.21)^a^ |
| No. of unique reflections | 40085 (3939)^a^ | 39396 (2135)^a^ | 28608 (2101)^a^ | 15560 (1508)^a^ | 129245(18565)^a^ | 46026 (3027)^a^ | 16753 (838)^a^ | 73230 (2751)^a^ |
| *R*_merge_ | 0.168 | 0.073 | 0.131 | 0.120 | 0.177 | 0.204 | 0.190 | 0.097 |
| *R*_pim_ | 0.050 | 0.035 | 0.059 | 0.032 | 0.059 | 0.062 | 0.097 | 0.030 |
| *R*_meas_ | 0.176 | 0.081 | 0.145 | 0.124 | 0.187 | 0.214 | 0.214 | 0.102 |
| *I/*σ(*I*) | 9.9 (4.2)^a^ | 16.6 (2.9)^a^ | 5.6 (3.6)^a^ | 16.8(2.4)^a^ | 8.7 (1.2)^a^ | 7.3 (2.1)^a^ | 5.6 (1.6)^a^ | 21.9 (1.3)^a^ |
| *CC*_1/2_ | 0.958 (0.860)^a^ | 0.996 (0.787)^a^ | 0.995 (0.617)^a^ | 0.999 (0.903)^a^ | 0.995 (0.421)^a^ | 0.989 (0.860)^a^ | 0.919 (0.516)^a^ | 0.996 (0.787)^a^ |
| Completeness (%) | 99.7 (99.8)^a^ | 82.4 (30.8)^a^ | 99.5 (99.3)^a^ | 99.5 (98.3)^a^ | 99.7 (99.2)^a^ | 99.0 (89.4)^a^ | 91.2 (45.3)^a^ | 76.8 (20.4)^a^ |
| Redundancy | 12.5 (11.6)^a^ | 4.6 (1.5)^a^ | 5.6 (3.6)^a^ | 14.1 (12.4)^a^ | 9.3 (7.3)^a^ | 11.2 (6.6)^a^ | 5.0 (4.4)^a^ | 9.2 (1.2)^a^ |
| **Refinement** |  |  |  |  |  |  |  |  |
| Resolution (Å) | 37.03 - 2.08 (2.15 - 2.08)^b^ | 26.06 - 1.25 (1.30 - 1.25)^b^ | 26.40 - 1.41 (1.46 - 1.41)^b^ | 37.65 - 2.16 (2.24 – 2.16)^b^ | 36.15 - 1.41 (1.47 - 1.41)^b^ | 34.94 - 1.58 (1.64 - 1.58)^b^ | 80.97 - 3.05 (3.16 - 3.05)^b^ | 35.06 - 1.21 (1.26 - 1.21)^b^ |
| No. reflections | 39984 (3930)^b^ | 37494 (2094)^b^ | 28598 (2837)^b^ | 15487 (1499)^b^ | 129073(12650)^b^ | 43478 (4140)^b^ | 16680 (383)^b^ | 73134 (1343)^b^ |
| *R*_work_ (%) | 22.02 (28.17)^b^ | 12.08 (17.30)^b^ | 14.22 (24.22)^b^ | 23.10 (39.76)^b^ | 15.01 (28.52)^b^ | 18.01 (24.84)^b^ | 21.93 (33.91)^b^ | 12.44 (26.04)^b^ |
| *R*_free_ (%) | 24.88 (29.24)^b^ | 12.90 (17.31)^b^ | 15.88 (22.86)^b^ | 23.58 (42.35)^b^ | 16.61 (29.75)^b^ | 20.32 (27.13)^b^ | 26.79 (36.41)^b^ | 13.78 (30.69)^b^ |
| No. of non-hydrogen atoms | 5060 | 1420 | 1359 | 1191 | 5453 | 2886 | 6751 | 2836 |
| Protein | 4697 | 1178 | 1168 | 1168 | 4690 | 2336 | 6709 | 2336 |
| Ligand/ion | 0 | 5 | 5 | 6 | 30 | 13 | 42 | 13 |
| Water | 363 | 237 | 186 | 17 | 733 | 537 | 0 | 487 |
| *B* factors (Å^2^) | 35.77 | 15.14 | 13.68 | 62.58 | 24.03 | 22.07 | 105.97 | 17.03 |
| Protein | 35.70 | 13.66 | 11.85 | 62.69 | 23.02 | 20.61 | 106.11 | 14.85 |
| Ligand | / | 7.51 | 5.90 | 55.98 | 20.89 | 14.39 | 83.09 | 11.09 |
| Solvent | 36.63 | 22.68 | 25.40 | 56.69 | 30.65 | 28.60 | / | 27.64 |
| Ramachandran |  |  |  |  |  |  |  |  |
| Favored (%) | 98.99 | 98.66 | 98.65 | 97.30 | 99.16 | 99.66 | 98.58 | 100.00 |
| Allowed (%) | 1.01 | 1.34 | 1.35 | 2.70 | 0.84 | 0.34 | 1.42 | 0.00 |
| Outliers (%) | 0.00 | 0.00 | 0.00 | 0.00 | 0.00 | 0.00 | 0.00 | 0.00 |
| R.m.s. deviations |  |  |  |  |  |  |  |  |
| Bond lengths (Å) | 0.007 | 0.016 | 0.016 | 0.014 | 0.015 | 0.008 | 0.003 | 0.019 |
| Bond angles (°) | 0.73 | 1.38 | 1.36 | 1.58 | 1.49 | 0.88 | 0.61 | 1.53 |
| Clashscore | 4.13 | 2.59 | 1.74 | 7.83 | 2.71 | 4.57 | 8.10 | 1.74 |
| ^a^ Values in parentheses are for the highest-resolution shell (outer shell) in data collection.  ^b^ Values in parentheses are for the highest-resolution shell (outer shell) in refinement, the reflection number and resolution may be different from data collection due to the resolution-range cut off in refinement.  ^†^The data are collected from one crystal. | | | | | | | | |

Supplementary Table S9. REE concentrations in the loading solution for full spectrum separation in Fig. 4a. Concentrations were determined by ICP-OES analysis.

| La (mg/L) | Ce (mg/L) | Pr (mg/L) | Nd (mg/L) | Sm (mg/L) | Eu (mg/L) | Gd (mg/L) | Tb (mg/L) |
| --- | --- | --- | --- | --- | --- | --- | --- |
| 10.29 | 10.29 | 10.30 | 10.31 | 10.27 | 10.30 | 10.27 | 10.29 |
| Dy (mg/L) | Ho (mg/L) | Er (mg/L) | Tm (mg/L) | Yb (mg/L) | Lu (mg/L) | Sc (mg/L) | Y (mg/L) |
| 10.30 | 10.38 | 10.29 | 10.27 | 10.21 | 10.21 | 7.24 | 8.43 |

Supplementary Table S10. Amounts and purities of starting materials, and citrate concentration of gradient elution for ultra-high purity REE preparation.

| REEs | Purity of starting  materials (%) | Loading amount (mg) | Concentration of citrate for impurity wash (mg/L) | Concentration of  citrate gradient (mg/L) | |
| --- | --- | --- | --- | --- | --- |
|  |  |  |  | Initial | Final |
| Ce | 99.65 | 1.67 | 270 | 270 | 330 |
| Nd | 99.84 | 4.40 | 252 | 252 | 270 |
| Gd | 99.98 | 11.60 | 192 | 192 | 216 |
| Dy | 99.76 | 0.67 | 156 | 156 | 186 |
| Ho | 99.66 | 4.01 | 138 | 138 | 156 |
| Yb | 99.93 | 3.50 | 84 | 84 | 102 |

**Reference**

1.	G. Böhm, R. Muhr, R. Jaenicke, Quantitative analysis of protein far UV circular dichroism spectra by neural networks. *Protein Engineering, Design and Selection* **5**, 191-195 (1992).

2.	A. Brown, Analysis of cooperativity by isothermal titration calorimetry. *International journal of molecular sciences* **10**, 3457-3477 (2009).

3.	P. R. Evans, G. N. Murshudov, How good are my data and what is the resolution? *Biological crystallography* **69**, 1204-1214 (2013).

4.	Z. Otwinowski, W. Minor, in *Methods in enzymology*. (Elsevier, 1997), vol. 276, pp. 307-326.

5.	D. Liebschner *et al.*, Macromolecular structure determination using X-rays, neutrons and electrons: recent developments in Phenix. *Biological Crystallography* **75**, 861-877 (2019).

6.	J. Abramson *et al.*, Accurate structure prediction of biomolecular interactions with AlphaFold 3. *Nature* **630**, 493-500 (2024).

7.	P. Emsley, B. Lohkamp, W. G. Scott, K. Cowtan, Features and development of Coot. *Biological crystallography* **66**, 486-501 (2010).

8.	J. M. Walker, *The proteomics protocols handbook*. (Springer, 2005).

9.	T. Ku *et al.*, Predicting melting temperature directly from protein sequences. *Computational Biology and Chemistry* **33**, 445-450 (2009).

10.	F. Teufel *et al.*, SignalP 6.0 predicts all five types of signal peptides using protein language models. *Nature Biotechnology* **40**, 1023-1025 (2022).

11.	B. Hess, C. Kutzner, D. Van Der Spoel, E. Lindahl, GROMACS 4: algorithms for highly efficient, load-balanced, and scalable molecular simulation. *Journal of chemical theory and computation* **4**, 435-447 (2008).

12.	S. Páll *et al.*, Heterogeneous parallelization and acceleration of molecular dynamics simulations in GROMACS. *The Journal of chemical physics* **153**, (2020).

13.	R. B. Best *et al.*, Optimization of the additive CHARMM all-atom protein force field targeting improved sampling of the backbone ϕ, ψ and side-chain χ1 and χ2 dihedral angles. *Journal of chemical theory and computation* **8**, 3257-3273 (2012).

14.	V. Migliorati, A. Serva, F. M. Terenzio, P. D’Angelo, Development of Lennard-Jones and Buckingham potentials for lanthanoid ions in water. *Inorganic chemistry* **56**, 6214-6224 (2017).

15.	U. Essmann *et al.*, A smooth particle mesh Ewald method. *The Journal of chemical physics* **103**, 8577-8593 (1995).

16.	H. C. Andersen, Rattle: A “velocity” version of the shake algorithm for molecular dynamics calculations. *Journal of computational Physics* **52**, 24-34 (1983).

17.	G. Bussi, D. Donadio, M. Parrinello, Canonical sampling through velocity rescaling. *The Journal of chemical physics* **126**, (2007).

18.	M. Parrinello, A. Rahman, Polymorphic transitions in single crystals: A new molecular dynamics method. *Journal of Applied physics* **52**, 7182-7190 (1981).

19.	J. Řezáč, P. Hobza, A halogen-bonding correction for the semiempirical PM6 method. *Chemical Physics Letters* **506**, 286-289 (2011).

20.	J. Rezac, P. Hobza, Advanced corrections of hydrogen bonding and dispersion for semiempirical quantum mechanical methods. *Journal of Chemical Theory and Computation* **8**, 141-151 (2012).

21.	J. J. Stewart, Application of localized molecular orbitals to the solution of semiempirical self‐consistent field equations. *International journal of quantum chemistry* **58**, 133-146 (1996).

22.	Q. Tan, Y. Ding, Z. Qiu, J. Huang, Binding Energy and Free Energy of Calcium Ion to Calmodulin EF-Hands with the Drude Polarizable Force Field. *ACS Physical Chemistry Au* **2**, 143-155 (2022).

23.	F. Himo, Recent trends in quantum chemical modeling of enzymatic reactions. *Journal of the American Chemical Society* **139**, 6780-6786 (2017).

24.	C. Adamo, V. Barone, Toward reliable density functional methods without adjustable parameters: The PBE0 model. *The Journal of chemical physics* **110**, 6158-6170 (1999).

25.	S. Grimme, J. Antony, S. Ehrlich, H. Krieg, A consistent and accurate ab initio parametrization of density functional dispersion correction (DFT-D) for the 94 elements H-Pu. *The Journal of chemical physics* **132**, (2010).

26.	S. Grimme, S. Ehrlich, L. Goerigk, Effect of the damping function in dispersion corrected density functional theory. *Journal of computational chemistry* **32**, 1456-1465 (2011).

27.	E. Cances, B. Mennucci, J. Tomasi, A new integral equation formalism for the polarizable continuum model: Theoretical background and applications to isotropic and anisotropic dielectrics. *The Journal of chemical physics* **107**, 3032-3041 (1997).

28.	J. S. Binkley, J. A. Pople, W. J. Hehre, Self-consistent molecular orbital methods. 21. Small split-valence basis sets for first-row elements. *Journal of the American Chemical Society* **102**, 939-947 (1980).

29.	F. Weigend, R. Ahlrichs, Balanced basis sets of split valence, triple zeta valence and quadruple zeta valence quality for H to Rn: Design and assessment of accuracy. *Physical Chemistry Chemical Physics* **7**, 3297-3305 (2005).

30.	A. V. Marenich, C. J. Cramer, D. G. Truhlar, Universal solvation model based on solute electron density and on a continuum model of the solvent defined by the bulk dielectric constant and atomic surface tensions. *The Journal of Physical Chemistry B* **113**, 6378-6396 (2009).

31.	Z. Dong, J. A. Mattocks, J. A. Seidel, J. A. Cotruvo, D. M. Park, Protein-based approach for high-purity Sc, Y, and grouped lanthanide separation. *Separation and Purification Technology* **333**, 125919 (2024).

32.	H. Cui *et al.*, The Construction of a Microbial Synthesis System for Rare Earth Enrichment and Material Applications. *Advanced Materials* **35**, 2303457 (2023).

33.	E. F. Pettersen *et al.*, UCSF ChimeraX: Structure visualization for researchers, educators, and developers. *Protein science* **30**, 70-82 (2021).

34.	R. H. Byrne, B. Li, Comparative complexation behavior of the rare earths. *Geochimica et Cosmochimica Acta* **59**, 4575-4589 (1995).

35.	E. C. Cook, E. R. Featherston, S. A. Showalter, J. A. Cotruvo Jr, Structural basis for rare earth element recognition by Methylobacterium extorquens lanmodulin. *Biochemistry* **58**, 120-125 (2018).

36.	J. A. Mattocks *et al.*, Enhanced rare-earth separation with a metal-sensitive lanmodulin dimer. *Nature* **618**, 87-93 (2023).

37.	J. L. Hemmann *et al.*, Lanpepsy is a novel lanthanide-binding protein involved in the lanthanide response of the obligate methylotroph Methylobacillus flagellatus. *Journal of Biological Chemistry* **299**, (2023).

38.	N. M. Good *et al.*, Lanthanide-dependent alcohol dehydrogenases require an essential aspartate residue for metal coordination and enzymatic function. *Journal of Biological Chemistry* **295**, 8272-8284 (2020).

39.	A. O'Malley *et al.*, Comparative studies of seafood and reptile α‐and β‐parvalbumins. *Protein Science* **33**, e5226 (2024).

40.	W. B. Larrinaga, J. J. Jung, C.-Y. Lin, A. K. Boal, J. A. Cotruvo, Modulating metal-centered dimerization of a lanthanide chaperone protein for separation of light lanthanides. *Proceedings of the National Academy of Sciences* **121**, e2410926121 (2024).

41.	D. S. Yarramala *et al.*, Cytotoxicity of apo bovine α-lactalbumin complexed with La3+ on cancer cells supported by its high resolution crystal structure. *Scientific Reports* **9**, 1780 (2019).

42.	Y. W. Deng, S. Y. Ro, A. C. Rosenzweig, Structure and function of the lanthanide-dependent methanol dehydrogenase XoxF from the methanotroph Methylomicrobium buryatense 5GB1C. *JBIC Journal of Biological Inorganic Chemistry* **23**, 1037-1047 (2018).

43.	I. Bertini *et al.*, Accurate solution structures of proteins from X-ray data and a minimal set of NMR data: calmodulin− peptide complexes as examples. *Journal of the American Chemical Society* **131**, 5134-5144 (2009).

44.	J. King-Scott *et al.*, The structure of a full-length response regulator from Mycobacterium tuberculosis in a stabilized three-dimensional domain-swapped, activated state. *Journal of Biological Chemistry* **282**, 37717-37729 (2007).

45.	S. M. Mustafi, S. Mukherjee, K. V. Chary, G. Cavallaro, Structural basis for the observed differential magnetic anisotropic tensorial values in calcium binding proteins. *Proteins: Structure, Function, and Bioinformatics* **65**, 656-669 (2006).

46.	I. Bertini *et al.*, Paramagnetism-based versus classical constraints: an analysis of the solution structure of Ca Ln calbindin D9k. *Journal of biomolecular NMR* **21**, 85-98 (2001).

47.	P. Grygier, K. Pustelny, G. Dubin, A. Czarna, Crystal structure of GSK3b in complex with CX-4945. *Worldwide Protein Data Bank*, (2023).

48.	A. Pol *et al.*, Rare earth metals are essential for methanotrophic life in volcanic mudpots. *Environmental microbiology* **16**, 255-264 (2014).

49.	M. Wehrmann *et al.*, Engineered PQQ-dependent alcohol dehydrogenase for the oxidation of 5-(hydroxymethyl) furoic acid. *ACS catalysis* **10**, 7836-7842 (2020).

50.	S. J. Caldwell *et al.*, Tight and specific lanthanide binding in a de novo TIM barrel with a large internal cavity designed by symmetric domain fusion. *Proceedings of the National Academy of Sciences* **117**, 30362-30369 (2020).

51.	G. Pompidor, S. Zimmermann, C. Loew, T. Schneider, in *Worldwide Protein Data Bank*. (2021).

52.	Y.-Y. Hsiao *et al.*, Crystal structure of CRN-4: implications for domain function in apoptotic DNA degradation. *Molecular and cellular biology* **29**, 448-457 (2009).

53.	M. Tracz, I. Górniak, A. Szczepaniak, W. Białek, E3 ubiquitin ligase SPL2 is a lanthanide-binding protein. *International Journal of Molecular Sciences* **22**, 5712 (2021).
